# Comparison of analytical methods for rapid & reliable quantification of plant-based carbohydrates for the quintessential bioenergy educator

**DOI:** 10.1101/2020.05.21.106468

**Authors:** Shishir P. S. Chundawat

**Affiliations:** Department of Chemical and Biochemical Engineering, Rutgers The State University of New Jersey, Busch Campus, 98 Brett Road, Engineering Building Wing C, Room C150A, Piscataway, NJ 08854

**Keywords:** *Audience-specific* (General public, High school, First-year undergraduate), *domain-specific* (Analytical chemistry, chemical engineering, interdisciplinary, laboratory instruction, polymer chemistry), *pedagogy-specific* (Collaborative learning, hands-on learning, inquiry-based learning, problem solving), *topic-specific* (Agricultural chemistry, bioanalytical chemistry, calibration, carbohydrates, enzymes, green chemistry, kinetics, plant chemistry, quantitative analysis, spectroscopy, undergraduate research)

## Abstract

Biochemical conversion of plant-based insoluble carbohydrate polymers, such as starch from corn grains or cellulose-hemicellulose from corn stover, into soluble fermentable sugars (e.g., glucose and xylose) for bioenergy production has seen tremendous research activity and commercial-scale biorefineries deployment over the last three decades, particularly in regions around the world that have a dominant agricultural-based economy. Therefore, educators in schools and universities have developed various hands-on experimental activities to engage the general public and students either in outreach events or lab/classroom-based settings to instruct students on various inter-disciplinary concepts relevant to bioenergy and biochemicals production. One of the limitations of most available protocols is the lack of systematic and comprehensive comparison of educator-friendly analytical tools and protocols for quantitative analysis of water-soluble carbohydrates commonly encountered in a biorefinery backdrop during the biochemical conversion of lignocellulosic biomass to biofuels/biochemicals. Here, we systematically compare and validate findings from four leading analytical approaches for detection and quantification of lignocellulosic biomass derived soluble carbohydrates. We compare these assay methods based on the overall ease of use, detection accuracy/sensitivity, commercial availability, analytical cost per assay run, and suitability for use by instructors in biorefining specific hands-on activity protocols. Next, we provide a detailed instructional protocol that utilizes one of these validated soluble sugar assays as part of a ∼90 min hands-on bioenergy focused activity (called ‘Grass-to-Gas!’) conducted at Rutgers University with pre-university high school students. ‘Grass-to-Gas!’ activity involves students running biochemical assays that helps them understand the various facets of cellulosic biomass hydrolysis by commercial cellulase enzymes and monitoring the total glucose product released using our validated sugar assays to finally estimate the fractional conversion of cellulose-to-glucose. Lastly, we further demonstrate how such carbohydrate-based analytical methods can be used by instructors to help university students explore and understand various chemistry, biochemistry, and chemical engineering concepts relevant to other advanced operations involved in lignocellulose biorefining. These activity protocols would greatly aid educators teaching interdisciplinary science and engineering concepts to students in the backdrop of lignocellulose biorefining.

## Introduction

Deconstruction of complex plant biomass ultimately into carbon dioxide by microbial communities is a critical process in the global cycling of terrestrial organic carbon. Studying these processes also has practical relevance for converting inedible biomass into desired biofuels or bioproducts and decreasing our overall dependence on fossil fuels.^1^ Most agricultural plants (e.g., maize, sugarcane) grown worldwide generate a significant quantity of waste cellulosic biomass (e.g., corn stalks, sugarcane bagasse) after separation of the desired food-products (e.g., cereals, grains, sucrose). Biochemical processes are promising routes that utilize chemical catalysts, enzymes, and/or microbes to upgrade biomass-derived carbohydrate polymers from either waste or food-grade biomass to fuels and chemicals (**Figure 1**).^2^ In the last decade there has been interest in educating students about STEM concepts, both in high-schools and universities, in the context of biofuels production from renewable biomass enriched in either lipids (e.g., algae, oilseeds) or carbohydrates (e.g., grains, grass, wood).^3,4^ The Next Generation Science Standards and the framework for school science education has identified seven interdisciplinary concepts to help learners develop their understanding of these complex issues.^5^ One of these core concepts is the ability to trace material and energy flows through systems of varying scales. Several researchers have thus developed bioenergy education focused hands-on laboratory protocols for the analysis of conversion processes and production of biomass-derived intermediates (e.g., lipids, sugars) or final bioproducts (e.g., biodiesel, bioethanol).^3,4,6–13^ However, despite the development of significant bioenergy teaching curriculum,^7,11,14,15^ there has been limited emphasis to systematically trace material balances around bioenergy-relevant processes such as cellulosic biofuels production in a biorefinery backdrop. One of the issues identified has been the lack of inexpensive, validated analytical methods for the rapid analysis of carbohydrates derived from native plant biomass and associated biorefining processes for the production of either soluble sugars or ultimately biofuels/biochemicals from diverse lignocellulosic feedstocks.

**Figure 1.**
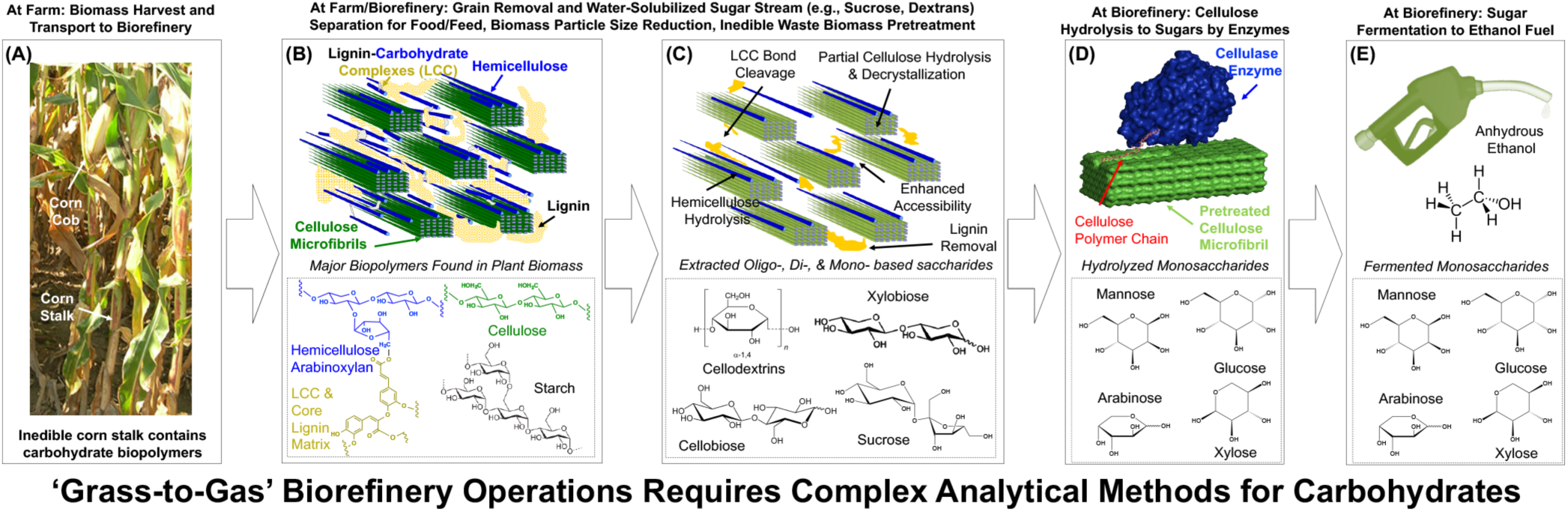
Overview of complex carbohydrate structures found within cellulosic biomass and relevant carbohydrate intermediates associated with biorefinery operations necessary for conversion of biomass-to-biofuels (i.e., ‘Grass-to-Gas!’). Here, we use maize derived cellulosic biomass as an example feedstock for a next-generation ethanol biorefinery. Cellulosic biomass is enriched in complex carbohydrate-based polymers (e.g., sucrose, starch, cellulose, and hemicellulose) that can be converted in a biorefinery using multiple processing steps (i.e., thermochemical pretreatment, enzymatic hydrolysis, and fermentation). This highlights the critical need for analytical assays that enable plant-based carbohydrates quantification for developing advanced laboratory experiments or simple hands-on outreach activities for student instruction.

In a classical corn-based biorefinery, the mature corn grains harvested from the cobs are converted into ethanol using enzymes/microbes, but the inedible biomass fraction (i.e., corn stalks or stover) is not utilized. However, in next-generation corn-based biorefineries that are currently being commercialized, the waste agricultural residues composed of mostly inedible fractions of the corn plant (e.g., stalk, cobs, leaves) are also transported to a biorefinery for biochemical processing as highlighted in **Figure 1**. Waste lignocellulosic biomass like corn stalks are mostly enriched in complex carbohydrate-polymers that are tightly embedded within plant cell walls in an amorphous matrix of lignin-carbohydrates based biopolymers.^16^ Lignocellulosic biomass feedstock is a highly heterogeneous biomaterial composed of three major biopolymers, namely, cellulose (a glucose based semi-crystalline and homogeneous polymer), hemicellulose (a complex pentose/hexose sugars based highly branched and heterogeneous polymer), and lignin (a phenyl-propanoid monomer based complex aromatic polymer).^2^ The major carbohydrate biopolymers found in waste cellulosic biomass like corn stalks includes cellulose (e.g., *β*-1,4-linear glucose polymer), hemicellulose (e.g., *β*-1,4-xylose polymer with arabinose branched side-chains), and starch (e.g., *α*-1,4/6-branched glucose polymer). However, in addition to these insoluble polysaccharides, plant biomass is also enriched in a variety of water-soluble carbohydrates like sucrose and cellodextrins. Biomass from municipal solid waste (e.g., cardboard), woody dicots (e.g., poplar), monocot grasses (e.g., corn stover, switchgrass), and woody gymnosperms (e.g., Douglas Fir, Aspen) are all currently being targeted as candidate feedstocks for biorefinery processing. But these diverse feedstocks have considerable variation in their carbohydrate composition that would impact biorefinery processes.^17,18^ Once the whole corn plant biomass is received at next-generation biorefineries, unit operations like particle size reduction and water-washing are often conducted to remove any soluble sugars (e.g., sucrose) prior to separate thermochemical processing of the lignocellulosic biomass fraction separate from the starch-enriched biomass fractions (i.e., corn grains). Since the carbohydrate polymers trapped within cell walls are highly recalcitrant to chemical or enzyme-catalyzed conversion into soluble fermentable sugars like hexoses (e.g., glucose, mannose) or pentoses (e.g., xylose, arabinose), cellulosic biomass is first pretreated by heat and chemicals (e.g., acid) to make the underlying carbohydrate polymers more readily accessible to hydrolytic enzymes like cellulases or hemicellulases. During pretreatment some of the carbohydrate polymers are partially hydrolyzed into soluble oligosaccharides (e.g., *β*-1,4-glucooligosaccharides like cellobiose, *β*-1,4-xylooligosaccharides like xylobiose). Schematic outlines some of the structural changes that take place within corn plant cell walls during pretreatment based on previously published work.^2,16^ Thermochemical pretreatment of biomass can also significantly alter biomass composition and produce a multiple product streams enriched in carbohydrates.^2^ Next, a complex suite of carbohydrate-active enzymes (CAZymes) is then needed to efficiently hydrolyze the pretreated biomass-derived polysaccharides/oligosaccharides into intermediate soluble sugars (e.g., hexose and pentose sugars like glucose and xylose, respectively). Here, we highlight a commercially-relevant cellulase enzyme (Cel7A from *Trichoderma reesei* with available PDB 8CEL crystal structure^19^) with cellulose chain within its active site docked onto the crystalline cellulose microfibril surface to showcase an important unit operation for solubilization of cellulose-to-glucose.^20^ Finally, the cellulosic biomass derived hydrolysate enriched in soluble sugars can be upgraded into desired bioproducts (like ethanol or other industrial-grade biochemicals) via chemical catalysis or fermentation using engineered microbes. In summary, as highlighted in Figure 1, in order to teach high-school or university students core chemistry, biochemistry, or chemical engineering principles relevant to cellulosic biorefinery processes would require development and validation of various assay methods that are suited for accurate quantification of plant-derived carbohydrates.

There are several commercially available methods for analysis of carbohydrates in human blood or food that have been adapted previously for various student laboratory experiments as also highlighted in a recent ACS Chemical Reviews article.^21^ Briefly, these analytical methods can be classified into two common approaches; (a) Spectrophotometry coupled to colorimetric based sugar detection assays,^11,13^ and (b) Amperometry coupled to enzyme-based sugar detection assays.^22^ Colorimetric based assays were first developed over 150 years ago to quantify glucose in urine and blood samples to diagnose diabetic patients and have been adapted over the years as part of several undergraduate-level teaching and clinical laboratories.^23^ Bio-Rad Laboratories Incorporated developed a biofuel enzyme instructional kit for students to explore the *Aspergillus* derived cellobiase enzyme kinetics using model soluble cellulosic substrates tethered to chromophoric leaving groups (e.g., p-nitrophenyl based cellobiose).^24^ But, as shown previously for CAZymes active on disaccharides,^25,26^ unnatural nitrophenyl-derivatized soluble enzyme substrates often result in biochemical kinetic parameters quite different than those measured on unmodified natural substrates of increasing compositional complexity (e.g., cellodextrins, cellulose, or lignocellulose). In the last two decades, with the development of inexpensive and reliable blood glucose dry-reagent test strips, educators have now adapted the use of simple hand-held glucometers in lower-level biochemistry laboratories to teach students enzymatic hydrolysis kinetics for model disaccharides (e.g., lactose and sucrose).^22^ Researchers from the DOE-funded Great Lakes Bioenergy Research Center (GLBRC) have also adapted these glucometers in a flexible K-16 level lab activity to monitor conversion of cellulosic biomass to ethanol (CB2E) using a complex sequence of processing steps relevant to modern-day biorefinery operations. The CB2E protocol employs hot-water biomass pretreatment to reduce biomass recalcitrance to bioconversion, followed by cellulase enzymes addition to catalyze hydrolysis of pretreated biomass polysaccharides into soluble sugars, and finally followed by yeast catalyzed fermentation of solubilized glucose into bioethanol.^4,27^ The CB2E lab protocol provides students and instructors with detailed instructions on how to produce biofuels starting from diverse plant-derived materials (e.g., corn stalks, grasses, sawdust, or cardboard) over the course of multiple class periods using inexpensive and widely-available reagents/consumables. The published CB2E protocol can be readily used as an introductory science lab for high school students, university students, or in a general public-outreach setting (https://energy.wisc.edu/education/for-educators/educational-materials/cb2e-converting-cellulosic-biomass-ethanol). The CB2E protocol has been also successfully adopted at the University of Wisconsin as part of a one semester based twelve-part bioenergy lab series for undergraduates taught by bioenergy educators at the GLBRC and Wisconsin Energy Institute (https://energy.wisc.edu/education/for-educators/educational-materials/bioenergy-lab-series). However, the CB2E protocol has limited utility for use with advanced high-school or university students as a detailed laboratory experiment due to the lack of validated analytical methods to perform detailed carbohydrates-based compositional analysis on insoluble plant biomass feedstocks and biomass-derived aqueous complex mixed-sugar hydrolysates. For example, it is not possible for instructors to know *a priori* the relative sensitivity of the glucometer-based test strip methods to various biomass-specific carbohydrates since this closely depends on the exact type of enzyme-coenzyme system immobilized on the commercial glucometer test strip biosensor. Also, currently none of these test-strip glucometers have been utilized to estimate the composition of soluble sugars in a non-blood matrix background which results in inaccurate direct meter-reading based quantitation of biomass-specific carbohydrates without the use of calibration measurements using suitable sugar standards. This makes it challenging for instructors to reliably use such assay methods in advanced high school or university laboratory activities focused on lignocellulose biorefining. Overall, the lack of accurate and validated biomass specific carbohydrates assays is one of the major shortcomings of the current CB2E protocol.

Here, we analyze and validate four leading spectrophotometric (i.e., dinitrosalicylic acid and hexokinase based reducing sugars and glucose detection, respectively) and amperometric (i.e., glucose oxidase and glucose dehydrogenase enzyme-coenzyme based glucometer test strips) methods that can be used to accurately quantify plant specific carbohydrates composition. We demonstrate how our validated glucometer based soluble sugars analytical method can be incorporated into a simple hands-on experiment conducted over an introductory ∼90 mins outreach lab activity (called ‘Grass-to-Gas!’) with pre-university high school students to characterize the conversion kinetics of insoluble cellulose polysaccharides and soluble cellobiose disaccharides hydrolysis into glucose using suitable commercial-grade CAZymes. Lastly, we provide several advanced examples of how these analytical methods could be incorporated by bioenergy educators, either in an advanced high-school or university lab setting, to help students explore and understand advanced chemistry, biochemistry, and chemical engineering concepts in the context of lignocellulose biorefining.

### Experimental Section

Four leading soluble carbohydrate sugar assay methods were specifically evaluated based on the following six criteria; ease of use with limited technical training, analyte specificity, detection sensitivity, quantification accuracy, commercial availability, and total cost per assay (**Table 1**). In all cases, we developed calibration measurements for a range of sugar concentrations typically expected in bioenergy focused laboratory protocols (i.e., 0.1-5.0 g/L or 10-500 mg/dL range). More detailed information regarding the soluble sugar assay protocols and our detailed findings are presented in supplementary information.

**Table 1.**
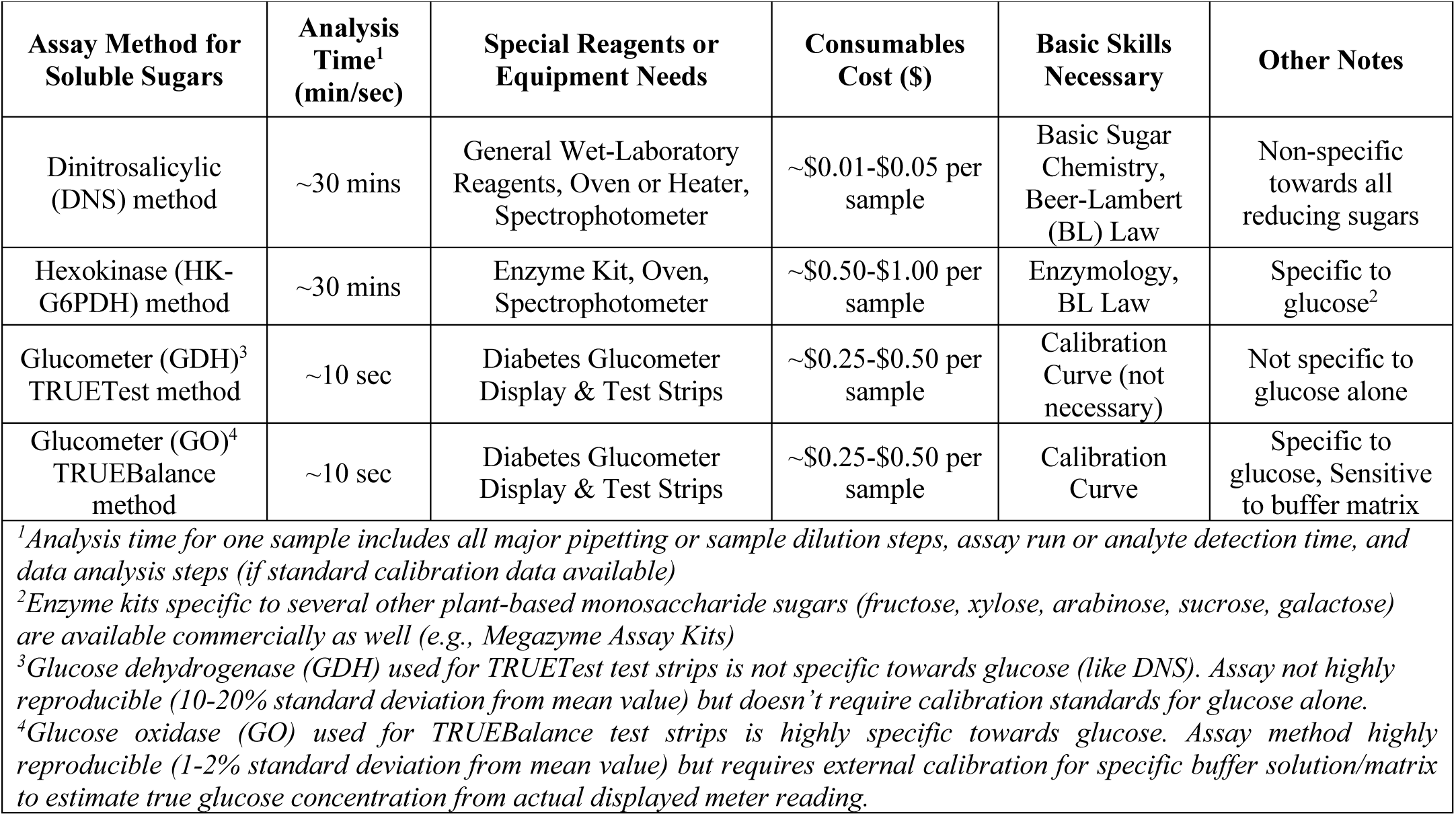
Comparison of four leading analytical methods for soluble carbohydrates analysis for bioenergy focused teaching and hands-on experimental student activity protocols. See SI-I to SI-III appendix for detailed results for all reported soluble sugar assay methods.

Briefly, the following four types of analytical methods were specifically evaluated as described below:

(A) DNS (3,5-dinitrosalicylic acid) based total soluble reducing sugars assay method was used to estimate total soluble reducing sugars in solution using a classical colorimetric method first developed in the 1950’s.^28^ Our protocol was developed based on Miller’s reducing sugar analysis protocol as outlined in the literature and a scaled-down microplate or microtube based version of Miller’s protocol.^28–30^ DNS reacts with all reducing sugars to form 3-amino-5-nitrosalicyclic acid which strongly absorbs light at 540 nm wavelength range under alkaline conditions. This includes sugar aldehydes capable of acting as reducing agents such as most plant-derived monosaccharides (e.g., glucose), disaccharides (e.g., cellobiose), oligosaccharides (e.g., cellodextrins), and polysaccharides (e.g., cellulose). Therefore, while the DNS method has low specificity towards a single monosaccharide or reducing sugar aldehyde,^28,31^ this method allows using an inexpensive colorimetric method to estimate total reducing sugars concentration (as equivalents of desired standard sugar like glucose). The DNS reaction is accompanied by a color change from a bright yellow (i.e., the natural color of the DNS containing reagent stock solution) to dark red-brown color. Dissolved sugar samples containing a higher reducing sugar concentration turning to an increasing darker red hue. Samples that contain higher concentrations of reducing sugars thus display higher absorbances upon absorbing 540 nm wavelength light. For our current method we used a standard spectrophotometer to measure the absorbance intensity of the color change. However, a suitable phone camera and image processing app may be used as well, as highlighted in a recent *Journal of Chemical Education* article.^11^ Finally, we plotted the measured absorbance values against known standard sugar concentrations (e.g., glucose) to allow the user to generate standard calibration curves that were used to determine the unknown concentrations of reducing sugars present in biomass-derived samples.

(B-C) Over-the-counter available blood glucose test strips/meters using two distinct families of immobilized enzymes (TRUEBalance & TRUETest from Trividia Health; http://www.trividiahealth.com) were specifically evaluated here, namely; Glucose Oxidase versus Glucose Dehydrogenase based glucometer test strip kits. These glucometer test strip kits broadly cover two major classes of all commercially available glucometer kits. Commercial blood glucose meters are readily available in most pharmacy stores over-the-counter for diabetic patients to track fluctuations in patient blood glucose levels. Various types of blood glucose meters are available in the market that vary in the type of enzyme-coenzyme specific detection chemistry and/or sensitivity for monitoring soluble sugars like glucose.^23,32,33^ The intensity of electrical current generated is specific to the chemical or electrochemical steps undertaken by the enzyme-coenzyme system immobilized on the biosensor device (i.e., test strip) and can be measured by a suitable glucometer device (i.e., digital meter). Amperometric glucose detection methods utilize changes in measurable current that is proportional to the actual sample glucose concentration to quantify glucose and are often pre-calibrated for glucose detection specifically in the blood matrix. Note that the TRUEtest test strip amperometrically estimates reducing sugars utilizing a non-specific *Acinetobacter calcoaceticus* derived glucose dehydrogenase-pyrroloquinoline quinone (GDH-PQQ) co-factor based immobilized enzymatic reaction. While the TRUEbalance test strip amperometrically estimates glucose only utilizing a highly specific *Aspergillus* sp. derived engineered glucose oxidase (GO) based immobilized enzymatic reaction. Both test strip reactions are accompanied by a change in the digital meter reading, with samples containing a higher concentration of reducing sugars giving a higher meter reading. However, since these test strips and glucose meters have been optimized for estimating the concentration of either reducing sugars or glucose in blood plasma, suitable calibration curves need to be generated for the meter readings using standard sugar solutions from biomass-based sources. Finally, plotting the glucometer digital output reading for known standard sugar concentrations (e.g., xylose dissolved in water) allows generation of standard calibration curves that can be used to determine the unknown concentrations of reducing sugars in desired samples. We tested the sensitivity of both glucometer assay kits for a range of biomass specific carbohydrates (e.g., cellobiose, glucose, xylose, arabinose, mannose, and galactose). We also determined the sensitivity of both these assay methods to common interfering matrix components expected in the sugar solutions. The matrix components in samples include sulfuric acid based neutral salts (e.g., sodium sulfate) expected to be present post-sulfuric acid based hydrolysis or acidic pretreatments generated samples often seen in typical lignocellulose biorefining operations.

(D) Finally, a representative commercially available clinical laboratory-grade enzyme kit was used to estimate glucose concentration using the hexokinase & glucose-6-phosphate dehydrogenase or HK-G6PDH enzyme system (glucose detection kit from Catachem; http://www.catacheminc.com/). The HK-G6PDH assay was based on a two-step enzymatic reaction method, where D-Glucose was first phosphorylated to D-glucose-6-phosphate (G6P) using ATP and a glucose-specific Hexokinase (HK) enzyme. The G6P was then reacted with oxidized nicotinamide adenine dinucleotide (NADP+) by glucose-6-phosphate dehydrogenase to form D-gluconate-6-phosphate and reduced NADPH. The reactions are stoichiometric to the amount of D-glucose and thus the corresponding increase in NADPH concentration can be measured at 340 nm to estimate glucose concentrations based on a suitable standard calibration curve. We also determined the sensitivity of this assay method to common interfering matrix components (e.g., sodium sulfate). Alternative enzyme-based glucose assay kits can be procured and used from Megazyme (Bray, Ireland) and R-Biopharm (Marshall, Michigan), if necessary. Furthermore, for analogous accurate estimation of xylose and arabinose, one could use other monosaccharides specific enzyme-based kits available from other commercial vendors (like Megazyme).

Supporting Information (SI) section also provides additional supporting information (SI-I to SI-VII) relevant to the results reported in the paper. See the following SI appendix sections for details (SI Section No., SI appendix page number); Reducing Sugars Estimation Using DNS Supporting Information (SI-I, p. S2), Glucose vs. Reducing Sugars Estimation Using Glucometers Supporting Information (SI-II, p. S6), Enzyme Kit Based Glucose Estimation Supporting Information (SI-III, p. S11), Insoluble Biomass Composition Analysis Supporting Information (SI-IV, p. S15), Biomass Acid Chlorite Delignification Pretreatment Supporting Information (SI-V, p. S24), Crystalline Cellulose Decrystallization Pretreatment Supporting Information (SI-VI, p. S27), and Cellulosic Biomass Enzymatic Hydrolysis Supporting Information (SI-VII, p. S29). Note that the SI section also provides a detailed ‘Grass-to-Gas!’ Outreach Student Activity & Relevant Supporting Materials (SI-VIII to SI-X) in word/pdf/excel document formats. See the following SI appendix sections for details (SI Section No., SI appendix page number); ‘Grass to Gas!’ Outreach Activity Student Questionnaire-Background Information (SI-VIII, p. S35), ‘Grass to Gas!’ Outreach Activity Student Protocol (SI-IX, p. S38, ‘Grass to Gas!’ Outreach Activity Instructor Protocol & Notes (SI-X, p. S40).

### Hazards

Dinitrosalicylic (DNS), sodium hydroxide, and sulfuric acid are all corrosive and harmful to human health if accidentally touched, inhaled or swallowed. All DNS-based reagents (particularly if containing phenol) are toxic and therefore must be handled with care by users and properly disposed. HK-G6PDH enzyme kit reagents solutions are not to be consumed and enzyme/sugar containing buffers/solutions may cause eye/skin irritation. Commercial CAZyme based enzymes solutions may cause skin sensitization. Researchers must wear laboratory gloves and eye protection glasses at all times when running sugar assays. None of these analytical methods or calibration curves should be supposed to quantify soluble sugar concentrations in bodily fluids to draw any relevant medical conclusion. See supplementary information for specific hazards associated with other supporting bioenergy laboratory protocols relevant to this work.

## Results and Discussion

All experimental protocols were established and executed by closely mentored undergraduate students in the bioenergy-focused research laboratories at the University of Wisconsin-Madison and Rutgers-The State University of New Jersey. We first provide detailed comparison of the four soluble sugar assay methods and discuss their suitability for use by educators during cellulosic bioenergy relevant hands-on student experimental activities. We also share a detailed student instruction protocol that utilized one of these validated sugar assay methods and overall results from our hands-on outreach activity conducted at Rutgers University with pre-university high school students. This outreach activity involves students running biochemical assays to help them understand the various facets of insoluble cellulose hydrolysis by commercial cellulase enzymes and monitoring the total glucose product released using our validated glucometer assay to estimate total cellulose-to-glucose conversion efficiency. Finally, we discuss three advanced applications of how such carbohydrate analytical methods could also be adapted and used by bioenergy educators to; (a) determine the overall carbohydrate composition of biorefinery relevant lignocellulosic biomass feedstocks to setup a more advanced CB2E protocol, (b) perform material balances around biorefinery unit operations such as pretreatments, and (c) analyze the relative enzymatic hydrolysis kinetics of biorefinery-relevant cellulosic substrates of increasing compositional complexity (i.e., soluble cellobiose or cellodextrins, insoluble microcrystalline cellulose, and pretreated lignocellulosic biomass).

### Comparison of carbohydrate assays

First, we compared the analytical accuracy and analyte specificity of the four spectrophotometric and glucometer-based amperometric methods to directly quantify concentrations of soluble carbohydrate mono-/di-saccharides often found in cellulosic biomass or biorefinery processes. The HK-G6PDH enzyme kit and TRUEBalance assays alone were highly specific to detection of glucose only in all tested soluble sugars (i.e., cellobiose, glucose, xylose, arabinose, mannose, and galactose). However, as expected, the DNS based colorimetric method and the TRUETest assays were sensitive to all major lignocellulose-derived reducing sugar aldehydes.

In terms of detection sensitivity, the DNS assay method tested gave a linear calibration curve response for sugar concentrations ranging from 0.1-5 g/L (or 10-500 mg/dL). The TRUEBalance assay method tested gave a linear calibration curve response for sugar concentrations ranging from 0.5-2.0 g/L (or 50-200 mg/dL). The TRUETest assay method gave a direct displayed meter reading for glucose standards that was within 25% of the expected value (for glucose standards in the 0.5-1 g/L range), while the TRUEBalance assay method gave a ∼2 fold higher displayed meter reading than the actual expected value suggesting that the latter method was highly sensitive to the solution matrix/buffer composition (**Figure 2**). However, since the TRUETest method was not specific towards glucose alone, one should use the TRUEBalance method to monitor glucose for mixed biomass derived sugar hydrolyzates (and based on an external glucose calibration dataset) while the TRUETest should be used when glucose alone is present (and in the absence of an external calibration dataset for glucose). The exact fold difference in actual versus displayed meter reading may depend on the actual age and lot number of the glucometer test strips used. Thus, older glucometer test strips should be checked often using a known glucose standard control prior to use and calibration curve should be generated for known standards on the day of use. The HK-G6PDH enzyme kit tested gave a linear calibration curve response for sugar concentrations ranging from 0.5-5.0 g/L (or 50-500 mg/dL). Note that if the assay results were beyond the upper range of the calibration curve in any case, then the samples were diluted in stock buffer solution or deionized water and reanalyzed.

**Figure 2.**
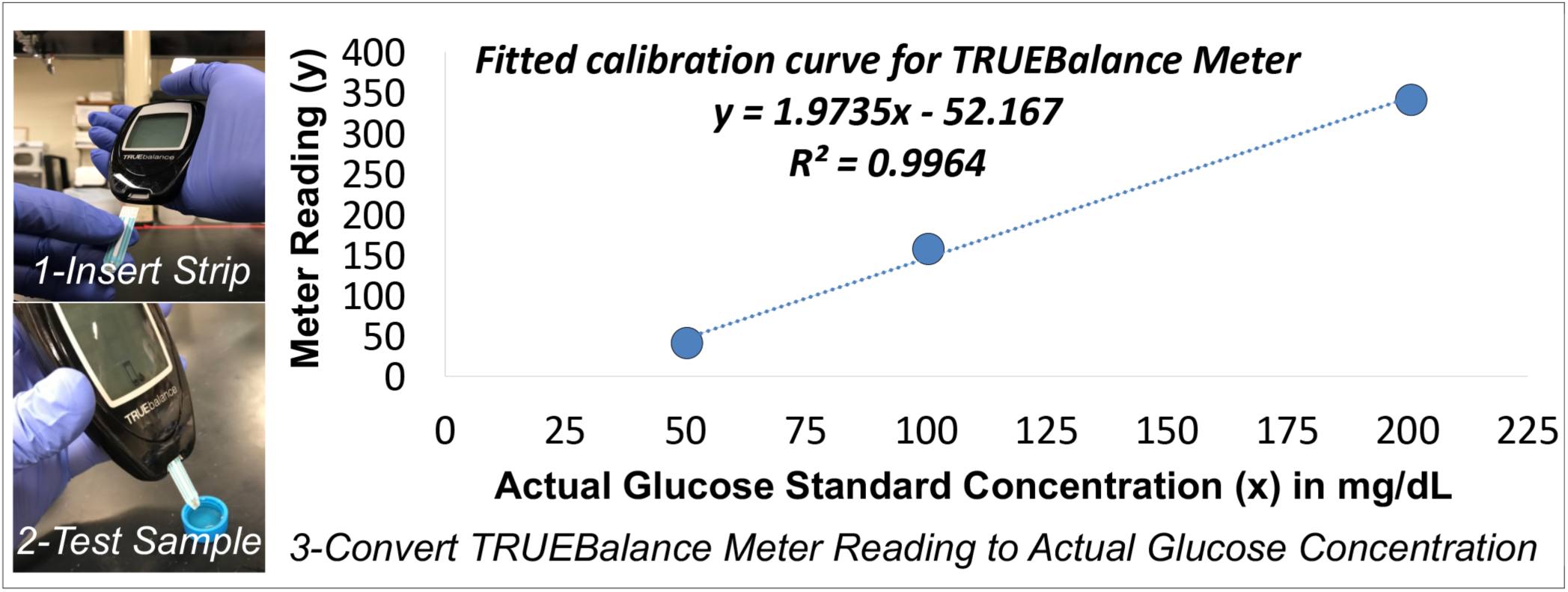
Glucose-oxidase based glucometer method (i.e., TRUEBalance) for detection and quantification of soluble glucose in plant-biomass derived samples. Here, we outline a simple 3-step process to rapidly and specifically detect and quantify glucose present in biomass derived solutions. The TRUEBalance test strip shows no interference for other soluble sugars but requires use of a calibration curve (with standards dissolved in identical sample buffer/matrix) to convert meter readings to actual glucose concentration (mg/dL). See SI-II appendix for additional details regarding the glucometer-based assays for detecting plant-based carbohydrates.

Sodium sulfate is a major byproduct formed during NaOH catalyzed neutralization of sulfuric acid containing biomass hydrolysis samples that is useful for estimation of total insoluble cellulose/hemicellulose composition of biomass samples or analysis of acid pretreatment generated samples. Only the TRUETest and HK-G6PDH showed no significant interference in the presence of 1M Na2SO4 salt concentration. But note that users can still use the TRUEBalance method if a calibration curve is generated using sugar standards of known sugar concentrations spiked with high Na2SO4 concentration to account for salt matrix interference.

In terms of analytical reproducibility, most methods (i.e., DNS, TRUEBalance HK-G6PDH) gave a standard deviation of under 2% from mean reading for all deionized water dissolved glucose standards analyzed. However, the TRUEtest method gave much higher standard deviations of 10-20%, suggesting that this test strip assay was not highly reproducible. This work highlights the variability in some glucometer test strips for detection of sugars in non-blood based samples and under non-ideal self-testing conditions.^23,34^ However, this could also reflect the manufacturer associated product quality issues as suggested by a recent 2016 FDA regulated recall of some Trividia Health TRUETest glucometer test strips.^35^

In terms of ease of use, only the glucometer-based kits required limited user training with minimal handling of pipettes/micropipettes if all stock reagents and standards are prepared ahead of time by the instructor or support staff. The glucometer-based detection methods are suitable for use with mostly high-school or first-year university students with limited laboratory experience and/or limited knowledge about the relevant analytical method. The DNS and HK-G6PDH based assays require more extensive training of students with extensive handling of pipettes/micropipettes to reproducibly maneuver microliter volumes between microtubes/microplates and utilization of a UV-Vis spectrophotometer for quantification of response. The non-glucometer based detection methods are thus more suitable for use with mostly advanced university students with some laboratory experience and knowledge of basic organic/physical chemistry concepts.

In terms of overall assay time, the glucometer-based assays give a readout for each sample within ∼10 secs to allow students to estimate unknown sample concentrations (assuming standards are prepared ahead of time by instructor or lab staff). However, the DNS and HK-G6PDH based assays require a total assay time of under ∼30-45 mins based on total number of samples analyzed by each student and the users pipetting skills to estimate unknown sample concentrations (assuming standards and stock reagent solutions are prepared ahead of time as well). However, the glucometer-based assay kit is not setup for running multiple samples at once, while the DNS and HK-G6PDH based assays are amenable to be run in a multiplexed manner using 96-well based microtubes/microplates and a UV-Vis microplate reader. Lastly, in terms of ancillary equipment or reagents/consumables, glucometer-based assays do not require access to significant plastic consumables or even specialized equipment (like oven or heating block, UV-Vis spectrophotometer). However, the cost per sample assayed for reagents/consumables alone is the highest for HK-G6PDH based methods (∼$0.5-1.0), followed by glucometer-based assays (∼$0.25-0.5), and then by DNS assays (∼$0.01-0.05). In terms of commercial availability, supplies and reagents for all assay methods are readily available from multiple companies either at over-the-counter pharmacies for glucometers or through common laboratory grade reagent supplier (e.g., VWR/Sigma-Aldrich/Fisher).

### Carbohydrate assays used in outreach student activity to study enzymatic hydrolysis of cellulose

Enzymatic hydrolysis is a key biorefinery process for converting pretreated biomass embedded polysaccharides into sugars using synergistic cellulose and hemicellulose targeting CAZymes as biocatalysts. We showcase how our analytically validated hand-held glucometer assay method can be used as part of an instructional activity for students to monitor the enzymatic hydrolysis kinetics of model cellulose substrates into glucose. The glucometer assay method was incorporated into an actual bioenergy-focused outreach activity (∼90 min total experimental time and ∼30 min data analysis time) to demonstrate the simplicity and utility of such an analytical method to help pre-university school students understand and explore enzyme biochemistry concepts relevant to cellulosic bioenergy production. This outreach activity, called ‘Grass-to-Gas!’, has been conducted with over 70+ high school students so far as part of a cellulosic biofuels public outreach event conducted at Rutgers University on two separate occasions (July 14^th^ 2017 and July 20^th^ 2018). Detailed information regarding the ‘Grass-to-Gas!’ student activity handout and instructor notes are presented in the supplementary information.

Briefly, a simplified protocol was developed to allow students to test the relative hydrolytic activity of commercially available cellulase/hemicellulase based CAZyme cocktails on either soluble or insoluble cellulosic biomass based on the original NREL protocol (https://www.nrel.gov/bioenergy/biomass-compositional-analysis.html).^36^ Our current method, inspired by NREL’s method, utilizes readily available lab accessories like a hot water-bath (and/or regular oven) and common chemistry lab apparatus to setup reactions to hydrolyze the carbohydrate polymers like microcrystalline cellulose (e.g., Avicel PH-101), amorphous cellulose (e.g., derived from Avicel PH-101), and/or cellodextrins (e.g., cellobiose) into soluble sugar monomers like glucose using commercially available cellulase (e.g., Celluclast from Novozymes) or cellobiase (e.g., Novo 188 from Novozymes) enzyme cocktails. Immediately at the start of the activity session, the students are instructed to pipette the desired enzyme-buffer stock solution volumes into pre-weighed cellulosic substrates containing 15-ml plastic tubes and then incubating the tubes at the desired reaction conditions (∼15-30 mins). The instructor next introduces basic concepts relevant to cellulose structure, enzyme biochemistry, and glucose quantitation using glucometer assay kits over the next ∼60-90 mins during the course of enzymatic hydrolysis. The students are then split into various sub-groups to investigate the effect of various reaction parameters on enzymatic hydrolysis, namely; biomass type (soluble vs. insoluble), total enzyme loading (per unit mass cellulose), total cellulose loading (per unit reaction volume), reaction temperature, and reaction time. The soluble glucose released during the course of enzymatic hydrolysis is then assayed by students individually using the TrueBalance glucometer test strips kit followed by data analysis over the last ∼30 mins of the activity. All students are provided with a glucometer calibration curve dataset (to convert displayed meter reading into actual glucose concentration in mg/dL) and a list of handout queries to engage the students and promote group discussions. Relevant student activity protocols, background information handout, and questionnaire documents are provided in the supporting information section.

Representative results from enzymatic hydrolysis of microcrystalline and amorphous cellulose expected for this outreach activity at various enzyme loadings, cellulose concentrations, and reaction temperatures are summarized in **Figure 3**. Amorphous cellulose typically gives much higher cellulose-to-glucose yields compared to microcrystalline cellulose (at 50 mg enzyme/g cellulose loading) clearing showing that crystalline cellulose is significantly more recalcitrant to enzymatic hydrolysis. The underlying mechanisms responsible for varying susceptibility of different cellulosic substrates by cellulases is an active area of research and could be a topic for discussion at the end of this outreach activity.^37–40^

**Figure 3.**
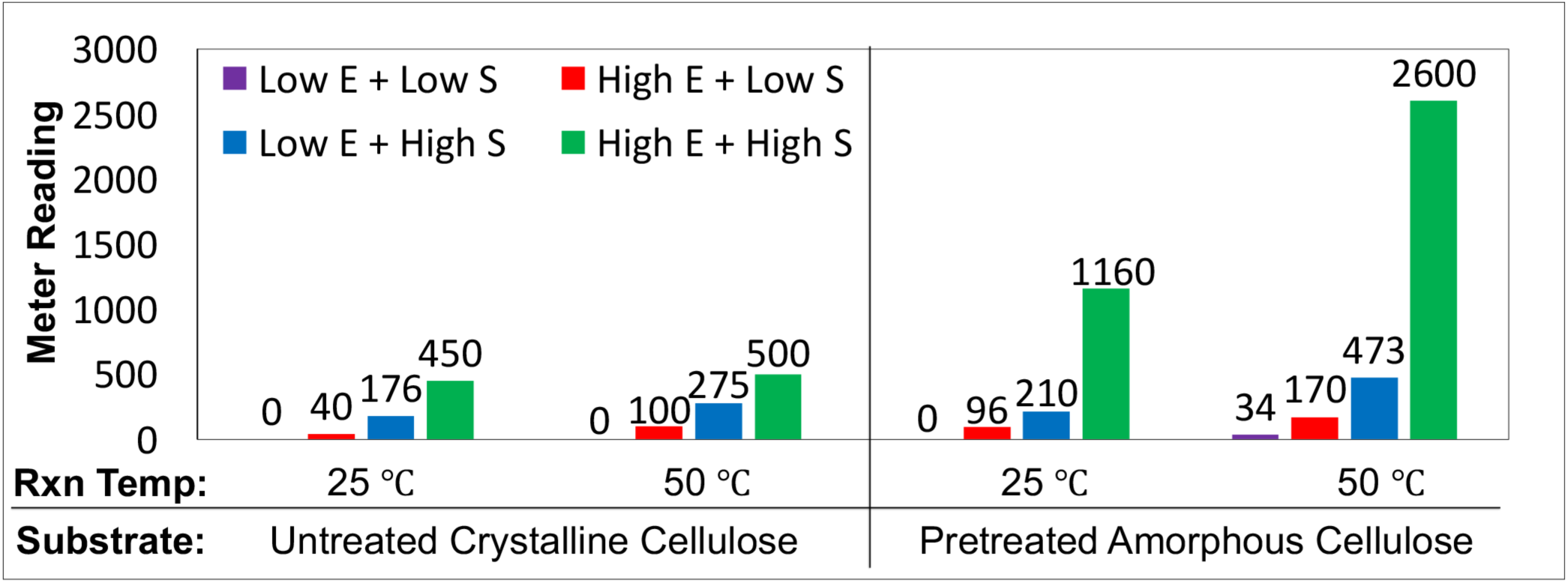
Representative student results from ‘Grass-to-Gas!’ outreach activity conducted to explore the effect of cellulase enzyme (E) concentration, cellulosic substrate (S) concentration, hydrolysis reaction temperature, and cellulose pretreatment on the relative rate of cellulosic substrate conversion into glucose mimicking a cellulosic biorefinery enzymatic hydrolysis unit operation. A total reaction volume of 10 ml in a stoppered plastic test tube, containing either 25 (Low S) or 250 (High S) mg of untreated microcrystalline cellulose (i.e., Avicel PH-101) or pretreated amorphous cellulose (i.e., Phosphoric Acid Swollen Cellulose or PASC) was hydrolyzed using a commercially available cellulase enzyme cocktail at either a 5 mg enzyme per gram cellulose (Low E) or 50 mg enzyme per gram cellulose (High E) enzyme loading at two different reaction temperatures (i.e., 25 and 50 °C). Avicel PH-101 was pretreated and decrystallized using concentrated phosphoric acid pretreatment into highly amorphous cellulose as detailed in the supplementary information SI-VI appendix. The enzymatically released soluble sugars (i.e., glucose) was measured using the TRUEBalance test kit after 90 mins of total reaction time (under static conditions with intermittent manual end-over-end mixing every 30 mins) with the actual glucometer readings reported here. Commercially available C_TEC2 cellulolytic enzyme cocktail (gift from Novozymes) was used for assays here. See supplementary information for detailed student activity protocol and accompanying instructor handout (SI-VIII to SI-X).

### Carbohydrate assays applied to advanced bioenergy laboratory protocols

Finally, we discuss additional examples here (and provide protocols in the supplementary section) on how carbohydrate analytical methods can be incorporated into simplified hands-on laboratory experiments appropriate for university students to be introduced to advanced chemistry, biochemistry, and chemical engineering concepts in the backdrop of cellulose biorefining and bioenergy production.

### (a) Carbohydrate assays to analyze lignocellulosic biomass composition

First, we showcase how our validated carbohydrate analytical methods can be used to accurately quantify concentrations of soluble carbohydrates released from insoluble polysaccharides using a modified acid hydrolysis protocol developed to estimate the carbohydrate polysaccharides composition of diverse bioenergy feedstocks. This would allow students to determine total carbohydrate composition of biorefinery relevant lignocellulosic biomass feedstocks to ultimately develop a more advanced CB2E protocol or perform overall carbohydrate material balances around individual biorefinery unit operations (like pretreatment). In order to achieve this goal, a simplified acid hydrolysis protocol was first developed to hydrolyze insoluble biomass polysaccharides (e.g., cellulose and hemicellulose) present within biomass feedstocks using sulfuric acid based on the original protocol developed by the National Renewable Energy Laboratory (NREL) (http://www.nrel.gov/biomass/analytical_procedures.html).^41^ The original NREL protocol recommends using a high-pressure acid hydrolysis procedure to hydrolyze the carbohydrate polymers (oligosaccharides and polysaccharides) into easily quantifiable soluble sugars (e.g., monosaccharides) utilizing specialized equipment (e.g., pressurized glass-reactor vessels, autoclaves) to estimate the total carbohydrate composition of plant-derived biomass. The released monosaccharides (e.g., glucose, xylose) are then separated and quantified using a high-pressure liquid chromatography system equipped with a suitable universal detector (e.g., Refractive Index, Light Scattering). However, since such equipment are not often available in most high schools and some undergraduate teaching laboratories, we modified the compositional analysis protocol to be more suitable for use by bioenergy educators in a typical university chemistry or biochemistry teaching laboratories (and possibly be used even in a typical high-school science lab). We have provided a detailed protocol in the supplementary information section that can be used by instructors to analyze biomass samples composition prior to application of the published CB2E protocol. Educators could also consider using this protocol as a general chemistry or biochemistry laboratory experiment held over multiple classes in a semester. Detailed information regarding insoluble polysaccharides acid hydrolysis step-by-step method (with illustrative pictures), soluble sugar assay protocols, and detailed formulas used to calculate biomass composition (with examples provided for feedstocks reported here) are presented in the supplementary information document.

Briefly, our simplified biomass composition analysis method, inspired by another published work,^42^ utilizes a multi-step acid hydrolysis procedure using a hot water-bath (and/or regular oven) alone and common school/university chemistry lab apparatus to hydrolyze the insoluble carbohydrate polymers into soluble sugar monomers without use of any high-pressure autoclave or specialized equipment/glassware. The soluble sugars released in the biomass acid hydrolyzate solution are first neutralized and then assayed using two different soluble sugar assay methods to finally estimate the initial cellulose and hemicellulose composition of the biomass feedstock. We used the DNS assay to indirectly estimate total soluble reducing sugar concentration (i.e., hemicellulosic polymer content) and the HK-G6PDH enzyme kit to directly estimate the total glucose concentration (i.e., glucan polymer content). Note that the TrueBalance glucometer assay method can also be used in place of the HK-G6PDH assay as well. Our biomass acid hydrolysis method, in combination with our validated soluble sugar assays, was first used to analyze the insoluble polysaccharides composition for a diverse range of leading cellulosic feedstocks (e.g., waste cardboard, switchgrass, corn stover/stalks, and Aspen wood) that were used previously in the CB2E protocol.^4^ Based on our sugar assays, we could roughly estimate the total glucan (or cellulose) and non-glucan (or hemicellulose) composition of the original starting biomass. Importantly, it was possible to achieve comparable cellulose and hemicellulose compositional results to what have been reported previously using advanced NREL protocols for comparable feedstocks (http://www.nrel.gov/biomass/data_resources.html). See **Table 2** outlining composition analysis results obtained for a diverse range of outreach activity relevant feedstocks that were analyzed using the methods outlined here. The compositions of diverse biomass feedstocks (e.g., Corn stover, Switchgrass, Aspen wood, Cardboard) currently used in the CB2E protocol were estimated. This comprehensive compositional dataset will allow students to understand why cardboard as a feedstock, with its relatively higher starting cellulose concentration for an equal weight of starting material taken for enzymatic hydrolysis at a fixed total enzyme loading, gave the highest glucose yields in the original reported CB2E protocol results.^4^ Adjusting the cellulase enzyme loaded per unit mass of equivalent cellulose (as mg enzyme per unit gram cellulose) would allow students to make more rigorous comparisons between the relative digestion efficiency amongst various cellulosic substrate using similar methods as often reported by bioenergy researchers in peer-reviewed scientific papers.

**Table 2.** Insoluble carbohydrate polymers compositional analysis for diverse cellulosic feedstocks (on dry weight basis) can be reliably estimated, post acid-hydrolysis, using appropriate sugar assays. Here, C1 is microcrystalline cellulose (Avicel PH-101) control, CS is untreated corn stover, D-CS is delignified corn stover, AW is native untreated aspen wood, SG is native untreated switchgrass, and CB is a cardboard sample. D-CS was delignified using the acid-chlorite pretreatment process as reported in SI-V appendix. See supporting section SI-IV appendix for details on how to calculate biomass composition using acid hydrolysis to solubilize polysaccharides into monosaccharides and relevant sugar assay based compositional analysis. Representative experimental results expected for this specific protocol are reported here. One standard deviation (±σ) is shown for triplicate experiments carried out on at least two separate days. ND stands for not detected.

| Cell wall Components | Avicel (C1) | Corn Stover (CS) | Delignified Corn Stover (D-CS) | Switch Grass (SG) | Aspen Wood (AW) | Card Board (CB) |
| --- | --- | --- | --- | --- | --- | --- |
| Cellulose<sup>1</sup> | 99±3 | 39±6 | 69±5 | 42±4 | 47±3 | 72±3 |
| Hemicellulose<sup>2</sup> | ND | 37±10 | 25±5 | 31±7 | 32±5 | 27±4 |
| Lignin & Others<sup>3</sup> | ND | 27±2 | 5±1 | 30±2 | 24±5 | 15±3 |
| Soluble Sugars<sup>4</sup> | ND | 1±0.5 | ND | 2±0.5 | ND | ND |
| Total Cell Wall Content<sup>5</sup> | 99±3 | 103±12 | 99±7 | 103±8 | 103±8 | 114±7 |
| <sup>1</sup>Total glucose released during biomass acid hydrolysis and estimated using either HK-G6PDH or TRUEBalance method<br><sup>2</sup>Difference between DNS or TRUETest assay estimated total reducing sugars of biomass acid hydrolyzate and HK-G6PDH or TRUEBalance estimated glucose concentration of biomass acid hydrolyzate<br><sup>3</sup>Gravimetric analysis of insoluble residue left behind, mostly enriched in lignin, after acid hydrolysis of polysaccharides<br><sup>4</sup>Soluble sugars present in biomass was estimated by boiling 1 g dry weight biomass in 25 ml water for 20 mins followed by DNS or TRUETest assay to estimate total soluble sugars on the water-soluble fraction<br><sup>5</sup>Total cell wall content here is sum of total estimated cellulose, hemicellulose, and lignin/others |  |  |  |  |  |  |

### (b) Carbohydrate assays to study analyze individual biorefinery unit operations

Next, we discuss how instructors can generate pretreated biomass feedstocks of varying compositional complexity to be included in the original CB2E protocol to allow students to compare impact of pretreatment on biomass composition and subsequent enzymatic hydrolysis operations.^4,27^ Pretreatment of biomass with heat and chemical solvents/catalysts can significantly alter biomass composition and produce multiple product streams enriched in carbohydrates.^2^ Here, we highlight two pretreatment protocols that can be used by instructors to generate cellulosic biomass of varying ultrastructural and compositional complexity to be used by students to invoke their curiosity about actual biorefinery processes and the issues associated with biomass recalcitrance towards industrial biorefining. See supplementary information for details on supporting protocols to allow instructors and/or students to conduct two types of biomass pretreatments (using acid chlorite for lignin removal from corn stover and using phosphoric acid to decrystallize microcrystalline cellulose to make amorphous cellulose) as an extension to the current CB2E hot-water pretreatment protocol. These additional pretreatment protocols can be used by the instructor alone to also generate alternative biomass feedstocks for total compositional analysis or to be used in combination with the existing CB2E protocol instead. These pretreated substrates could be specifically included in the enzymatic hydrolysis hands-on experimental activity along with off-shelf commercial grade model celluloses (e.g., Cellobiose, Avicel PH-101) as substrates. These substrates would allow students to explore the rate-limiting role of lignin and cellulose crystallinity on biomass enzymatic hydrolysis.

Here, we provide illustrative results on the composition of acid-chlorite pretreated corn stover pretreated for increasing durations to showcase how this specific pretreatment method allows significant delignification of the biomass to increase relative cellulose concentration comparable to cardboard (on dry weight basis). As discussed in the supplementary section, total lignin content of corn stover decreased from 27% to ∼2% after few hours of acid-chlorite treatment. Furthermore, we show that the relative enzymatic hydrolysis yield for cellulose-to-glucose for untreated corn stover (CS), acid-chlorite pretreated delignified corn stover (D-CS), switchgrass (SG), Aspen wood (AW), and cardboard (CB) was 31%, 85%, 31%, 29%, and 19%, respectively. These results clearly showcase that cellulose-to-glucose hydrolysis yield for delignified biomass is much higher than untreated biomass. Here, 50 mg Celluclast/Novo 188 enzymes each per gram cellulose was the exact enzyme-to-substrate loading (normalized to actual biomass cellulose composition) used to enzymatically hydrolyze all biomass substrates at 50°C for 24 hours (without shaking). The data clearly shows that cellulose in cardboard is actually more recalcitrant towards enzymatic hydrolysis compared to other feedstocks, likely due to extensive hornification of cellulose fibers during the pulping and subsequent paper-making process that makes it more recalcitrant to enzyme access.^43,44^ Composition analysis of these samples was thus critical to accurately estimate the relative fraction of cellulose hydrolyzed to glucose for equivalent enzyme loadings (per unit mass of cellulose). This conclusion would not be possible if instructors/students only follow the originally reported CB2E protocols.

Finally, we provide supporting methods for generation of amorphous cellulose from Avicel PH-101 microcrystalline cellulose to be used as substrates to explore cellulase enzymology as a function of cellulose crystallinity. The amorphous substrates were digested using commercial cellulase enzymes using the reported outreach activity and released sugars were measured using the TRUEBalance glucose assay. Snapshot summaries of typical results expected from amorphous cellulose versus microcrystalline cellulose are already summarized in **Figure 3**. In addition, we have also compared the relative enzymatic hydrolysis rate of amorphous cellulose to cellobiose in the supplementary information document. These assays could also be used to monitor the relative enzymatic hydrolysis kinetics and conversion efficiency of soluble cellodextrins like cellobiose compared to insoluble amorphous cellulose. The cellobiose-cellobiase based biochemical assays can be particularly conducted at various enzyme loadings, substrate concentrations, and reaction temperatures to help students explore various concepts related to biocatalysis and cellobiase enzymology relevant to cellulosic bioenergy. We found that near-complete and 30-35% cellobiose hydrolysis to glucose was achieved within 60 mins using a 10 and 1 mg cellobiase (Novo 188) enzyme loading per g cellobiose, respectively. On the other hand, only 10-15% conversion of amorphous cellulose hydrolysis to glucose was achieved within 60 mins using a 1.5 mg cellulase (Celluclast supplemented with equivalent amount of Novo 188) enzyme loading per g cellulose. Near-theoretical conversion of amorphous cellulose hydrolysis to glucose was only achieved after 24 h using a 1.5 mg cellulase (Celluclast) enzyme loading per g cellulose. These results can help instructors showcase how soluble cellulosic substrates like cellobiose are more readily digestible than insoluble substrates into glucose by the relevant enzymes to highlight the challenges faced during biochemical conversion of cellulosic biomass to fermentable sugars. Furthermore, in order to properly capture cellobiose hydrolysis kinetic profiles, we conducted hydrolysis reactions at a much lower enzyme loading (i.e., 1 mg cellobiase enzyme loading per g cellobiose) as reported in the supplementary section. Students could ultimately conduct more advanced analysis on similar initial hydrolysis rate datasets collected at much lower enzyme loadings (<1 mg Novo 188 loading per g cellobiose) for analyzing cellobiase kinetic parameters on soluble cellulosic substrates (like cellobiose) as part of an extended lab activity using our proposed glucometer assay. A similar approach has been used to determine Michaelis-Menten model fitted kinetic parameters of lactase enzymes using another soluble disaccharide (i.e., lactose).^22^ Instructors could consider including assays on cellobiose alone using the TRUEBalance glucometer assay to monitor glucose released in the background of cellobiose substrate with no interference. These assays would be complementary to the biofuel enzyme kit available from Bio-Rad that use pNP-based substrates to assay cellobiase enzymes kinetic parameters. Lastly, instructors could also use such enzyme hydrolysis datasets for a range of cellulosic substrates to highlight the challenges associated with modeling insoluble substrate enzyme kinetics using overly simplistic Michaelis-Menten type models,^45–47^ unlike soluble substrates like cellobiose.

## Conclusions

We have compared, validated, and proposed a set of carbohydrate detection assays, relevant to quantification of plant-specific carbohydrates in a biorefinery backdrop, that are ideal for student laboratory training and instruction in a high-school or university setting. We demonstrate how our validated glucometer based soluble sugars analytical method were incorporated into a simple hands-on experiment conducted over an introductory ∼90 mins outreach lab activity with pre-university high school students to characterize the conversion kinetics of insoluble cellulose polysaccharides and soluble cellobiose disaccharides hydrolysis into glucose using suitable commercial-grade carbohydrate-active enzymes. Lastly, we provided detailed supplementary protocols and discuss several advanced examples of how these carbohydrate analytical assays could be flexibly adapted either in a hands-on exploratory outreach event or as a comprehensive set of experiments in an undergraduate laboratory to increase student understanding of a range of scientific and engineering-based concepts (e.g., cellulosic biorefinery unit operations, material balances, UV-Vis spectroscopy, enzymology, carbohydrate chemistry, plant and microbial sciences) with the ultimate goal of encouraging student and public enthusiasm towards bioenergy research. These protocols would allow students to systematically inquire and understand some of the open-ended questions relevant to cellulosic biofuels production operations and associated challenges.

## Supporting information

SI-X Activity Excel

## Supplementary Information (SI) & Associated Content

Supplementary information section provides additional supporting protocols/results (SI-I to SI-VII) and Student Outreach Lab Activity Handout with Instructor Notes (SI-VIII to SI-X).

## Acknowledgements

SPS Chundawat acknowledges support from the US National Science Foundation CBET awards (1604421, 1846797), Department of Energy (DOE) Office of Science award (DE-SC0019313), ORAU Ralph E. Powe Award, Rutgers Division of Continuing Studies, and Rutgers School of Engineering. This work was also partially supported by the DOE Great Lakes Bioenergy Research Center (DOE BER Office of Science DE-FC02-07ER64494). We are very grateful to Marin Dobson (Fort Atkinson High School), Cameron Sieser (UW-Madison), Jia-Mei Hong (Rutgers University), and Benjamin Esposito (Rutgers University) for their critical contributions to the execution of all experiments reported here. Special thanks to Leith Nye, John Greenler, Brian Fox, James Runde, Troy Runge, and Kate Helmich for their timely support of part of this work through the GLBRC Research Education for Teachers (RET) Program. Lastly, thanks to Alex Bertuccio for useful feedback and implementation of relevant ‘Grass-to-Gas’ student activity protocols as part of the Rutgers Undergraduate CBE Senior Process Lab (155:416 course) curriculum.

## Cover Art/Graphical abstract & associated caption

Carbohydrate assays focused hands-on activities to facilitate student learning of inter-disciplinary concepts relevant to cellulosic bioenergy production

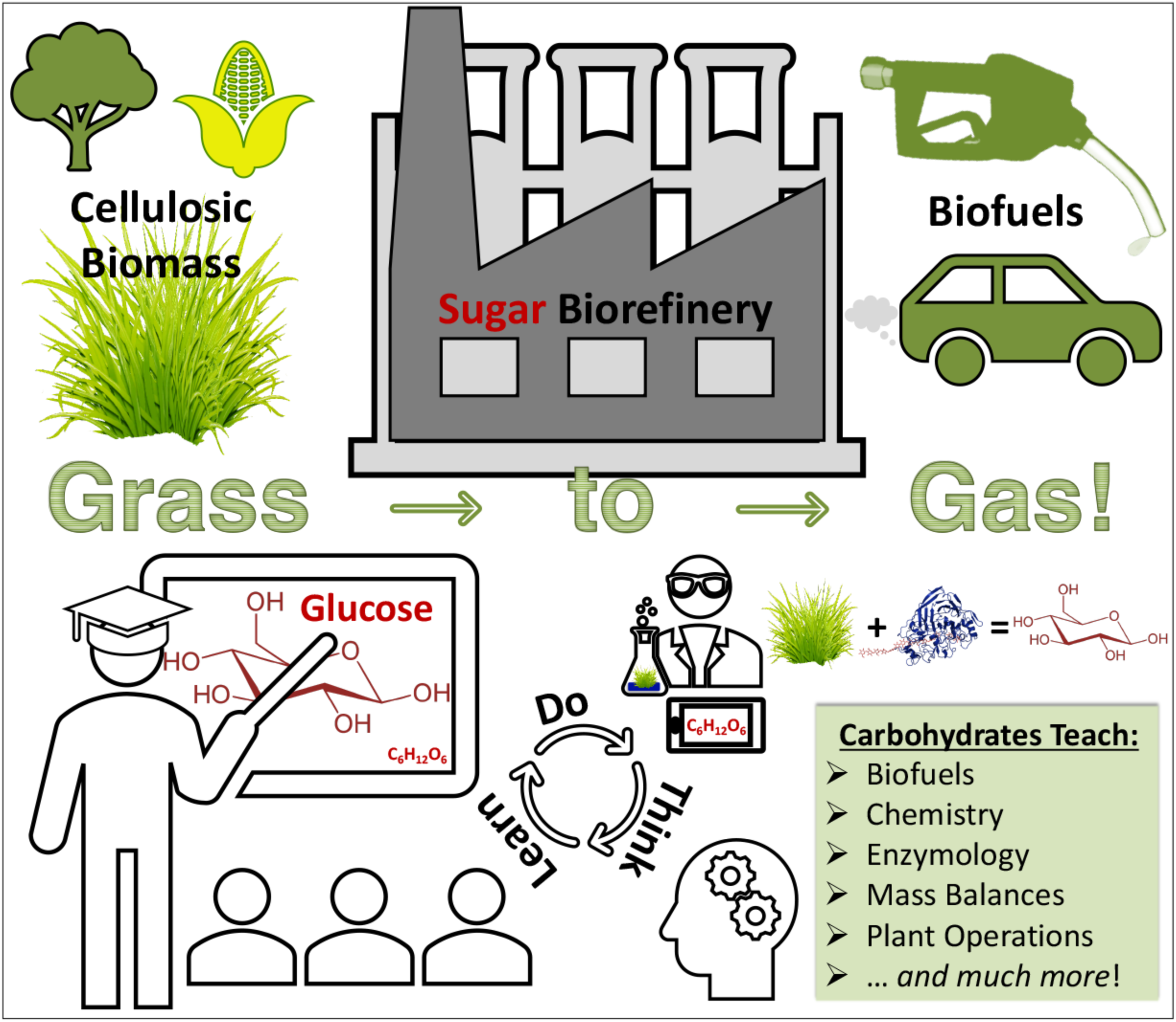

