## Supporting Information for "Comparison of analytical methods for rapid & reliable quantification of plant-based carbohydrates for the quintessential bioenergy educator"

#### Supporting Information (SI) Table of Contents

SI-I. Reducing Sugars Estimation Using DNS Supporting Information.........................................S2

SI-II. Glucose vs. Reducing Sugars Estimation Using Glucometers Supporting Information ............S6

SI-III. Enzyme Kit Based Glucose Estimation Supporting Information........................................S11

SI-IV. Insoluble Biomass Composition Analysis Supporting Information......................................S15

SI-V. Biomass Acid Chlorite Delignification Pretreatment Supporting Information .......................S24

SI-VI. Crystalline Cellulose Decrystallization Pretreatment Supporting Information......................S27

SI-VII. Cellulosic Biomass Enzymatic Hydrolysis Supporting Information...................................S29

SI-VIII. ‘Grass to Gas!’ Outreach Activity Student Questionnaire-Background Information..............S35

SI-IX. ‘Grass to Gas!’ Outreach Activity Student Protocol..........................................................S38

SI-X. ‘Grass to Gas!’ Outreach Activity Instructor Protocol & Notes.............................................S40

SI-XI. References for SI Document.............................................................................................S50

#### (SI-I) Reducing Sugars Estimation Using DNS Supporting Information

***Summary and Principle***: 3,5-dinitrosalicyclic acid (DNS) reacts with reducing sugars (those capable of acting as reducing agents; include all monosaccharides and some disaccharides, oligosaccharides, and polysaccharides) to form 3-amino-5-nitrosalicyclic acid which strongly absorbs light at 540 nm. This reaction is accompanied by a color change from yellow (the natural color of the DNS containing reagent) to dark red, with sugar samples containing more reducing sugars turning to a darker red hue. Samples that contain higher concentrations of reducing sugars will display higher absorbances upon absorbing 540 nm light. Plotting measured absorbance values against known standard sugar concentrations (e.g., glucose) will allow generation of a standard curve that can be used to determine the unknown concentrations of reducing sugars in desired samples. This protocol was developed based on Miller's original reducing sugar compositional analysis protocols outlined in the literature and a microplate version of the same protocol.[1–3] The current method uses a spectrophotometer to measure the intensity of the color change, however, a suitable phone camera and image processing app may be used as well, as highlighted in another recent study.[4]

***List of Chemicals and Materials for Experiment***: Glucose, Sodium Sulfate, 3,5-Dinitrosalicylic Acid, Sodium Hydroxide, Sodium Potassium Tartrate, Sodium Metabisulfite, and Phenol were all procured from Fisher-Scientific, Sigma Aldrich, or VWR to be used as is. Distilled water was used in the preparation of all aqueous solutions (unless specified otherwise).

All necessary glassware, lab equipment, and supplies were used to carry out all experiments are highlighted here: Single/multi-channel micropipettes and associated plastic tips (10, 100, 1000 μL range), Glass pipette (1-10 mL), Falcon conical plastic tubes (15, 50 mL range), Volumetric flask (100, 1000 mL), 96-well clear flat-bottom microplates (0.3 mL volume per well), PCR tubes with caps (0.2 mL), aluminum foil, and spectrophotometer/microplate reader. For all weighting operations an analytical balance with sensitivity of 0.0001 g was used. For boiling solutions in capped PCR tubes, a thermal cycler was used.

***Hazards***: Dinitrosalicylic acid (DNS) is corrosive and harmful if inhaled or swallowed. The prepared DNS reagent also contains 0.4 M sodium hydroxide, which is also an irritant. Phenol is also very hazardous in case of skin contact (corrosive, irritant), of eye contact (irritant), of ingestion, and of inhalation. The reduced DNS product (3-amino-5-nitrosalicylic acid) is toxic if swallowed and also causes eye irritation. Sugar containing buffers/solutions may cause eye and skin irritation. All reagents must be handled with care and properly disposed. Researchers must wear laboratory gloves and eye protection glasses at all times. Boiling hot water (or hot plates) can cause burns. Use heat resistant gloves when handling tubes immersed in a hot water bath or hot plate (or PCR thermal cycler).

***Stock Solution Preparation Experimental Procedure***:
1. Wear suitable personal protection equipment (PPE) before starting experiment: Gloves, Eye-Goggles, and Lab coat.
2. Prepare DNS assay stock reagent as explained below.

a. Dissolve 5.3 g of 3,5-dinitrosalicyclic acid (DNS) and 9.9 g sodium hydroxide (NaOH) in 708 mL of deionized water in a 1-L volumetric flask. Dissolve completely with the aid of an added magnetic stirrer bar, if available.
b. Next, add 153 g of sodium potassium tartarate, 4.2 g sodium meta-bisphosphate, and 3.8 mL phenol to the solution and dissolve completely. Add the phenol last. Use of phenol in the DNS reagent is to intensity the color but can be avoided if needed to minimize handling of hazardous reagent.
c. Transfer solution to a 1-L glass bottle and cover the bottle completely with aluminum foil to prevent any light from entering it. Store at 4$^o$ C, if available.

3. Prepare stock sugar standards for regular assays, as explained below.
    a. Glucose Stock Solution: Add 1.000 g glucose to 50 mL of water in 100 mL volumetric flask. Plug flask and invert to dissolve. Make up volume to 100 mL with addition of deionized water and mix well to fully dissolve glucose to prepare 10 g/L stock solution.
    b. Next, prepare 0 g/L (blank deionized or DI water), 1 g/L, 2 g/L, 3 g/L, 5 g/L glucose standards by preparing suitable dilutions (in deionized water) of original 10 g/L stock in 50 mL plastic conical Falcon tubes. Additional lower concentration glucose standards ranging from 0.1-1 g/L can be prepared as well if necessary.
    c. Label all glucose standards tubes and store at 4$^o$ C, if fridge is available.
    d. Note: Prepare sugar stock sugar standards for samples neutralized with NaOH following sulfuric acid hydrolysis (see SI-IV detailed experimental procedure below for details). Include neutralized $H_2SO_4$ in the glucose standard solution when evaluating sugar concentrations following acid hydrolysis to account for possible interference by neutralized salts on sugar detection.

***Detailed Experimental Procedure***:

1. Pipette 30 $\mu$L of unknown sample solution (or glucose standards provided by instructor) and 60 $\mu$l of orange DNS reagent into a small PCR tube. Perform all assays in duplicates.
2. Vortex tubes to mix solutions thoroughly.
3. Heat tubes at 95°C for 5 minutes (use thermal cycler, hot plate, or water bath, if available) and let samples cool back to room temperature for 15 min afterwards.
    a. Increasing reddish brown color intensity will indicate corresponding increasing concentration of total reducing sugars present in solution. See image below for characteristic colors seen for the glucose standards (starting with water blank on left followed by glucose standards with increasing concentration from 1 to 5 g/L). A light-yellow color indicates low sugar concentration while a dark brown or reddish black color indicates higher sugar concentration.

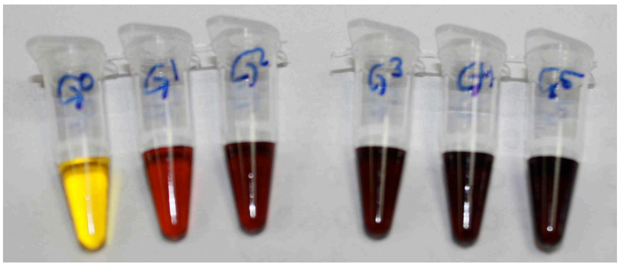

b. Measure the absorbance of each known-glucose standard (0 g/L to 5 g/L) to create a linear calibration curve from which to compare samples of unknown sugar concentrations based on absorbance values alone.

4. Use a 96-well clear round-bottom plate and add 178 μL deionized $H_2O$ to each well. Next, add 17.8 μL of each sample post the 5 min reaction (from step 3 above). Mix solutions by aspirating with pipette. See image below of dilute samples ready for absorbance measurement.

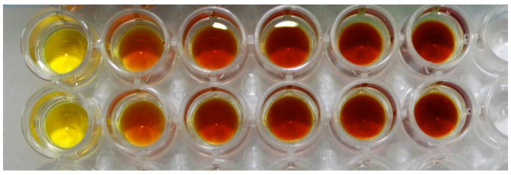

*Yellow solutions in wells on left side of image are 0 g/L of glucose while darkest wells on the right side increase progressively to 5 g/L of glucose. Glucose standard samples have been duplicated (top vs. bottom row). Unknown concentration samples can be included on a separate row or column as instructed by the teacher.*

5. Using a microplate-reader based UV-Vis spectrophotometer, measure the absorbance of each well volume using 540 nm wavelength light. Cuvette (0.1-1 mL) based UV-Vis spectrophotometers can also be used but the user will need to scale up the reaction volumes by at least 10 folds. See image below showing screenshot of excel file used to create calibration curve for standards using absorbance readings.

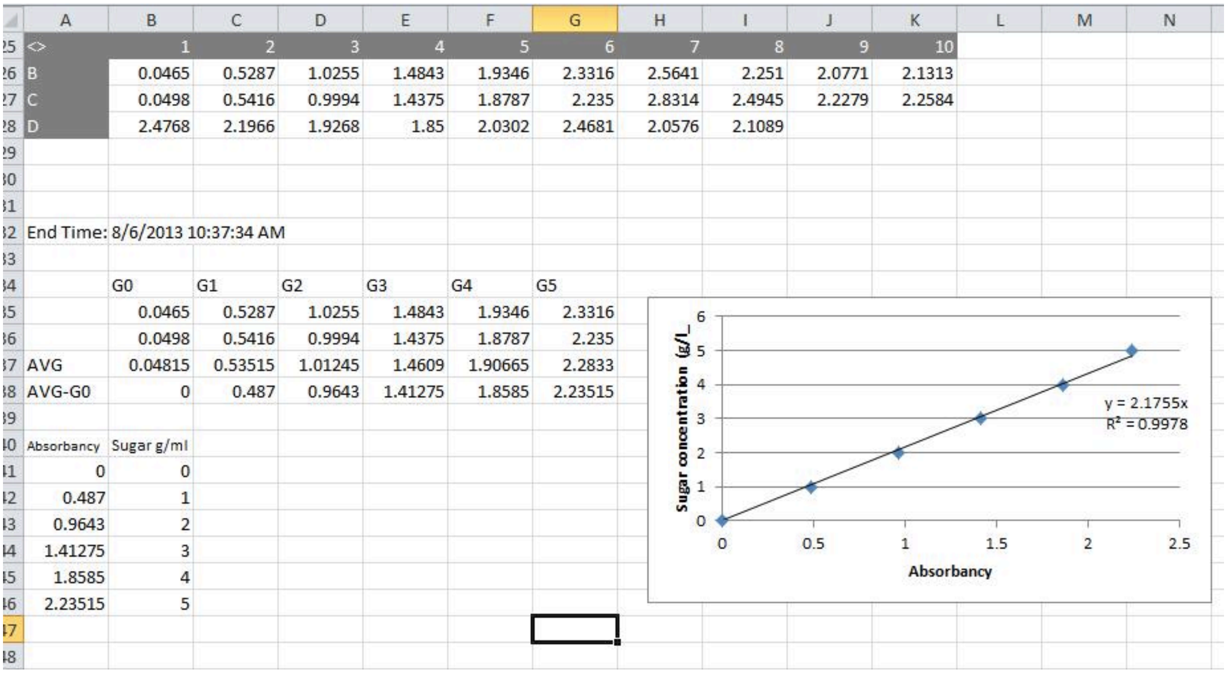

| <> | 1 | 2 | 3 | 4 | 5 | 6 | 7 | 8 | 9 | 10 |
|---|---|---|---|---|---|---|---|---|---|---|
| B | 0.0465 | 0.5287 | 1.0255 | 1.4843 | 1.9346 | 2.3316 | 2.5641 | 2.251 | 2.0771 | 2.1313 |
| C | 0.0498 | 0.5416 | 0.9994 | 1.4375 | 1.8787 | 2.235 | 2.8314 | 2.4945 | 2.2279 | 2.2584 |
| D | 2.4768 | 2.1966 | 1.9268 | 1.85 | 2.0302 | 2.4681 | 2.0576 | 2.1089 | | |

End Time: 8/6/2013 10:37:34 AM

| | G0 | G1 | G2 | G3 | G4 | G5 |
|---|---|---|---|---|---|---|
| | 0.0465 | 0.5287 | 1.0255 | 1.4843 | 1.9346 | 2.3316 |
| | 0.0498 | 0.5416 | 0.9994 | 1.4375 | 1.8787 | 2.235 |
| AVG | 0.04815 | 0.53515 | 1.01245 | 1.4609 | 1.90665 | 2.2833 |
| AVG-G0 | 0 | 0.487 | 0.9643 | 1.41275 | 1.8585 | 2.23515 |

| Absorbancy | Sugar g/ml |
|---|---|
| 0 | 0 |
| 0.487 | 1 |
| 0.9643 | 2 |
| 1.41275 | 3 |
| 1.8585 | 4 |
| 2.23515 | 5 |

6. Glucose standards results are averaged for duplicate samples and adjusted by subtracting the absorbance of the 0 g/L blank water sample from all other standards and unknown samples. Create a scatter plot of blank subtracted absorbance to known standard reducing sugar concentration to generate a calibration curve after linear-regression curve fitting.

7. Find average corrected absorbance for each unknown sample (make sure to subtract the 0 g/L blank from each reading). Use the slope of the calibration curve line to determine

the sugar concentration by multiplying unknown corrected sample absorbance by slope of the best-fit line. See image below showing screenshot of excel file used to determine unknown reducing sugar concentration using calibration curve.

8. At the end of the experiment, all waste reagents are to be carefully discarded in the waste solution container.

| | | | | | | |
|---|---|---|---|---|---|---|
| | G0 | G1 | G2 | G3 | G4 | G5 |
| | 0.0465 | 0.5287 | 1.0255 | 1.4843 | 1.9346 | 2.3316 |
| | 0.0498 | 0.5416 | 0.9994 | 1.4375 | 1.8787 | 2.235 |
| AVG | 0.04815 | 0.53515 | 1.01245 | 1.4609 | 1.90665 | 2.2833 |
| AVG-G0 | 0 | 0.487 | 0.9643 | 1.41275 | 1.8585 | 2.23515 |
| Absorbancy | Sugar g/ml | | 1 | 2 | 3 | 4 |
| 0 | 0 | | Glucose | Glucose ' | Aspen | Aspen ' |
| 0.487 | 1 | | 2.5641 | 2.251 | 2.0771 | 2.1313 |
| 0.9643 | 2 | | 2.8314 | 2.4945 | 2.2279 | 2.2584 |
| 1.41275 | 3 | | 2.4768 | 2.1966 | 1.9268 | 1.85 |
| 1.8585 | 4 | | 2.0302 | 2.4681 | 2.0576 | 2.1089 |
| 2.23515 | 5 | | | | | |
| | | AVG | 2.48 | 2.35 | 2.07 | 2.09 |
| | | AVG-G0 | 2.43 | 2.30 | 2.02 | 2.04 |
| | | Sugar g/ml | 5.28 | 5.01 | 4.40 | 4.44 |

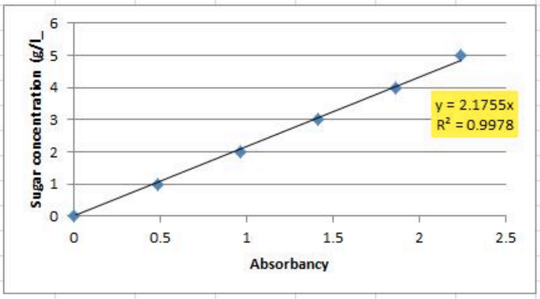

#### (SI-II) Glucose vs. Reducing Sugars Estimation Using Glucometers Supporting Information

***Summary and Principle***: Commercial blood glucose meters are available in standard pharmacy stores over-the-counter for diabetic patients to track fluctuations in their blood glucose level. Various types of blood glucose meters are available in the market that vary in the type of detection chemistry and/or sensitivity for monitoring soluble sugars like glucose etc. However, most diabetes kits come equipped with a monitoring test strip and a digital meter to read the test strip. The meter is a digital device that uses electrical energy to determine how much sugar or glucose is present in the blood sample. Here we evaluate the suitability of using two distinct test strips from Trividia Health (TRUEtest and TRUEbalance) with varying electrochemical detection methods to test their suitability for this biofuels lab protocol. Note that the TRUEtest test strip amperometrically estimates reducing sugars utilizing a non-specific *Acinetobacter calcoaceticus* derived glucose dehydrogenase-pyrroloquinoline quinone co-factor based immobilized enzymatic reaction. While the TRUEbalance test strip amperometrically estimates glucose only utilizing a highly specific *Aspergillus* sp. derived glucose oxidase based immobilized enzymatic reaction. Both test strip reactions are accompanied by a change in the digital reading, with sugar samples containing more reducing sugars giving a higher response. However, since these test strips and glucose meters have been optimized for estimating the concentration of either reducing sugars or glucose in blood plasma, suitable calibration curves will need to be prepared to use these kits for sugar solutions from non-blood plasma based sources. Plotting the digital output reading for known standard sugar concentrations (e.g., xylose dissolved in water) will allow generation of standard curves that can be used to determine the unknown concentrations of reducing sugars in desired samples.

***List of Chemicals and Materials for Experiment***: TRUEtest and TRUEbalance strips and Trividia glucose meters are available from Trividia Health (http://www.trividiahealth.com). Glucose, xylose, mannose, galactose, arabinose and cellobiose were all procured from Fisher-Scientific, Sigma Aldrich, or VWR to be used as is. Distilled water was used in the preparation of all aqueous solutions (unless specified otherwise).

All necessary glassware, lab equipment, and supplies were used to carry out all experiments are highlighted here: Single/multi-channel micropipettes and associated plastic tips (10, 100, 1000 μL range), Glass pipette (1-10 mL), Falcon conical plastic tubes (15, 50 mL range), Volumetric flask (100, 1000 mL), 96-well clear flat-bottom microplates (0.3 mL volume per well), PCR tubes with caps (0.2 mL), aluminum foil, and spectrophotometer/microplate reader. For all weighting operations an analytical balance with sensitivity of 0.0001 g was used.

***Hazards***: Diabetes meter test strips are not to be consumed and are also not supposed to be used by students for testing sugar concentrations in any other bodily fluids. Sugar containing buffers/solutions may cause eye and skin irritation. All reagents must be handled with care and properly disposed. Researchers must wear laboratory gloves and eye protection glasses at all times.

***Stock Solution Preparation Experimental Procedure***:

1. Wear suitable personal protection equipment (PPE) before starting experiment: Gloves, Eye-Goggles, and Lab coat.
2. Prepare stock sugar standards for regular assays, as explained below.
    a. Glucose Stock Solution: Add 1.000 g glucose to 50 mL of water in 100 mL volumetric flask. Plug flask and invert to dissolve. Make up volume to 100 mL with addition of deionized water and mix well to fully dissolve glucose to prepare 10 g/L stock solution.
    b. Next, prepare water blank (0 g/L control) and glucose standards (1 g/L, 2 g/L, 3 g/L, and 5 g/L) by preparing suitable dilutions (in deionized water) of original 10 g/L stock in 50 ml plastic conical Falcon tubes. Additional glucose standards of varying concentrations can be prepared as well, if necessary, by serial dilutions.
    c. Label all glucose standards tubes and store at 4$^{o}$ C, if fridge is available.
    d. Note: Prepare sugar stock sugar standards for samples neutralized with NaOH following sulfuric acid hydrolysis (see SI-IV detailed experimental procedure below for details). Include neutralized $H_2SO_4$ in the glucose standard solution when evaluating sugar concentrations following acid hydrolysis to account for possible interference by neutralized salts on sugar detection.
3. Prepare other sugar (Xylose, mannose, galactose, arabinose and cellobiose) standards using similar protocol outlined above.

***Detailed Experimental Procedure***:

1. Obtain a test strip from the diabetes test strip kit bottle and immediately recap bottle after use (to prevent exposure to air).
2. Insert the test strip into glucose meter (as shown in image below), and the meter will turn on automatically. Be careful to not damage the test strip.

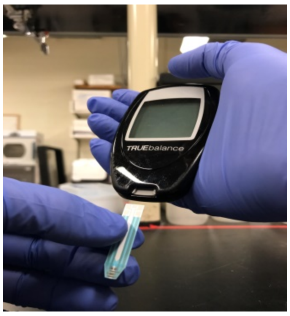

*Insert strip into test meter*

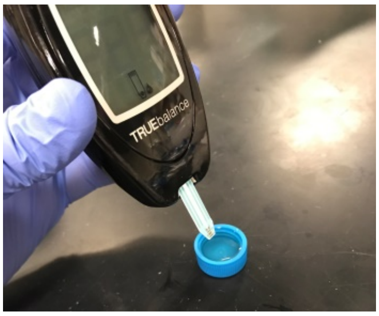

*Dip test strip into sugar solution*

3. Pipette ~10-20 μL of sample on to bottom edge of the test strip. Include all glucose or sugar standards as samples to prepare calibration curve. <u>OR</u> insert the test strip into sample solution and immediately remove the meter/tip from the tube cap once the meter beeps. See image above.
4. Wait ten seconds, and then read display to find your sample's estimated sugar concentration given in mg/dL concentration units.
5. Suitable sugar stock standards should be analyzed using this protocol and results can be averaged for duplicate samples to confirm the linearity of the scatter plot of displayed sugar or glucose concentration to known standard glucose sugar concentration after linear-regression curve fitting (as highlighted in SI-I protocol earlier).

<u>**Sensitivity of test strips for analysis of cellulosic biomass-derived sugars:**</u>

Here the authors have outlined some key results that were obtained that justify use of either the TRUEtest or TRUEbalance test strips to quantify different monosaccharide and disaccharide sugars derived from cellulosic biomass under different experimental scenarios. The instructor can use reported results to setup appropriate experiments and challenge students with relevant charge questions.

***A. TRUEtest Strips Reproducibility and Sensitivity***: The TRUEtest glucose test strips were used with the TRUEresult meter to measure glucose concentration in known glucose-only containing standards (prepared in deionized water alone). Two known standards were tested by pipetting 10-20 μL solution onto the tip of the test strip. The data shown in table below suggests that these test strips are capable of reading glucose concentrations within ~10-20% standard error of reported mean value. Total number of replicates measured here were 3-4. The values reported on the meter display were also close to the actual values (within 20% of actual value).

| Actual Conc. (mg/dL) | Meter Readings (mg/dL) | Average | Std Dev | % Std Error |
|---|---|---|---|---|
| 56.25 | 64 | 51 | 12 | 24 |
| | 48 | | | |
| | 40 | | | |
| 112.5 | 90 | 89 | 2 | 2 |
| | 87 | | | |

In addition, separate stock sugar solutions were made of xylose, mannose, galactose, arabinose and cellobiose by dissolving 50 mg of respective sugar in 10 mL deionized water (and diluted by 10-fold for some sugar solutions). These sugar solutions were then characterized using the same method as earlier to check for false glucose responses and checking the sensitivity of the TRUEtest immobilized enzymes for various plant biomass relevant sugars. Unfortunately, all tested five sugars also gave varying false responses for "glucose", therefore if any of these sugars are present in a sample, the glucose concentration will either likely be under or over-estimated (depending on sugar type; see results in table below where N.D. stands for not determined).

| Concentration A | | Concentration B (if needed) | |
|---|---|---|---|
| 500 mg/dL | Meter Reading | 50 mg/dL | Meter Reading |
| xylose | outside range | xylose | 64 |
| | outside range | | 65 |
| mannose | 251 | | N.D. |
| | 259 | | |
| galactose | 325 | | N.D. |
| | 264 | | |
| arabinose | outside range | arabinose | 57 |
| | outside range | | 65 |
| cellobiose | 423 | | N.D. |
| | 313 | | |

Therefore, these test strips cannot be used if other sugars are present in the sample solution but can be used to give a rough estimation of glucose if and only if glucose alone is present in the samples. In summary, the TRUEtest test strip is highly non-specific towards reducing sugars like glucose similar to the non-specific DNS assay type colorimetric methods.

***B. TRUEbalance Strips Reproducibility and Sensitivity***: The TRUEbalance blood glucose test strips were used with the TRUEbalance meter to measure the glucose concentration in known glucose-only standards. Two known standards were tested by pipetting 10-20 μL solution onto the tip of the test strip. The data shown in table below suggests that these test strips are capable of reading glucose concentrations within narrower ~1-2% standard error of reported mean value unlike TRUEtest strips. Total number of replicates was at least 3. However, the values reported on the meter display were consistently higher than the actual values (by ~2.5-3-fold of actual value). Therefore, a correction factor was determined using a calibration curve (shown below) to correct for the over prediction of glucose in solutions in the linear range of 50-200 mg/dL of known glucose concentrations in water.

| Actual Conc. (mg/dL) | Meter Readings (mg/dL) | Average | Std Dev | % Std Error |
|---|---|---|---|---|
| 50 | 161 | 158 | 3 | 2 |
| | 157 | | | |
| | 156 | | | |
| 100 | 324 | 330 | 7 | 2 |
| | 328 | | | |
| | 338 | | | |
| 200 | 528 | 526 | 2 | 0 |
| | 526 | | | |
| | 524 | | | |

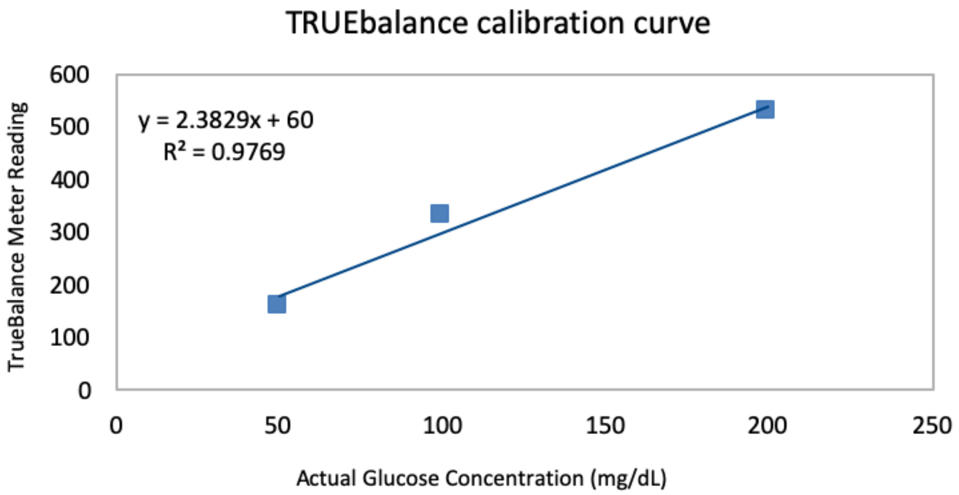

In addition, separate stock sugar solutions were also prepared for xylose, mannose, galactose, arabinose, and cellobiose by dissolving 50 mg of respective sugar in 10 mL deionized water. These sugar solutions were then also measured using the same method as earlier to check for false "glucose" responses and checking the sensitivity of the

TRUEbalance immobilized enzymes for various plant biomass relevant sugars. Fortunately, none of the other five sugars gave any significant false response for "glucose", so if any of these sugars are present in a sample, the test strip will give no response (or give error message for samples being below the detection range) due to high sensitivity of the immobilized enzyme towards glucose alone (see response tabulated below). Therefore, these test strips can be used if other sugars are present to give a rough estimation of glucose present in complex sugar composition samples. In principle, the TRUEbalance test strip is highly specific towards glucose alone like the spectrophotometric assay employing glucose hexokinase and glucose-6-phosphate dehydrogenase (See SI-III protocol for details). These test strips can be used to quickly identify glucose concentration between 50-200 mg/dL in the presence of other sugars as long as suitable glucose standards are used in the exact same buffer conditions to estimate a correction factor for over-prediction of glucose concentrations.

| **Concentration A** | |
|---|---|
| **500 mg/dL** | **Meter Reading** |
| xylose | below range |
| | below range |
| mannose | below range |
| | below range |
| galactose | below range |
| | below range |
| arabinose | below range |
| | below range |
| cellobiose | below range |
| | below range |

***C. Impact of sodium sulfate on TRUEbalance vs. TRUEtest Strips Response***: Both test strips were tested to see if there was any interference effect due to presence of high concentrations of sodium sulfate ($Na_2SO_4$) salt on electrochemical detection of suitable sugars. Sodium sulfate is a major byproduct formed during NaOH catalyzed neutralization of sulfuric acid containing hydrolysis samples that is useful for estimation total insoluble cellulose/hemicellulose composition of biomass samples (See SI-IV protocol for details). A known glucose stock of 225 mg/dL was spiked with an equal volume of water and compared to a sample that was spiked with an equal volume of 1M $Na_2SO_4$. The TRUEtest strips showed no effect due to inclusion of $Na_2SO_4$ whereas the TRUEbalance strips showed about a 33% decrease in the glucose concentration reading. In conclusion, as long as the sugar standards that are used to make the TRUEbalance calibration curve are in the same buffer conditions as the samples to be measured, the TRUEbalance strips can still be used to accurately measure glucose concentration.

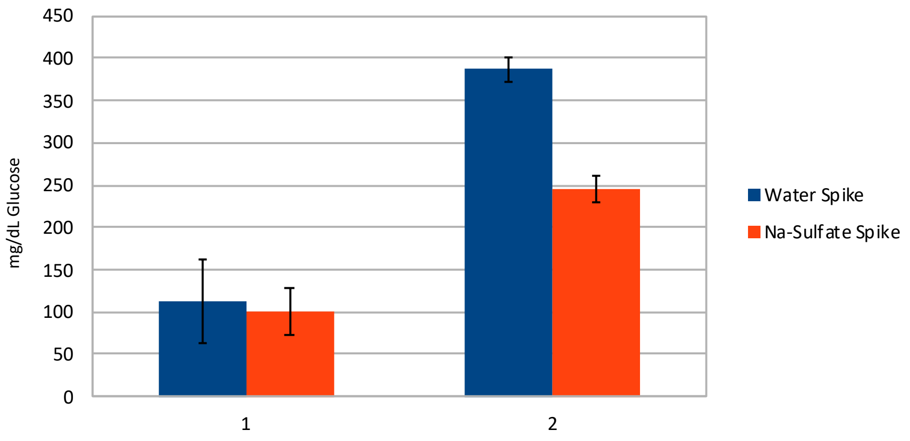

#### (SI-III) Enzyme Kit Based Glucose Estimation Supporting Information

***Summary and Principle***: Commercial enzyme based glucose detection assays are available via various vendors for detection/quantification of glucose. Here we evaluate an enzyme-based glucose detection/quantification kit available commercially from Catachem (Product Catalog No. C124-12; http://www.catacheminc.com/). The glucose assay was based on two-step enzymatic reactions, where D-Glucose was first phosphorylated to D-glucose-6-phosphate (G6P) using ATP and a glucose-specific Hexokinase enzyme. The G6P was then reacted with oxidized nicotinamide adenine dinucleotide (NADP+) by glucose-6-phosphate dehydrogenase to form D-gluconate-6-phosphate and reduced NADPH. The reactions are stoichiometric to the amount of D-glucose and the corresponding increase in NADPH concentration was measured at 340 nm to estimate glucose concentration based on a suitable standard curve. Alternative enzyme based glucose assay kits can be procured and tested from Megazyme (Bray, Ireland) and R-Biopharm (Marshall, Michigan), if necessary. Furthermore, for analogous accurate estimation of xylose and arabinose derived xylan and arabinan polymer fractions, one could use other sugar monosaccharides specific enzyme based kits available from other commercial vendors (like Megazyme) to estimate the hemicellulose mass balance more accurately.

Here we evaluate the suitability of the Catachem glucose enzyme assay alone to test its suitability for this biofuels lab protocol. As before, since these assay methods have been optimized for estimating the concentration of either reducing sugars or glucose in blood plasma, suitable calibration curves will need to be prepared to use these kits for sugar solutions from non-blood plasma based sources. Plotting the UV absorbance readings for known standard sugar concentrations (e.g., glucose dissolved in water) will allow generation of standard curves that can be used to determine the unknown concentrations of reducing sugars in desired samples. This protocol was developed based on vendor supplied instruction manual and a microplate version of a similar protocol.[2,3]

***List of Chemicals and Materials for Experiment***: Catachem Glucose Enzyme Assay Kit (Product Catalog No. C124-12; http://www.catacheminc.com/). Glucose, xylose, mannose, galactose, arabinose and cellobiose were all procured from Fisher-Scientific, Sigma Aldrich, or VWR to be used as is. Distilled water was used in the preparation of all aqueous solutions (unless specified otherwise). All necessary glassware, lab equipment, and supplies were used to carry out all experiments are highlighted here: Single/multi-channel micropipettes and associated plastic tips (10, 100, 1000 μl range), Glass pipette (1-10 mL), Falcon conical plastic tubes (15, 50 mL range), Volumetric flask (100, 1000 mL), 96-well clear flat-bottom microplates (0.3 mL volume per well), PCR tubes with caps (0.2 mL), aluminum foil, and spectrophotometer/microplate reader. For all weighting operations an analytical balance with sensitivity of 0.0001 g was used. For warming solutions, in capped PCR tubes or sealed microplates, a thermal cycler or temperature-controlled oven or water bath can be used.

***Hazards***: Enzyme kit reagents solutions are not to be consumed and are also not supposed to be used by students for testing sugar concentrations in any other bodily fluids. Enzyme/sugar containing buffers/solutions may cause eye and skin irritation. All reagents must be handled with care and properly disposed. Researchers must wear laboratory gloves and eye protection glasses at all times.

***Stock Solution Preparation Experimental Procedure***:

1. Wear suitable personal protection equipment (PPE) before starting experiment: Gloves, Eye-Goggles, and Lab coat.
2. Prepare stock sugar standards for regular assays, as explained below.
    a. Glucose Stock Solution: Add 1.000 g glucose to 50 mL of water in 100 mL volumetric flask. Plug flask and invert to dissolve. Make up volume to 100 mL with addition of deionized water and mix well to fully dissolve glucose to prepare 10 g/L stock solution.
    b. Next, prepare water blank (0 g/L control) and glucose standards (1 g/L, 2 g/L, 3 g/L, and 5 g/L) by preparing suitable dilutions (in deionized water) of original 10 g/L stock in 50 ml plastic conical Falcon tubes. Additional glucose standards of varying concentrations can be prepared as well, if necessary, by serial dilutions.
    c. Label all glucose standards tubes and store at 4$^{o}$ C, if fridge is available.
    d. Note: Prepare sugar stock sugar standards for samples neutralized with NaOH following sulfuric acid hydrolysis (see SI-IV detailed experimental procedure below for details). Include neutralized $H_2SO_4$ in the glucose standard solution when evaluating sugar concentrations following acid hydrolysis to account for possible interference by neutralized salts on sugar detection.
3. Prepare other sugar (Xylose, mannose, galactose, arabinose and cellobiose) standards using similar protocol outlined above.

***Detailed Experimental Procedure***:

1. Pipette 500 μL of original stock HK Reagent (available in kit from Catachem) and 5 μL of sample containing sugars into small glass or plastic tubes. Include all glucose standards as samples to prepare calibration curve.
2. Gently aspirate to mix and incubate samples in oven at 37°C for 10 minutes.
3. Transfer 150 μL of incubated samples into a UV transparent cuvette or microplate.
4. Using a microplate-reader based spectrophotometer, measure absorbance of each sample using 340 nm UV light. Cuvette (0.1-1 mL) based spectrophotometers can be used but the user will need to scale up the reaction volumes by at least 10 folds.
5. Glucose standards results are averaged for duplicate samples and adjusted by subtracting the absorbance of 0 g/L blank from all standards and unknown samples. Create a scatter plot of absorbance to known standard glucose concentration to generate a linear calibration curve after curve fitting. See image below as example.

| | A | B | C | D | E | F | G | H | I | J | K | L | M | N | O |
|---|---|---|---|---|---|---|---|---|---|---|---|---|---|---|---|
| 26 | C | 0.1545 | 0.4597 | 0.7371 | 0.9815 | 1.2475 | 1.5196 | 1.7534 | 1.4858 | 1.0031 | 1.1169 | | | | |
| 27 | D | 0.1623 | 0.466 | 0.7472 | 0.9866 | 1.2772 | 1.5579 | 1.8024 | 1.5093 | 1.0171 | 1.1355 | | | | |
| 28 | E | 1.8061 | 1.4403 | 0.9978 | 1.1292 | 1.8427 | 1.4736 | 1.0474 | 1.1638 | | | | | | |
| 29 | | | | | | | | | | | | | | | |
| 30 | | | | | | | | | | | | | | | |
| 31 | | | | | | | | | | | | | | | |
| 32 | End Time: 8/6/2013 11:48:50 AM | | | | | | | | | | | | | | |
| 33 | | | | | | | | | | | | | | | |
| 34 | | G0 | G1 | G2 | G3 | G4 | G5 | | | | | | | | |
| 35 | | 0.1545 | 0.4597 | 0.7371 | 0.9815 | 1.2475 | 1.5196 | | | | | | | | |
| 36 | | 0.1623 | 0.466 | 0.7472 | 0.9866 | 1.2772 | 1.5579 | | | | | | | | |
| 37 | AVG | 0.1584 | 0.46285 | 0.74215 | 0.98405 | 1.26235 | 1.53875 | | | | | | | | |
| 38 | AVG-G0 | 0 | 0.30445 | 0.58375 | 0.82565 | 1.10395 | 1.38035 | | | | | | | | |
| 39 | | | | | | | | | | | | | | | |
| 40 | Absorbancy | Glu g/L | | | | | | | | | | | | | |
| 41 | 0 | 0 | | | | | | | | | | | | | |
| 42 | 0.30445 | 1 | | | | | | | | | | | | | |
| 43 | 0.58375 | 2 | | | | | | | | | | | | | |
| 44 | 0.82565 | 3 | | | | | | | | | | | | | |
| 45 | 1.10395 | 4 | | | | | | | | | | | | | |
| 46 | 1.38035 | 5 | | | | | | | | | | | | | |
| 47 | | | | | | | | | | | | | | | |
| 48 | | | | | | | | | | | | | | | |
| 49 | | | | | | | | | | | | | | | |

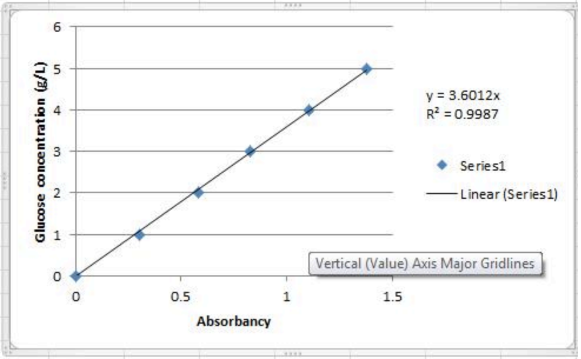

6. Find average corrected absorbance for each unknown sample (make sure to subtract the 0 g/L glucose blank from each unknown sample reading). Use the slope of the calibration slope line to determine the glucose concentration by multiplying unknown corrected sample absorbance by slope of the best-fit line. See image below showing screenshot of excel file used to determine unknown glucose sugar concentration using calibration curve.

| | | | | | | | | | | | |
|---|---|---|---|---|---|---|---|---|---|---|---|
| 25 | <> | 1 | 2 | 3 | 4 | 5 | 6 | 7 | 8 | 9 | 10 |
| 26 | C | 0.1545 | 0.4597 | 0.7371 | 0.9815 | 1.2475 | 1.5196 | 1.7534 | 1.4858 | 1.0031 | 1.1169 |
| 27 | D | 0.1623 | 0.466 | 0.7472 | 0.9866 | 1.2772 | 1.5579 | 1.8024 | 1.5093 | 1.0171 | 1.1355 |
| 28 | E | 1.8061 | 1.4403 | 0.9978 | 1.1292 | 1.8427 | 1.4736 | 1.0474 | 1.1638 | | |
| 29 | | | | | | | | | | | |
| 30 | End Time: 8/6/2013 11:48:50 AM | | | | | | | | | | |
| 31 | | | | | | | | | | | |
| 32 | | G0 | G1 | G2 | G3 | G4 | G5 | | | | |
| 33 | | 0.1545 | 0.4597 | 0.7371 | 0.9815 | 1.2475 | 1.5196 | | | | |
| 34 | | 0.1623 | 0.466 | 0.7472 | 0.9866 | 1.2772 | 1.5579 | | | | |
| 35 | AVG | 0.1584 | 0.46285 | 0.74215 | 0.98405 | 1.26235 | 1.53875 | | | | |
| 36 | AVG-G0 | 0 | 0.30445 | 0.58375 | 0.82565 | 1.10395 | 1.38035 | | | | |
| 37 | | | | | | | | | | | |
| 38 | Absorbancy | Glu g/L | | 1 | 2 | 3 | 4 | | | | |
| 39 | 0 | 0 | | Glucose | Glucose' | Aspen | Aspen' | | | | |
| 40 | 0.30445 | 1 | | 1.7534 | 1.4858 | 1.0031 | 1.1169 | | | | |
| 41 | 0.58375 | 2 | | 1.8024 | 1.5093 | 1.0171 | 1.1355 | | | | |
| 42 | 0.82565 | 3 | | 1.8061 | 1.4403 | 0.9978 | 1.1292 | | | | |
| 43 | 1.10395 | 4 | | 1.8427 | 1.4736 | 1.0474 | 1.1638 | | | | |
| 44 | 1.38035 | 5 | | | | | | | | | |
| 45 | | | AVG | 1.80 | 1.48 | 1.02 | 1.14 | | | | |
| 46 | | | AVG-G0 | 1.64 | 1.32 | 0.86 | 0.98 | | | | |
| 47 | | | Glu g/L | 5.92 | 4.75 | 3.09 | 3.52 | | | | |

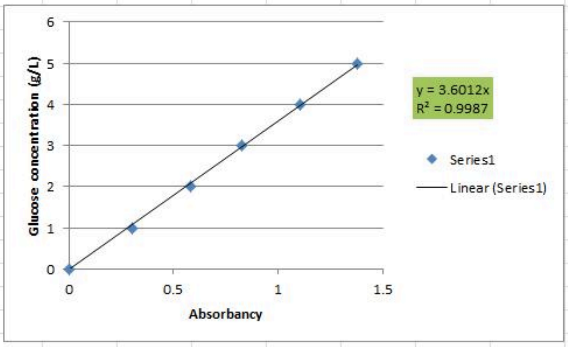

#### Sensitivity of test kit for analysis of cellulosic biomass-derived sugars:

Here the authors have outlined some key results that were obtained that justify use of the Catachem enzyme based assay kit to quantify glucose derived from acid hydrolyzed insoluble carbohydrate polymers from cellulosic biomass after NaOH catalyzed neutralization. The instructor can use reported results to setup appropriate experiments and challenge students with relevant charge questions.

***A. Impact of sodium sulfate on assay reproducibility and sensitivity***: Glucose Hexokinase and G6P-Dehydrogenase enzyme based glucose detection kit from Catachem (cat # C124-06) was tested for its sensitivity to detect glucose in the presence of high concentrations of $Na_2SO_4$. Unknown samples or glucose standards were tested by pipetting 10 μl of solution into 500 μL of the kit enzyme reagent in a 1 ml eppendorf plastic tube. After proper mixing, the tubes were capped and incubated in a 37 C oven for 10 minutes to ensure reaction kinetics endpoint is achieved. Next, 150 μL of each sample was added into a 96 well UV plate and the UV absorbance of the sample solution was measured at 340 nm. A standard curve was prepared as before with known concentrations of glucose against their respective absorbance values. The standard curve slope was used to calculate the glucose concentration of unknown samples based on absorbance readings (see image below).

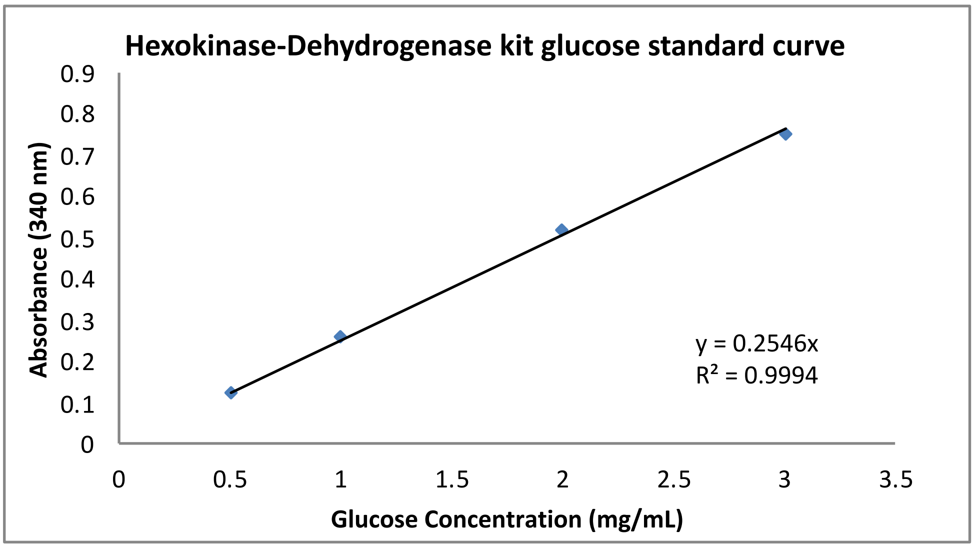

A stock glucose solution of 5 mg/mL was spiked with equal volumes of different concentrations of $Na_2SO_4$, with final concentrations of $Na_2SO_4$ in the sample volume ranging from 0 to 250 mM. These samples of known glucose concentration were then tested using the protocol detailed above to check if $Na_2SO_4$ had any effect on glucose detection. All of the calculated glucose concentrations were within 1.5% of the expected glucose concentrations (see image below). This result suggests that sodium sulfate does not have any significant impact on the hexokinase or G6P-Dehydrogenase enzyme activity to impact detection of glucose. In principle, this assay method is highly reproducible and specific towards glucose alone. All other sugars (e.g., cellobiose, mannose, galactose, xylose, arabinose) tested gave less than 0.5% response compared to glucose at equivalent 500 mg/dL or 5 mg/mL sugar concentrations (data not shown).

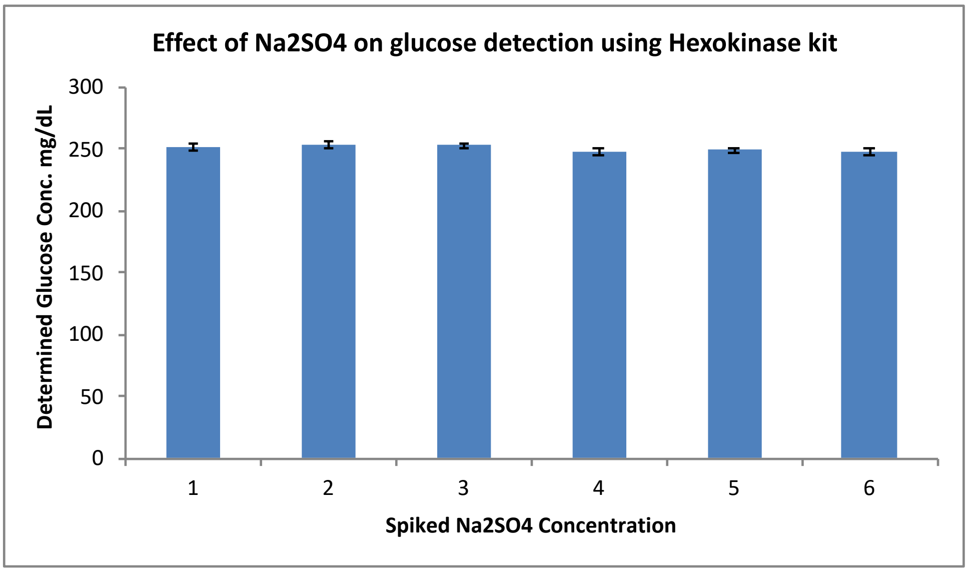

#### (SI-IV) Insoluble Biomass Composition Analysis Supporting Information

***Summary and Principle***: Insoluble polymeric carbohydrates (like β-1,4-glucan or cellulose and β-1,4-xylan which is a predominant hemicellulose in grasses) are either bound within the complex cell wall matrix of plant biomass or are released during biorefinery unit operations as high molecular weight oligosaccharides. Cellulose and hemicellulose content within samples can be estimated by hydrolyzing these polymers into their respective acid soluble monomeric forms (i.e., glucose and xylose) using concentrated/dilute sulfuric acid. A simple protocol was developed to estimate the composition of insoluble lignocellulosic biomass based on the compositional analysis protocols developed by the National Renewable Energy Laboratory (NREL) (https://www.nrel.gov/bioenergy/biomass-compositional-analysis.html). NREL recommends a two-step acid hydrolysis procedure to hydrolyze the carbohydrate polymers into easily quantifiable soluble sugars utilizing specialized equipment (e.g., pressurized vessels, autoclave, high pressure liquid chromatography systems) typically not available in schools and undergraduate laboratories. Our current method, inspired by another published study,[5] utilizes a multi-step acid hydrolysis procedure using a hot water-bath (and/or regular oven) alone and common chemistry lab apparatus to hydrolyze the insoluble carbohydrate polymers into soluble sugar monomers. The soluble sugars in the neutralized biomass hydrolyzate solution can then be assayed using the customized sugar assays described earlier to provide a comprehensive composition analysis for a diverse range of cellulosic feedstocks. The insoluble residue left behind after acid hydrolysis was predominantly enriched in lignin and was estimated gravimetrically after drying in the oven. Here, lignin will also fractionate into acid insoluble residue and acid soluble material. Acid soluble lignin can be measured using UV-Vis analysis with a background of 4% sulfuric acid. It was possible to achieve comparable cellulose and hemicellulose compositional results to what have been reported using the NREL protocols for comparable feedstocks. Please note that since ash analysis on the acid insoluble residue requires access to a specialized muffle furnace to burn off all organic carbon, this analysis was not carried out here. Therefore, the total acid insoluble lignin residue reported here is marginally higher and would like likely be skewed for biomass samples with high ash content. The current method is adapted from two other published studies to hydrolyze the insoluble carbohydrate polymers into soluble sugar monomers that are detected using previously outlined method (see protocols SI-I to III for details).[5,6] The following protocol can be used by the instructor specifically to prepare reagents for use by students as part of other learning activities OR can be used by students as part of a lab activity held in conjunction with other protocols outlined in this document.

***List of Chemicals and Materials for Experiment***: Glucose, xylose, mannose, galactose, arabinose, cellobiose, concentrated sulfuric acid, sodium sulfate, and sodium hydroxide were all procured from Fisher-Scientific, Sigma Aldrich, or VWR to be used as is. All lignocellulosic biomass substrates were a gift provided by the Great Lakes Bioenergy Research Center (GLBRC). Distilled water was used in the preparation of all aqueous solutions (unless specified otherwise). All necessary glassware, lab equipment, and supplies were used to carry out all experiments are highlighted here: Single/multi-channel micropipettes and associated plastic tips (10, 100, 1000 μL range), Glass pipette (1-10 mL), Glass tubes with plastic screw caps (10 mL), Falcon conical plastic tubes (15, 50 mL range), ceramic filtration crucibles, Volumetric flask (100, 1000 mL), round bottom flask (250 mL),

96-well clear flat-bottom microplates (0.3 mL volume per well), PCR tubes with caps (0.2 mL), aluminum foil, and spectrophotometer/microplate reader. For all weighting operations an analytical balance with sensitivity of 0.0001 g was used. For heating solutions or drying solids a hot water bath and temperature-controlled oven can be used, respectively. A standard reflux condenser setup (with attached chilled or room temperature water lines to condense reflux contents) should be available to boil dilute sulfuric acid aqueous solution at 95 Degrees Celsius using a heating mantle or boiling water bath. Buchner filtration flask-vacuum line assembly is needed to filter separate solids from liquids. A desiccator is needed to keep crucibles in prior to weighing dry weight on balance.

***Hazards***: Sulfuric acid is corrosive and harmful if inhaled or swallowed. The neutralization agent (sodium hydroxide) is also an irritant. Sugar containing buffers/solutions may cause eye and skin irritation. Enzyme kit reagents solutions are not to be consumed and are also not supposed to be used by students for testing sugar concentrations in any other bodily fluids. Enzyme/sugar containing buffers/solutions may cause eye and skin irritation. All reagents must be handled with care and properly disposed. Researchers must wear laboratory gloves and eye protection glasses at all times. Boiling hot water (or hot ovens) can cause burns. Use heat resistant gloves when handling tubes immersed in a hot water bath or hot oven.

***Stock Solution Preparation Experimental Procedure***:

1. Wear suitable personal protection equipment (PPE) before starting experiment: Gloves, Eye-Goggles, and Lab coat.
2. Prepare stock sugar standards for regular assays, as explained below.
    a. Glucose Stock Solution: Add 1.000 g glucose to 50 mL of water in 100 mL volumetric flask. Plug flask and invert to dissolve. Make up volume to 100 mL with addition of deionized water and mix well to fully dissolve glucose to prepare 10 g/L stock solution.
    b. Next, prepare water blank (0 g/L control) and glucose standards (1 g/L, 2 g/L, 3 g/L, and 5 g/L) by preparing suitable dilutions (in deionized water) of original 10 g/L stock in 50 ml plastic conical Falcon tubes. Additional glucose standards of varying concentrations can be prepared as well, if necessary, by serial dilutions.
    c. Label all glucose standards tubes and store at $4^{o}$ C, if fridge is available.
    d. Note: Prepare sugar stock sugar standards for samples neutralized with NaOH following sulfuric acid hydrolysis (see SI-IV detailed experimental procedure below for details). Include neutralized $H_2SO_4$ in the glucose standard solution when evaluating sugar concentrations following acid hydrolysis to account for possible interference by neutralized salts on sugar detection.
3. Prepare other sugar (Xylose, mannose, galactose, arabinose and cellobiose) standards using similar protocol outlined above.
4. Prepare 25% and 77% $H_2SO_4$ solutions (by weight; where Specific Gravity is 1.84 at 25 Degrees Celsius) from original concentrated sulfuric acid stock. Handle stock solution with care using appropriate PPE and store prepared diluted acid solutions in suitable capped glass bottles for long-term storage.
5. Prepare 10 M NaOH stock solution and store in capped glass bottles for long-term storage.
6. Refer to appropriate protocols (see SI-I to SI-III) for detection of released soluble sugars.

7. Determine total moisture content of each biomass sample by drying in oven set at 100°C, ideally overnight, to remove any residual moisture till constant dry weight.

***Detailed Experimental Procedure***:

1. Weigh out 100 mg of each lignocellulosic biomass sample to be analyzed. Include a 100 mg sample of glucose as sugar recovery standard. Write out mass of each biomass as precisely as possible (to the hundredth of a mg).
2. Transfer 100 mg of biomass to small 5-10 mL glass capped vials. See image below. Here in the image below, the respectively labeled tubes contain; (1) glucose recovery standard, (2) cardboard, (3) switchgrass, (4) corn stover, and (5) aspen wood.

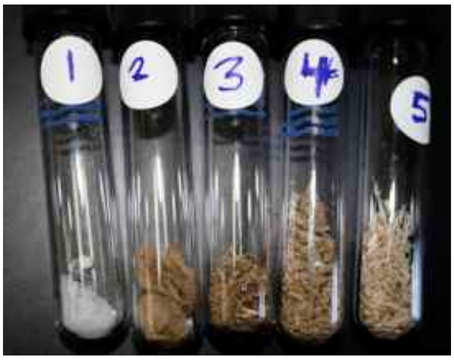

3. Carefully add 1 mL of 77% $H_2SO_4$ solution to each biomass sample using a glass pipette, and continue stirring with a small capillary tube or glass rod to ensure complete immersion of biomass into acid solution. Then, cap the tubes and allow the samples to hydrolyze in this concentrated acid solution at room temperature (25 Degrees Celsius) for 90 minutes with occasional mixing every 15 mins. See image below of representative samples after 90 mins incubation period.

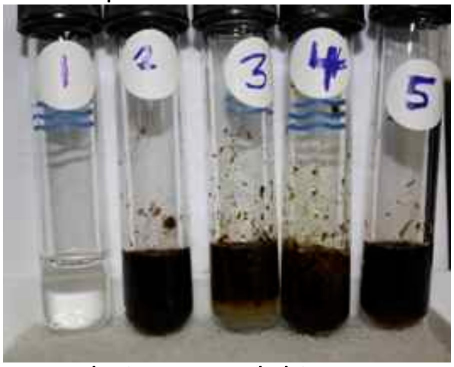

4. Next, add 1 mL of 25% $H_2SO_4$ solution to each biomass sample after 90 mins, and again stirring with a small capillary tube or glass rod to ensure complete immersion of biomass samples with limited sticking to tube walls. Cap the tubes and allow the samples to hydrolyze in this dilute acid solution in an incubator or water bath set at 55°C for 2 hours total hydrolysis time. See image below of representative samples after 2 hrs incubation period.

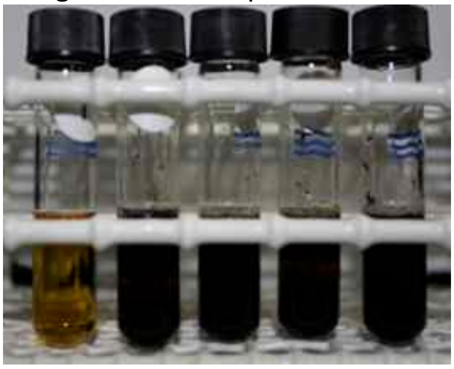

5. Use 10 mL of chilled deionized water to then quantitatively transfer contents of the capped tubes into 250 mL round bottom reflux flasks, ensuring that all contents from tubes are fully transferred. The total volume in round bottom flask should be ~12 mL (10 mL of $H_2O$ + 2 mL $H_2SO_4$ + some insoluble biomass residues). The flasks with the biomass immersed in very dilute acid should be then heated at 95°C boiling water bath or heating mantle for 45 minutes in a reflux setup. See representative images below.

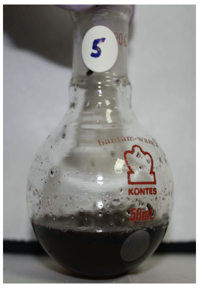

*Sample transferred to reflux flask*

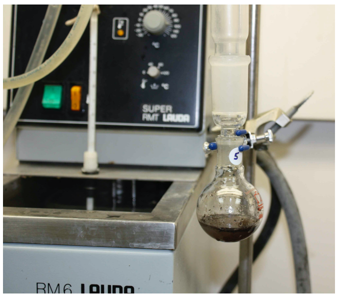

*Setup Reflux apparatus*

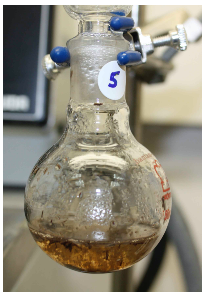

*After 45 minutes reflux*

6. The flask containing glucose control should be neutralized using 10 M NaOH solution (it should take approximately ~2.9 mL of 10 M NaOH to neutralize the volume). Note the exact volume of liquid required to bring solution to neutral pH and write down total volume after transferring liquid to plastic 50 ml falcon tube or glass tube container. The total volume added would be important data needed during compositional analysis calculations as shown later in this protocol.
7. The biomass flasks containing solid particulates (mostly acid insoluble lignin) mixed with acid hydrolyzates are then to be filtered through a ceramic crucible fitted atop a Buchner funnel attached to vacuum suction line. In this example, cardboard, switchgrass, corn stover, and aspen wood all have non-hydrolyzed particles mixed with the liquid and therefore need to be filtered. See representative images below.

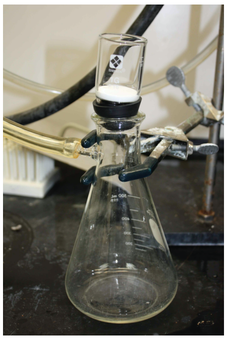

*Setup before filtering*

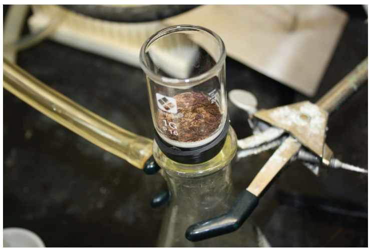

*Solids in crucible are mostly lignin*

8. Determine the mass of the dry crucible before using it for filtration and to eventually estimate dry weight of acid insoluble lignin after drying crucible.
9. For the biomass sample slurries that are filtered, obtain ~10 mL of liquid that poured through the funnel, this solution contains the soluble sugars that were hydrolyzed from insoluble polysaccharides into soluble monosaccharides by $H_2SO_4$. Neutralize this liquid using 10 M NaOH (it should take approximately 2.3-2.5 mL of 10 M NaOH). Note the exact volume of liquid required to bring solution to neutral pH and write down total

volume after transferring liquid into a plastic or glass tube container. The total volume added would be important data needed during compositional analysis calculations as shown later in this protocol.

10. After 10 mL of liquid has been removed, the solid particles left in the crucible should be washed with copious amounts of deionized $H_2O$ (~100-500 mL) in order to remove any soluble sugars and other water-soluble components. The vacuum suction should remove most of the liquid from the solid left in the crucible. To further dry the sample, dry the crucible in an oven set at 100°C for at least 1-2 hours, ideally overnight, to remove any residual moisture till constant dry weight. Afterwards, solid samples in crucibles should be placed in a desiccator for 1 hour as they cool. Finally, samples in crucibles should be weighed and compared to initial dry weight of the crucible before samples were filtered to determine amount of unhydrolyzed residue that is presumed here to be mostly acid-insoluble lignin.
11. Run appropriate soluble sugar assays (e.g., DNS, TRUEbalance, TRUEtest, and HK-G6PH sugar assays) on all sample hydrolyzates (see other relevant protocols).
12. Using results of DNS assay (i.e., total soluble reducing sugar concentration in each sample hydrolyzate), HK or TrueBalance assays (i.e., total glucose concentration in each sample hydrolyzate) and difference of crucible weight (i.e., total acid insoluble lignin), one can roughly estimate the composition of the starting biomass. Note that microcrystalline cellulose or Avicel based control biomass acid hydrolyzates (neutralized by base to pH 7.5) should give comparable cellulose compositions using either the DNS, TRUEBalance, or HK assays (within 10% error).
13. Calculate the composition of each biomass using the example worksheet below. For this example calculation worksheet, we will consider Glucose (as a recovery control), Aspen wood, Corn Stover, and Cardboard as the biomass samples analyzed using this protocol. Instructors can use this template to generate an excel worksheet with inserted formulas to simplify student data analysis as part of an extended laboratory protocol.

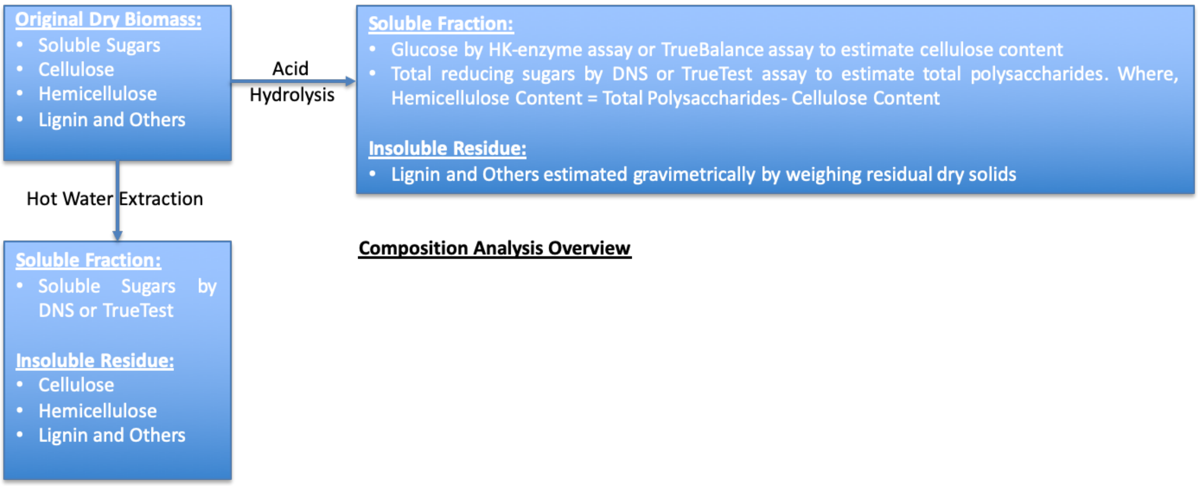

**Composition Analysis Overview**

| Biomass Type -> | Glucose Recovery Standard | Aspen Wood | Corn Stover | Cardboard | Explanation of Steps |
|---|---|---|---|---|---|
| **Part I: Acid Hydrolysis and DNS based Assay Relevant Calculations** | | | | | |
| Total weight of biomass (in glass vial) | 100.65 mg | 100.44 mg | 100.68 mg | 99.75 mg | See step 1 in protocol |
| % Moisture (in biomass on total weight basis) | 7.13% | 7.73% | 6.56% | 7.34% | Determined experimentally by oven drying till constant weight ahead of time by instructor |
| Dry weight of biomass (in glass vial) | 93.47 mg | 92.68 mg | 94.08 mg | 92.43 mg | Subtract the total 'wet (i.e., water)' weight of biomass based on % moisture from the initial mass to determine dry mass. For example: glucose dry weight = 100.65-(100.65*.0713) |
| Total reducing sugar concentration | 6.20 g/L | 5.03 g/L | 4.17 g/L | 5.89 g/L | Determined from DNS assay for recovered biomass acid hydrolysate in Step 9 |
| Total volume of recovered biomass acid hydrolysate in Step 9 | 14.91 mL | 11.85 mL | 11.83 mL | 11.86 mL | For glucose: 2 ml of $H_2SO_4$ + 10 mL of $dH_20$ + volume of NaOH required to neutralize. For other biomasses: 10 mL of the {2 ml of $H_2SO_4$ + 10 mL of $dH_20$} solution is removed, so volume will be less (see protocol steps 6-11) |
| Absolute amount of total reducing sugar from starting biomass | 92.44 mg | 59.61 mg | 49.33 mg | 69.86 mg | Amount of sugar (mg) is calculated by multiplying concentration (g/L) by total volume (mL) |
| Corrected absolute amount of total reducing sugar | | 71.53 mg | 59.20 mg | 83.83 mg | Amount of total sugar (mg) should include 2 mL of residual volume left behind in flask and crucible in step 10. Assume concentration of sugar is the same. Using Aspen as an example: multiply 59.61 by 12/10 to give total sugar in sample |
| Equivalent amount of total poly-saccharide theoretically present in starting biomass | | 59.61 mg | 49.33 mg | 69.86 mg | To determine the equivalent amount of polysaccharide polymer from the mass of sugars (hydrolyzed by $H_2SO_4$) divide mass of sugars by 1.11 conversion factor, assuming total polysaccharides are predominantly composed of |

| | | | | | |
|---|---|---|---|---|---|
| | | | | | hexose-based sugars |
| % Glucose Standard Recovered | 98.9 | | | | Divide absolute amount of total reducing sugar in glucose recovery standard (calculated) by the dry initial mass to account for irreversible loss during acid hydrolysis. Subtract % recovered glucose from 100% to calculate %Lost. |
| % Glucose Standard Lost | 1.1 | | | | |
| Corrected total amount biomass poly-saccharide | | 60.27 mg | 49.88 mg | 70.64 mg | Assume % reducing sugars lost due to acid-catalyzed dehydration is same for all tested biomasses as it is for glucose. Multiply amount of polysaccharide by 100 and ÷ 98.9 |
| Final % Poly-saccharide content of biomass (dry weight basis) | | 65% | 53% | 76% | Divide amount of theoretical polysaccharide by dry initial weight of biomass |
| **Part II: Acid Hydrolysis and Hexokinase (HK-G6PDH) or TrueBalance based Glucose-specific Assay Relevant Calculations** | | | | | |
| Glucose concentration estimated in acid hydrolyzate | 6.03 g/L | 3.09 g/L | 2.01 g/L | 4.68 g/L | Value determined from HK-G6PDH or True Balance glucometer assay using appropriate standards based calibration curve (see method section for details) |
| Absolute amount of glucose from starting biomass | 89.91 mg | 36.62 mg | 23.78 mg | 55.50 mg | Amount of glucose (mg) is calculated by multiplying concentration (g/L) by total volume (mL) recovered after acid hydrolysis as before |
| Corrected absolute amount of glucose | | 43.94 mg | 28.54 mg | 66.60 mg | Same principle as disuccsed earlier using 12/10 correction factor. |
| Equivalent amount of total glucan polymers or cellulose theoretically present in starting biomass | | 39.59 mg | 25.71 mg | 60.0 mg | To determine the amount of cellulose from the mass of glucose (hydrolyzed by $H_2SO_4$) divide mass of glucose by 1.11 conversion factor, assuming total cellulose is composed of hexose-based sugars only |
| % Glucose Standard Recovered | 96.2% | | | | Divide absolute amount of total glucose in glucose recovery standard (calculated) by the dry initial mass to account for |
| % Glucose Standard | 3.8% | | | | |

| | | | | | |
|---|---|---|---|---|---|
| Lost | | | | | irreversible loss during acid hydrolysis. Subtract % recovered glucose from 100% to calculate %Lost. |
| Corrected total amount biomass cellulose | | 41.15 mg | 26.73 mg | 62.37 mg | Assume % glucose lost due to acid-catalyzed dehydration is same for all tested biomasses as it is for glucose standard. Multiply amount of cellulose by 100 and ÷ by 96.2 |
| Final % Cellulose content of biomass (dry weight basis) | | 44% | 28% | 67% | Divide amount of theoretical cellulose by dry initial weight of biomass |
| Final % Hemi-cellulose content of biomass (dry weight basis) | | 21% | 25% | 9% | Assuming % Total Polysaccharide - % cellulose = % Hemicellulose |
| Lignin absolute weight recovered from biomass after acid hydrolysis and drying | | 26.5 mg | 25.27 mg | 16.5 mg | Final mass of crucible – initial mass of crucible (see step 9 in protocol) |
| Final % Lignin content of biomass (dry weight basis) | | 29% | 27% | 18% | Any biomass mass not hydrolyzed by $H_2SO_4$ is presumed to be lignin, assuming near-complete hydrolysis of all poly-saccharides into sugars. Divide amount of lignin by dry initial mass. |

**Example Compositional Analysis Summary for various biomass substrates analyzed using proposed method (based on dry weight basis of starting material):**

| Aspen wood | | | Corn Stover | | | Cardboard | | |
|---|---|---|---|---|---|---|---|---|
| Cellulose | Hemicellulose | Lignin | Cellulose | Hemicellulose | Lignin | Cellulose | Hemicellulose | Lignin |
| 44% | 21% | 29% | 28% | 25% | 27% | 67% | 9% | 18% |

***Key assumptions about composition analysis method and all relevant calculations:***

- Here, we are firstly assuming all lignocellulosic biomasses are only composed lignin, cellulose, and hemicellulose based biopolymers.

- In addition, total soluble sugars (e.g., sucrose) if present in the initial biomass are estimated directly by DNS assay as well separately (as glucose equivalents) and values are added to the final composition analysis data. This could result in slight over-estimation of the total polysaccharide fraction if simple sugars like sucrose are present in high concentrations. One can estimate the water-soluble DNS sugars to adjust the total polysaccharide composition of the biomass but was not attempted here.
- We assume acid hydrolysis step results in complete hydrolysis of all polysaccharides into monosaccharides that can be measured using DNS or HK-G6PDH assay. This is one of the limitations of this protocol as we might be slightly over-estimating the total acid insoluble lignin content if the acid hydrolysis step leaves behind any insoluble polysaccharides residue.
- We assume acid hydrolysis step results in no significant solubilization of lignin, which means residual unhydrolyzed solids measured gravimetrically are composed entirely of lignin.
- Soluble glucose (estimated by HK-G6PDH method) to glucan or cellulose polymer is back-calculated based on stoichiometric equivalent conversion factor of 1.11 (i.e., 180 g/L glucose is equivalent to 162 g/L anhydroglucosyl units). For sake of simplicity, the same conversion factor is used to also convert total DNS method estimated reducing sugars to total polysaccharide amount. For more accurate estimation of xylose, and arabinose derived xylan, arabinan polymer fractions, one could use enzyme kits available from commercial vendors (like Megazyme) to close the hemicellulose mass balance more accurately.
- Glucose recovery standard alone is used to account for any undesired loss of monosaccharides to acid-catalyzed dehydration products like furans. This is one of the limitations of this protocol as we should be using pentose sugar standards (e.g., xylose) when estimating hemicellulose fraction. However, since hemicellulose fraction is indirectly estimated by subtracting cellulose fraction from total polysaccharide fraction, we are likely over-estimating the hemicellulose fraction. This is likely due to errors associated with DNS based gross estimate of total reducing sugars using glucose as the standard (i.e., different class of reducing sugars like pentoses etc likely give a significant difference in the DNS responses).

#### (SI-V) Biomass Acid Chlorite Delignification Pretreatment Supporting Information

***Summary and Principle***: Lignin represents about 20-35% (total dry weight basis) of the mass in cellulosic biomass like corn stover and has been shown to deleteriously impact cellulose conversion by either physically impeding enzyme access to embedded cellulose or by reducing the effective concentration of available cellulases by non-specifically and irreversibly adsorbing enzymes to lignin.[7,8] Pretreatments that can effectively remove lignin can help avoid these issues. Depolymerization and extraction of lignin can be performed selectively using the acidified sodium chlorite method, originally known as the Wise method.[9] This process is usually performed at 60–70 °C for several hours with successive addition (every few hours) of fresh sodium chlorite and acetic acid at loadings of 0.3–0.6 g sodium chlorite/g biomass and 0.1–0.6 mL acetic acid/g biomass.[10] Here, the Wise delignification protocol can be used to showcase to students how lignin removal can dramatically impact overall substrate morphology and composition. Furthermore, the recalcitrant role of lignin towards cellulase-catalyzed hydrolysis of the pretreated substrate to fermentable sugars can be further highlighted. The following protocol can be used by the instructor specifically to prepare reagents for use by students as part of other learning activities OR can be used by students as part of a lab activity held in conjunction with other protocols outlined in this document.

***List of Chemicals and Materials for Experiment***: Sodium chlorite, glacial acetic acid, and sodium hydroxide were all procured from Fisher-Scientific, Sigma Aldrich, or VWR to be used as is. All lignocellulosic biomass substrates were a gift provided by the Great Lakes Bioenergy Research Center (GLBRC). Distilled water was used in the preparation of all aqueous solutions (unless specified otherwise). All necessary glassware, lab equipment, and supplies were used to carry out all experiments are highlighted here: Measuring cylinder (1-1000 mL range), Glass pipette (1-10 mL), Glass bottles with plastic screw caps (250 mL), and Whatman #41 filter paper. For all weighting operations an analytical balance with sensitivity of 0.0001 g was used. For heating solutions or drying solids a hot water bath and/or temperature-controlled oven can be used, respectively. Buchner filtration flask-vacuum line assembly is needed to filter separate solids from liquids.

***Hazards***: Glacial acetic acid and sodium chlorite are both hazardous in case of skin contact (irritant), eye contact (irritant), ingestion, or inhalation. Ideally all delignification protocols should be performed in the fume hood to avoid exposure to chemicals. All reagents must be handled with care and properly disposed. Researchers must wear laboratory gloves and eye protection glasses at all times. Boiling hot water (or hot ovens) can cause burns. Use heat resistant gloves when handling bottles immersed in a hot water bath or hot oven.

***Detailed Experimental Procedure***:
1. Weigh out 2 grams of dry milled corn-stover into a 250 mL wide-mouthed glass beaker
2. Add 175 mL of deionized RO water to the beaker (about 1.1% by weight of biomass) and uniformly mix/wet the biomass by swirling the bottle. See image below as example.

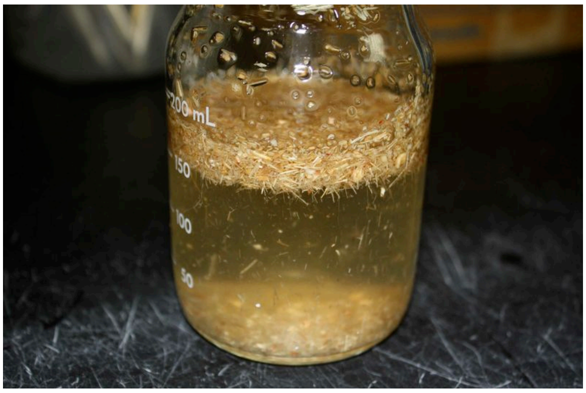

3. In a fume hood, add 8 g of sodium chlorite to the beaker and swirl gently (about 5% $NaClO_2$ by weight) to dissolve completely.
4. In a fume hood, add 3.5 mL of glacial acetic acid to the beaker and swirl gently to combine (about 2% Acetic Acid by weight). You will see bubbles rise in the bottle as shown in the image below. Do not close cap tightly and always keep bottle in fume hood!

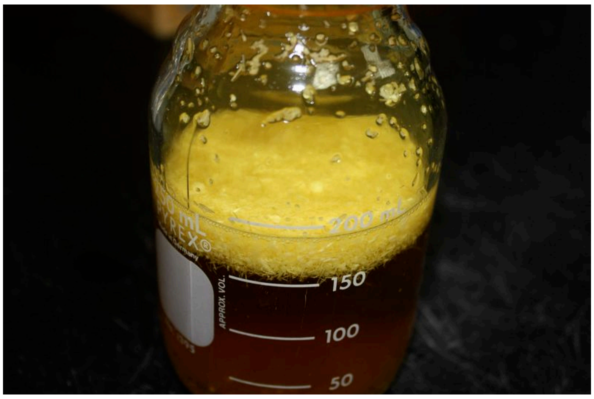

5. Screw on a cap with a small hole in the middle onto the glass bottle (or use parafilm with holes), submerge bottle to just below the cap into a water bath set at 70 degrees Celsius in a fume hood for two hours. Carefully swirl the bottle to mix contents every 15 minutes or so.
6. After the two-hour incubation period, add a second dose of both the 8 g of $NaClO_2$ and the 3.5 ml acetic acid into the bottle. Perform all additions always inside the fume hood. Make sure to add the $NaClO_2$ first and swirl *very* gently and then wait for the contents of the beaker to settle down a minute or so before adding the glacial Acetic Acid to avoid any spill over.
7. Return the loosely capped bottle to the 70 degrees Celsius water bath for two more hours, again swirling the bottle every 15 minutes.
8. After the second two-hour incubation (4 hours total incubation at this point of time), repeat step 6 by adding a third dose of both the 8 g of $NaClO_2$ and the 3.5 ml acetic acid.
9. Return the capped bottle to the 70 C water bath for two more hours, again swirling every 15 minutes.
10. At the end of the third two-hour incubation period (6 hrs total time), remove the beaker from the hot water bath and let it cool on the bench for about 20-30 minutes.
11. Set up a large Buchner vacuum funnel with Whatman #41 filter paper in a fume hood and pour the contents of the reaction beaker into the funnel with vacuum applied. Wash

the acid-chlorite treated biomass with copious amounts of water until biomass is white in color and the eluting water is close to neutral pH. Squeeze out remaining liquid and let the pretreated biomass dry in the fume hood overnight. See representative image below for 6-hour acid chlorite treated and original untreated corn stover samples generated using this protocol. Some biomass samples with partial delignification can be prepared by either 2 or 4-hour total reaction time as well, if needed.

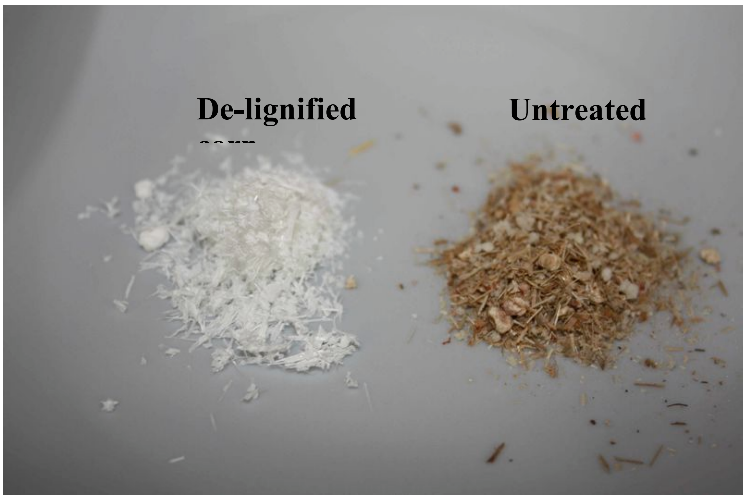

**Example Gravimetric Lignin Compositions Summary for untreated and acid chlorite treated corn stover for varying pretreatment time periods (on dry weight basis of starting material):**

| Untreated Corn Stover | 2 hr treated | 4 hr treated | 6 hr treated |
|---|---|---|---|
| 27% | 6% | 4% | 2% |

#### (SI-VI) Crystalline Cellulose Decrystallization Pretreatment Supporting Information

***Summary and Principle***: Cellulose is a linear polymer consisting of D-anhydroglucopyranose or glucose monosaccharide units joined together by β-1,4-glucosidic bonds to yield a polysaccharide composed of 100-1000's of glucose units. These polymer chains typically self-assemble during synthesis in plant cell walls to yield highly crystalline and ordered microfibrils that are composed of multiple cellulose chains joined together non-covalently via hydrogen bonding and van der Waals interactions. Crystalline cellulose has been shown to be highly recalcitrant to enzymatic hydrolysis. Pretreatments that can reduce cellulose crystallinity can enhance the rate of enzymatic hydrolysis of cellulose to glucose or cellobiose by several orders of magnitude. Here, we outline a simple pretreatment protocol develop by Walseth originally,[11] and later updated by Zhang and co-workers,[12] that uses cold 85% phosphoric acid to swell and decrystallize cellulose to reduce cellulose crystallinity. The swollen or dissolved cellulose is recovered from phosphoric acid by precipitation using cold water. This pretreated substrate is also called phosphoric acid swollen cellulose (PASC) and has been used in the literature to perform cellulase activity assays. Here, the modified Walseth decrystallization protocol can be used to showcase to students how cellulose decrystallization can dramatically impact overall substrate morphology. Furthermore, the recalcitrant role of cellulose crystallinity towards cellulase-catalyzed hydrolysis of the pretreated cellulose to fermentable sugars can be further highlighted. The following protocol can be used by the instructor specifically to prepare reagents for use by students as part of other learning activities OR can be used by students as part of a lab activity held in conjunction with other protocols outlined in this document. Lastly, instructors could consider using an alternative concentrated hydrochloric acid (HCl) based amorphous cellulose preparation protocol as well.[13]

***List of Chemicals and Materials for Experiment***: Avicel PH101 microcrystalline cellulose, phosphoric acid, sodium hydroxide, and sodium azide were all procured from Fisher-Scientific, Sigma Aldrich, or VWR to be used as is. Distilled water was used in the preparation of all aqueous solutions (unless specified otherwise). All necessary glassware, lab equipment, and supplies were used to carry out all experiments are highlighted here: Measuring cylinder (1-1000 mL range), Glass pipette (1-10 mL), Glass bottles with plastic screw caps (250 mL), Falcon conical plastic tubes (50 mL range), ice bucket, and Whatman #41 filter paper. For all weighting operations an analytical balance with sensitivity of 0.0001 g was used. For cooling solutions an ice bucket can be used. Buchner filtration flask-vacuum line assembly is needed to filter separate solids from liquids.

***Hazards***: Phosphoric acid is hazardous in case of skin contact (irritant), eye contact (irritant), ingestion, or inhalation. All reagents must be handled with care and properly disposed. Researchers must wear laboratory gloves and eye protection glasses at all times.

***Detailed Experimental Procedure***:

1. Weigh out 200 mg of microcrystalline cellulose (or Avicel PH-101) into a 50 mL Falcon or 250 mL glass bottle.
2. While swirling the tube, slowly add 10 mL of ice-cold phosphoric acid (~85% by weight). Ensure complete saturation of the Avicel by vigorous mixing; breakup any clumps with a glass stirring rod.

3. Let the sample tube sit on ice for at least one hour to fully swell and pretreat cellulose to produce soluble cellulose gel. Note that cold $H_3PO_4$ should cause the Avicel to swell into a gel-like state that is translucent and does not contain any traces of unswollen solids. See image below for illustration.

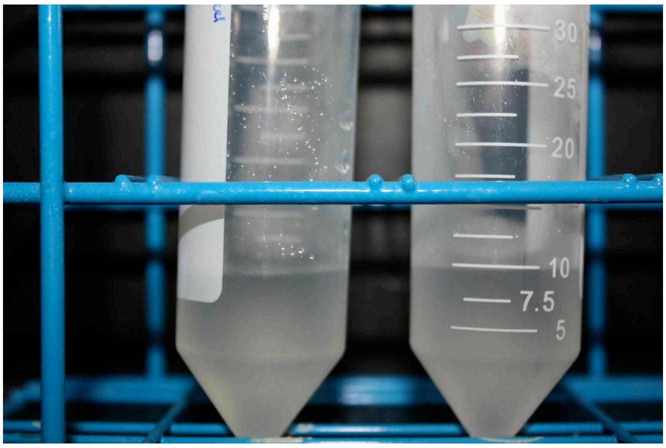

4. While keeping the tube on ice, add 40 mL of ice-cold water to the 50 mL Falcon tube in 10 mL increments, mixing thoroughly in between each addition, in order to precipitate dissolved cellulose out of concentrated acid solution in to its highly amorphous and decrystallization form. See image below as illustration.

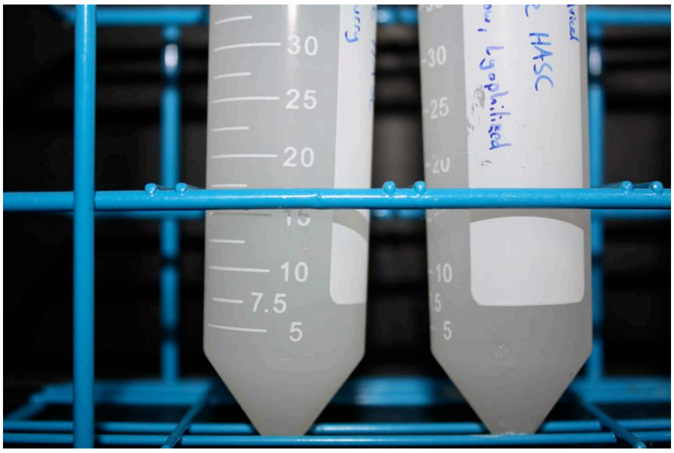

5. Centrifuge the tube containing the precipitated swollen cellulose for 5 minutes at 4200 rpm to pellet the cellulose. Discard the supernatant. Alternatively, in the absence of a centrifuge, the samples can be washed using a Buchner filter assembly.
6. Re-suspend the amorphous cellulose pellet with 40 ml of ice-cold water to wash away remaining acid. Centrifuge again at 4200 rpm for 5 minutes and discard the supernatant. Alternatively, in the absence of a centrifuge, the samples can be washed using a Buchner filter assembly. Repeat this washing process until the supernatant gives a neutral pH when tested with pH paper.
7. The neutralized, washed, phosphoric acid swollen microcrystalline cellulose or PASC can then be stored in a slurry form at 4C for one week as is or up to two months if sodium azide (0.05% w/v) is added. The cellulose pellet can also be frozen in liquid nitrogen and then lyophilized to store in dry form for future work.

#### (SI-VII) Cellulosic Biomass Enzymatic Hydrolysis Supporting Information

***Summary and Principle***: Insoluble polymeric carbohydrates (like β-1,4-glucan or cellulose and β-1,4-xylan which is a predominant hemicellulose in grasses) are locked inside plant materials as high molecular weight polysaccharides and oligosaccharides. Cellulose and hemicellulose can be hydrolyzed into their respective acid soluble monomeric forms (i.e., glucose and xylose) using suitable enzymes. Enzymatic hydrolysis is a key step for converting cellulosic biomass into sugars (using synergistic cellulase and hemicellulase enzymes as biocatalysts) in a bio-refinery to produce biofuels. However, enzymes are currently still expensive to produce (~$10/kg). There is a critical need to lower costs by minimizing the enzyme amount needed to achieve 100% reaction completion (by hydrolysis of cellulose and/or hemicellulose to soluble fermentable sugars like glucose or xylose) in a short period of time. The cellulases and hemicellulases, based on compositional analysis of most commercially available cellulase cocktails,[14] include diverse families of carbohydrate-active enzymes or CAZymes.[15] The critical families of CAZymes identified necessary to hydrolyze most plant cell wall polysaccharides in untreated or pretreated biomass include endo-glucanases, endo-xylanases, cellobiohydrolases, β-glucosidases, and β-xylosidases. Endo-glucanases randomly hydrolyze internal glycosidic bonds within cellulosic polymer chains, while cellobiohydrolases act processively along cellulose polymer chains cleaving off cellobiose or soluble gluco-oligomers, while with β-glucosidases ultimately hydrolyze gluco-oligomers to glucose. Similarly, endo-xylanases cleave the xylan backbone at internal β-1,4 xylosidic bonds, while β-xylosidases hydrolyze short xylo-oligomers to xylose. There are several other classes of accessory CAZymes that synergistically work with these core cellulase and hemicellulases.[2,16] Regardless, all these enzymes are thought to work synergistically, creating new accessible substrates or sites for each other to act upon.

A simple protocol titled NREL/TP-5100-63351 was originally developed by the National Renewable Energy Laboratory (NREL) to test the hydrolytic activity of commercially available cellulase/hemicellulase cocktails or individual enzymes and is available online at (https://www.nrel.gov/bioenergy/biomass-compositional-analysis.html). Our current method, inspired by NREL's standardized method, utilizes a readily available lab supplies like a hot water-bath (and/or regular oven) and common chemistry lab apparatus to setup reactions to hydrolyze the insoluble carbohydrate polymers into soluble sugar monomers using commercially available cellulase enzyme cocktails. The soluble sugars in the biomass hydrolyzate solutions can then be assayed using the customized sugar assays described earlier to provide a comprehensive composition analysis for a diverse range of cellulosic feedstocks. The following protocol can be used by the instructor specifically to prepare reagents for use by students as part of other learning activities OR can be used by students as part of a lab activity held in conjunction with other protocols outlined in this document.

***List of Chemicals and Materials for Experiment***: Avicel PH101 microcrystalline cellulose, glucose, xylose, mannose, galactose, arabinose, cellobiose, sodium citrate, citric acid, Celluclast cellulase cocktail, Novo 188 cellobiase cocktail, sodium azide, and sodium hydroxide were all procured from Fisher-Scientific, Sigma Aldrich, or VWR to be used as is. C_TEC2 cellulase cocktail was a gift from Novozymes. All lignocellulosic biomass substrates were a gift provided by the Great Lakes Bioenergy Research Center (GLBRC). Distilled water was used in the preparation of all aqueous solutions (unless specified otherwise). All necessary glassware, lab equipment, and supplies were used to carry out all experiments are highlighted here: Single/multi-channel micropipettes and associated plastic tips (10, 100, 1000 μL range), Glass pipette (1-10 mL), Glass tubes with plastic screw caps (10 mL), Falcon conical plastic tubes (15, 50 mL range), Volumetric flask (100, 1000 mL), 96-well clear flat-bottom microplates (0.3 mL volume per well), PCR tubes with caps (0.2 mL), aluminum foil, and spectrophotometer/microplate reader. For all weighting operations an analytical balance with sensitivity of 0.0001 g was used. For heating solutions or drying solids a hot water bath and temperature-controlled oven can be used, respectively.

***Hazards***: Sodium azide is very hazardous in case of skin contact (irritant), of eye contact (irritant) and can cause death in cases of over exposure (avoid use if possible). Sugar containing buffers/solutions may cause eye and skin irritation. Enzyme/sugar containing buffers/solutions may cause eye and skin irritation. Enzyme kit reagents solutions are not to be consumed and are also not supposed to be used by students for testing sugar concentrations in any other bodily fluids. All reagents must be handled with care and properly disposed. Researchers must wear laboratory gloves and eye protection glasses at all times. Boiling hot water (or hot ovens) can cause burns. Use heat resistant gloves when handling tubes immersed in a hot water bath or hot oven.

***Stock Solution Preparation Experimental Procedure***:

1. Wear suitable personal protection equipment (PPE) before starting experiment: Gloves, Eye-Goggles, and Lab coat.
2. Prepare stock sugar standards for regular assays, as explained below.
    a. Glucose Stock Solution: Add 1.000 g glucose to 50 mL of water in 100 mL volumetric flask. Plug flask and invert to dissolve. Make up volume to 100 mL with addition of deionized water and mix well to fully dissolve glucose to prepare 10 g/L stock solution.
    b. Next, prepare water blank (0 g/L control) and glucose standards (1 g/L, 2 g/L, 3 g/L, and 5 g/L) by preparing suitable dilutions (in deionized water) of original 10 g/L stock in 50 ml plastic conical Falcon tubes. Additional glucose standards of varying concentrations can be prepared as well, if necessary, by serial dilutions.
    c. Label all glucose standards tubes and store at $4^{o}$ C, if fridge is available.
3. Prepare other sugar (xylose, mannose, galactose, arabinose and cellobiose) standards using similar protocol outlined above.
4. Prepare enzyme stock solutions based on enzyme loading desired during cellulose hydrolysis reaction. Protein concentration will be provided by the commercial vendor or can be estimated using standard analytical techniques highlighted elsewhere.[14] Here, stock protein concentrations of Celluclast cellulase cocktail, Novo 188 cellobiase cocktail, and C_TEC2 cocktail were estimated using the Bradford assay method using bovine serum albumin (BSA) as standard to be 56 mg/mL, 26 mg/mL, and 86 mg/mL, respectively. For assays with cellulose, we used Celluclast or C_TEC2 from Novozyme (both are available from Sigma-Aldrich). For assays with cellobiose, we used Novo 188 based cellobiase cocktail (Aspergillus) from Novozyme which was recently discontinued by Sigma-Aldrich (but might be requested directly from Novozymes). Alternatively, one can use other cellobiase enzymes commercially available from Sigma-Aldrich (Catalog No. 49291 or G4511). Handle enzyme stock solution with care using appropriate PPE and store on ice in suitable capped plastic tubes or glass bottles. Do not use diluted enzymes stored in fridge for longer than a couple of days to minimize loss in enzyme activity.

Commercially available cellulase stocks are supplied as solutions containing stabilizers that prevent significant loss in enzyme activity when stored at 4 degrees Celsius for several months to years. However, dilution of stock enzyme into buffer of choice is not desirable beyond a couple of days in the fridge for classroom or demo activities.

5. Prepare 1 M sodium citrate buffer stock (pH 4.3) solution and store in capped glass bottles for long-term storage at 4 degrees Celsius. Prepare 20 g/L sodium azide stock solution as antimicrobial additive for overnight hydrolysis reactions, if needed.
6. Refer to appropriate protocols (see SI-I to SI-III) for detection of released soluble sugars.
7. Determine total moisture content of each biomass sample by drying in oven set at 100°C, ideally overnight, to remove any residual moisture till constant dry weight.

***Detailed Experimental Procedure***:

1. Weigh out the appropriate substrate amount in to a 15 mL Falcon tube or glass tube. Example: For lignocellulosic biomass of known composition, add 20 mg of cellulose* equivalent dry weight of material for a 10 mL reaction volume. For Avicel PH-101 or amorphous cellulose assays, add either desired cellulose amount (e.g., 25, 50, 200, and/or 250 mg depending on enzyme loading) to get a significant glucose release within 90 min hydrolysis time based on the actual student activity protocol.
    a. Note that it is possible to scale up the volume or amount of substrate concentration in a 15/50 mL Falcon tube for a 10 ml (or greater) reaction volume if needed. Scale the enzyme and buffer concentrations accordingly.
    b. 20 mg cellulose equivalent could mean 20 mg of weighed out pure cellulose like Avicel, or 20 mg of cellulose in slurry form that is pipetted into the reaction tube, or ex. ~50 mg lignocellulosic biomass that contains ~40% cellulose based on compositional analysis.
    c. Note that 20 mg cellulose in 10 mL total reaction volume is equivalent to 0.2% substrate concentration that would give a theoretical maximum glucose concentration of 2.22 g/L. While, a higher cellulose concentration (250 mg) would a theoretical maximum glucose concentration of 27.75 g/L. Here the conversion factor for cellulose (anhydroglucose) to glucose is 180/162 = 1.11.
2. Determine how much enzyme is to be used in the reaction. If the intended enzyme concentration (per mass unit of added cellulose) is to be 15 mg enzyme/g cellulose, and you have 20 mg of cellulose in the reaction volume, you will need 300 μg of enzyme per tube. Use Bradford assay or known specifications from enzyme manufacturer to determine stock enzyme protein concentration. Here, stock protein concentrations of Celluclast cellulase cocktail, Novo 188 cellobiase cocktail, and C_TEC2 cocktail are provided by the instructor to be 56 mg/mL, 27 mg/mL, and 86 mg/mL, respectively. Stock enzymes may need to be diluted before use in order to reduce pipetting errors. Wait until all other reagents/buffers are added to the reaction tubes before finally adding in the enzyme solution.
3. Determine the volume of stock buffer that is to be added to each reaction tube. Add enough concentrated buffer to the reaction tubes so that once diluted to the final reaction volume, the buffer concentration is 50 mM at the appropriate pH for the specific enzymes being used. For example, in a 10 mL reaction volume, add 500 μL of a 1 M Na-Citrate pH 4.3 buffer so that the final concentration and pH are 50 mM and pH ~4.5 respectively. See representative images below.
4. Determine the volume of antimicrobial agent (if the samples will be incubated overnight) that will be added to each reaction tube. Add an anti-microbial agent, such as

sodium azide, at a final concentration of 0.02-0.05 % (w/v). For example, in a 10 mL reaction volume, 5 mg of $NaN_3$ is needed so one could add 250 μL of a 20 mg/ml stock sodium azide solution. For hydrolysis reactions carried out over only 0.5-4 hours, there is no need to add any antimicrobial agent.

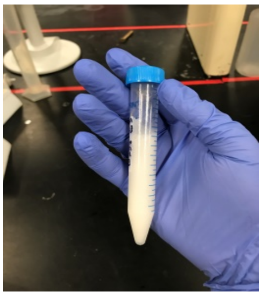

*Sample tube ready for incubation*

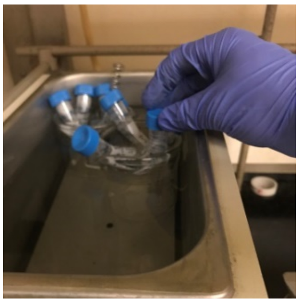

*Sample tubes incubated in water bath*

5. Determine the final make up volume of water that will be added to each reaction tube. Calculate this by taking the predetermined final reaction volume and subtracting out the calculated volumes of substrate slurry, enzyme, buffer, and sodium azide solutions. The remaining volume is the amount of deionized/sterile water that is to be added to each reaction.
6. Once all volumes are determined, add to the reaction tubes all of the reagents in the following order; substrate, water, buffer, sodium azide, and lastly followed by suitable diluted enzyme solution.
7. Seal reaction tube lids with Parafilm and incubate in oven at desired temperature (50 Degrees Celsius) to begin the reaction, either in a water bath, oven, or on a heating block. Reaction tubes can be left alone to react in stationary conditions without mixing, or they can be rotated in a rotational oven, or they can be left in a shaking hot plate block at 500-1000 rpm. Reaction times can range from 0.5 hours to several days depending on the reaction conditions, type of enzymes and type of substrate.
8. Upon completion of reaction time, centrifuge the reaction tubes to pellet solids, or let the solid settle slowly before removing supernatant for sugar analysis. If the substrate is insoluble, remove some of the supernatant and heat in sealed tubes to 95 C for 5 minutes to denature the enzymes and ensure halting the reaction. For soluble substrates, it may be necessary to increase pH > 10 (if DNS assay will be done eventually) to denature enzymes and/or halt reaction. Once the reaction is halted, proceed with sugar analysis (e.g., DNS, TRUEbalance/TRUEtest, HK-G6PH Enzyme assays etc). For insoluble substrates, decant small volume of the supernatant and proceed directly to sugar analysis with it (as described in earlier supplementary protocols). See image below as illustration.

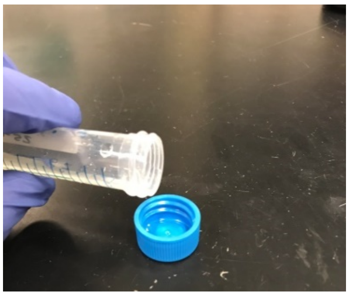

**1. Representative results expected from outreach activity conducted by students exploring the effect of cellulase enzyme (E) concentration, cellulosic substrate (S) concentration, and hydrolysis reaction temperature on the relative rate of native microcrystalline cellulosic substrate conversion into glucose mimicking a cellulosic biorefinery enzymatic hydrolysis unit operation**:

Here, a total reaction volume of 10 mL in a stoppered plastic test tube, containing either 50 (Low S) or 200 (High S) mg of untreated microcrystalline cellulose (i.e., Avicel PH-101) was hydrolyzed using a commercially available cellulase enzyme cocktail at either a 5 mg enzyme per gram cellulose (Low E) or 50 mg enzyme per gram cellulose (High E) enzyme loading at two different reaction temperatures (i.e., 25 and 50 $^{o}$C). The enzymatically released soluble sugars (i.e., glucose) was measured using the TRUEBalance test kit after 90 mins of total reaction time (under static conditions with intermittent end-over-end mixing every 30 mins) with the actual glucometer readings reported here. Commercially available C_TEC2 cellulolytic enzyme cocktail (gift from Novozymes) was used for assays here. Data shown in the main text (Fig. 3) are examples of assays run by one Rutgers undergraduate student to validate the protocol (with no replicates). Therefore, authors provide additional supporting data from eight replicate assays conducted per condition to highlight the reproducibility of overall activity protocol and glucometer assay.

| Hydrolysis Reaction Temperature | Cellulose Conc. Added (Avicel PH-101) | Enzyme Conc. Added (C.Tec2) | TRUEBalance Meter Readings for Replicate Hydrolysis Samples | | | | | | | | Average Reading | Standard Deviation |
|---|---|---|---|---|---|---|---|---|---|---|---|---|
| | | | Replicate 1 | Replicate 2 | Replicate 3 | Replicate 4 | Replicate 5 | Replicate 6 | Replicate 7 | Replicate 8 | | |
| 25 Degrees ℃ | 50 mg | 5 mg/g | 0 | 0 | 0 | 0 | 0 | 0 | 0 | 0 | 0 | 0 |
| | | 50 mg/g | 136 | 136 | 119 | 123 | 130 | 119 | 136 | 124 | 128 | 8 |
| | 200 mg | 5 mg/g | 102 | 106 | 104 | 104 | 106 | 109 | 110 | 104 | 106 | 3 |
| | | 50 mg/g | 432 | 442 | 431 | 444 | 442 | 439 | 440 | 437 | 438 | 5 |
| 50 Degrees ℃ | 50 mg | 5 mg/g | 40 | 36 | 38 | 36 | 35 | 37 | 43 | 42 | 38 | 3 |
| | | 50 mg/g | 228 | 231 | 217 | 215 | 227 | 225 | 230 | 227 | 225 | 6 |
| | 200 mg | 5 mg/g | 308 | 304 | 309 | 311 | 308 | 305 | 302 | 306 | 307 | 3 |
| | | 50 mg/g | 586 | 583 | 549 | 539 | 566 | 570 | 564 | 575 | 567 | 16 |

**2. Examples of amorphous cellulose versus cellobiose enzymatic hydrolysis kinetic profiles for at fixed enzyme loadings using DNS or TRUEbalance glucose assays for advanced enzymatic hydrolysis kinetic assays shown here as reference data for the instructor:**

Example of amorphous cellulose enzymatic hydrolysis kinetic profiles for varying time points at fixed cellulase cocktail loadings using DNS glucose assays are shown below. Briefly, a total reaction volume of 5 ml in a stoppered plastic test tube, containing either 20 mg of lyophilized amorphous cellulose was hydrolyzed using a commercially available cellulase enzyme cocktail at a low enzyme loading (1.5 mg Celluclast and Novo 188 enzymes each per gram cellulose). The samples were hydrolyzed at 50 $^{o}$C without shaking in a pH 4.5 Na-Citrate (50 mM) buffer also containing sodium azide for a total reaction period of 24 hours. Avicel cellulose was pretreated and decrystallized for 2 hrs using the phosphoric acid pretreatment process reported in elsewhere in supplementary information. The enzymatically released soluble sugars were measured using the DNS assay method using glucose as standard (similar results are expected with TRUEbalance assay). Please note that the reaction volume can be scaled up to 10 ml if desired to follow the identical SI-VII protocol.

Example of cellobiose enzymatic hydrolysis kinetic profiles for varying time points at fixed cellulase cocktail loadings using TRUEbalance glucose assays are shown below. Briefly, a total reaction volume of 0.25 mL in a stoppered plastic micro-centrifuge tube, containing either 2.5 mg of cellobiose was hydrolyzed using a commercially available cellulase enzyme cocktail at a low enzyme loading (1 mg Novo 188 enzyme per gram cellobiose). The samples were hydrolyzed at 50 °C without shaking in a pH 4.5 Na-Citrate (50 mM) buffer also containing sodium azide for a total reaction period of 24 hours. All assays were conducted in duplicate with error bars depicting a single standard deviation variation from reported mean results. Samples were taken at multiple time points by arresting the reaction after addition of 0.25 mL HEPES pH 7.5 buffer followed by immediate analysis using the TRUEbalance test strip assay to estimate total glucose released. The enzymatically released soluble sugars were measured using the TRUEbalance assay method using glucose as standard (Note that DNS assay cannot be used here due to high background signal by cellobiose). Please note that the reaction volume can be scaled up to 10 mL if desired to follow the identical SI-VII protocol.

Representative experimental results expected for both these specific enzyme kinetic assays for amorphous cellulose (left-A) and cellobiose (right-B) are reported below. This data can be used to generate initial reaction velocity profiles (as µmoles glucose released per mg added crude enzyme cocktail) for bonus question 6 reported in advanced student activity protocol data analysis questionnaire.

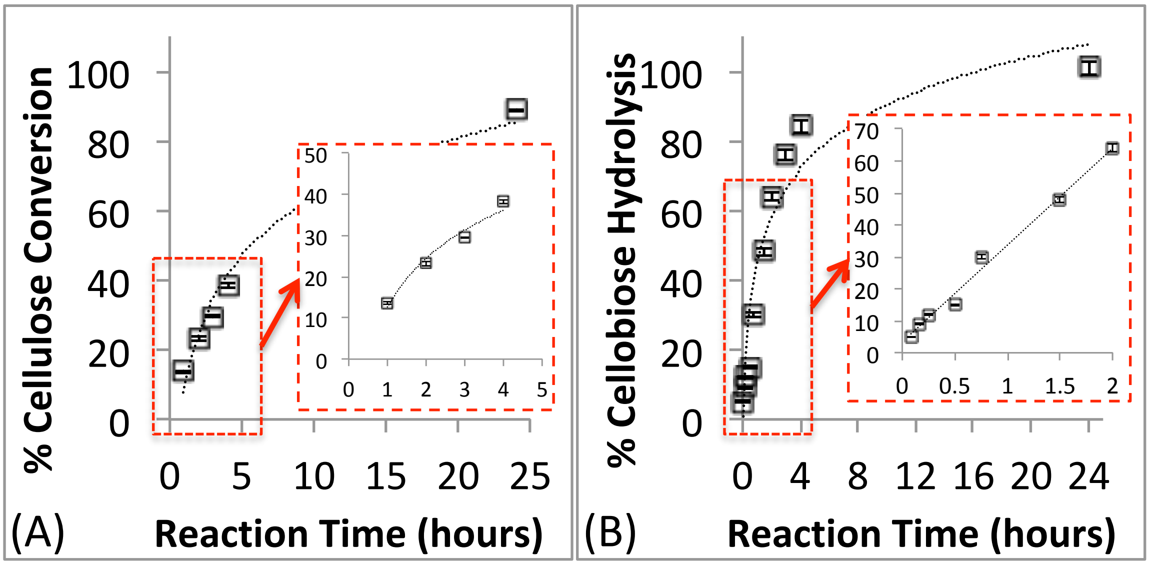

### Grass to Gas! – Pre-Activity Questionnaire/Survey (5 mins)

**Student Name/Group Number**:

**Part 1 – Before starting the experiment or reading additional handout materials on activity background, please answer these questions briefly**

**Question 1**: What are carbohydrates, monosaccharides, & polysaccharides? List as many examples of well-known carbohydrates that you find at home or elsewhere.

**Question 2**: What are enzymes & how do they work? List any example that you can think of.

**Question 3**: What is the impact of temperature on the rate of a chemical reaction? Can you briefly outline any theory that explains how temperature impacts chemical reaction rate?

**Question 4**: List as many examples of biofuels or lignocellulose biorefining processes used to produce biofuels or biochemicals in industries that you are familiar.

**Question 5**: Do you know what chemical and biochemical engineers do at work? List examples of job profiles of chemical and biochemical engineers.

### Grass to Gas! – Student Activity Background (10 mins)

**Student Name/Group Number**:

**Part 2 – Work with your partner to analyze your data and answer these questions**

**Background**: There are a variety of plants (e.g., corn, switchgrass) that grow around the world in a renewable and sustainable manner. Plant derived biomass are mostly composed of carbohydrate-based polymers such as sucrose, starch, and/or cellulose (e.g., ~70% of dry wood on total weight basis is composed of cellulose and hemicellulose alone). Therefore, cellulose is an abundant feedstock that humans can convert into useful bioproducts as fossil resources like petroleum dwindle in the coming centuries. For example, glucose derived from cellulosic biomass by processing it in bio-refineries (e.g., corn ethanol refinery) provides access to a renewable carbon-based feedstock that can then be easily fermented in biofuels like ethanol. However, researchers are working diligently to efficiently 'refine' biomass by overcoming various practical challenges associated with its bioconversion.

Firstly, solubility and chemical reactivity of carbohydrate-based polymers is closely dependent on its complex molecular structure. Plant-based carbohydrates are often composed of individual sugar monomer units joined together to form larger branched or linear shaped polymeric structures. These polymers are further assembled together to form more complex structures in plant biomass that reduces their accessibility to water. Some carbohydrate polymers like sucrose are highly soluble in water, while some larger sized polymers like cellulose are poorly or not soluble in water. Smaller fragment of cellulose polymers, like cellobiose (a model cellulose derived disaccharide as shown below), are more soluble in water.

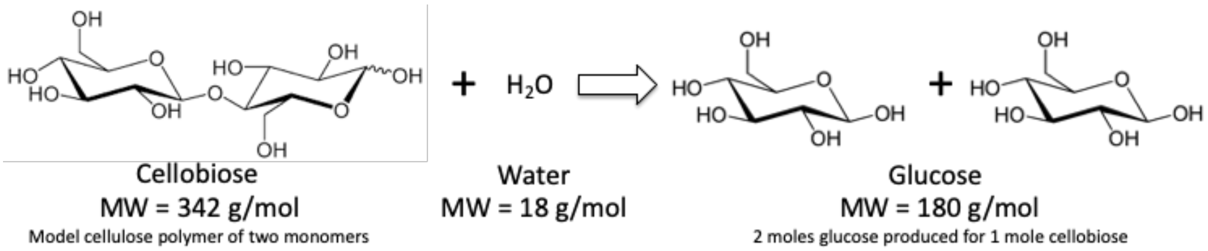

Secondly, the spontaneous hydrolysis rate of β-glycosidic bonds of cellulose in water is 'very' slow and it would take ~$10^{15}$ seconds (or 0.1 billion years) in the absence of any catalysts like sulfuric acid or hydrolytic enzymes to speed up this reaction.[17,18] Carbohydrate-derived polymers or polysaccharides like cellulose can be hydrolyzed into their constituent monomer units (e.g., simple soluble sugars like glucose as shown in image below) by water. From a reaction stoichiometry point of view, one cellobiose (Molecular Weight or MW = 342 g/mol) molecule hydrolysis reaction with water (MW = 18 g/mol) results in the formation of two molecules of glucose (MW = 180 g/mol).

Enzymes like cellulases (or cellobiases) are very efficient biocatalysts that can speed up the hydrolysis rate of cellulose (or cellobiose) into glucose to achieve complete conversion in a few hours to days. However, there are many challenges associated with enzymatic hydrolysis of biomass derived carbohydrate polymers. In this activity, we will analyze how some of the factors listed here impact the rate of enzymatic hydrolysis of cellulose (or cellobiose) to glucose; Biomass Structure, Biomass Concentration, Enzyme Concentration, and Hydrolysis Reaction Temperature.

### Grass to Gas! – Post-Activity Survey & Data Analysis (30 mins)

**<u>Student Name/Group Number</u>**:

**<u>Part 2 – Work with your partner to analyze your data and answer all these questions</u>**

**Question 1**: Clearly list the reaction conditions (e.g., enzyme loading, i.e. $\frac{mg\ Enzyme}{g\ Cellulose}$, cellulose or cellobiose substrate concentration, and reaction temperature) that you were assigned for the enzymatic hydrolysis experiment.

**Question 2**: List if your sample in the tube dissolved immediately upon addition of the enzyme/buffer solution or was it insoluble? Did you notice any change in the insoluble sample size, morphology, and/or total reaction volume before and after enzymatic hydrolysis for 90 mins? Try to explain your findings.

**Question 3**: Would increasing reaction temperature cause enzymes to work faster or slower? Is there an upper limit for the maximum temperature possible to be used? Why?

**Question 4**: In a biorefinery, $Cellulose \xrightarrow{Enzyme} Glucose \xrightarrow{Yeast} Ethanol$. Will increasing cellulose concentration during hydrolysis make it cheaper to produce bioethanol? Why? *Hint: Think energy-intensive operation needed in biorefinery to produce anhydrous ethanol.*

**Question 5**:

(A) Based on the calibration curve data provided below for TrueBalance test strips on actual glucose standards of known concentrations and the results displayed on your glucometer display for your unknown concentration samples, can you estimate the true glucose concentration in your sample using either the fitted equation or graphically?

(B) Knowing the actual glucose concentration in your recovered enzymatic hydrolysate sample, can you estimate what percent of the initial cellulose (or cellobiose) substrate was converted into glucose? Show your work below. *Hint: Use the reaction stoichiometry to estimate how many moles (or equivalent mass) of glucose would be produced for every mole (or equivalent mass) of cellulose or cellobiose, depending on actual substrate.*

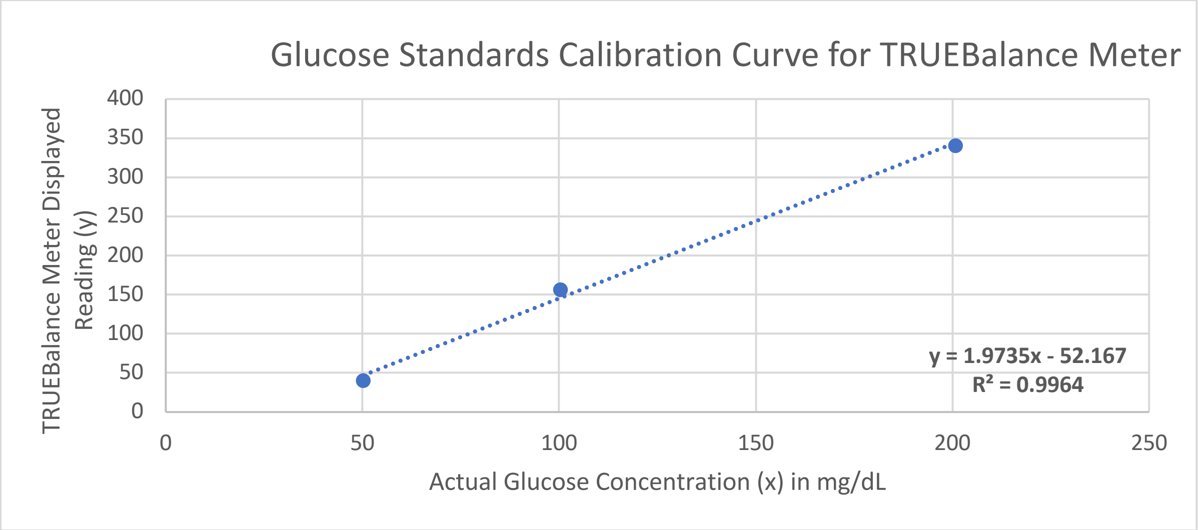

(SI-IX) 'Grass to Gas!' Outreach Activity Student Protocol

### Grass to Gas! – Outreach Activity Student Protocol (~90 mins)

**Note that this 'Grass to Gas! – Enzymatic Hydrolysis Protocol' is an abridged version of the reported detailed assays described in SI-VII. The instructor can pre-weigh the cellulose (or cellobiose) into tubes and prepare the diluted enzyme-buffer stock solutions ahead of time to save time (see SI-X for detailed instruction for instructors on pre-activity prep-work). Attached Excel file also provides a visual guide for the prep-work conducted prior to starting the activity. Students can start working on the post-activity survey/questionnaire immediately after starting the hydrolysis reaction in step 5.**

1. Safety first! Put on your gloves and safety goggles.

2. Label your falcon tube using a marker with your name.

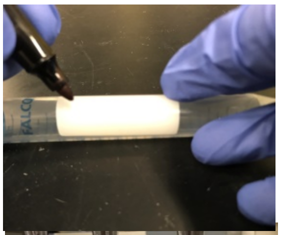

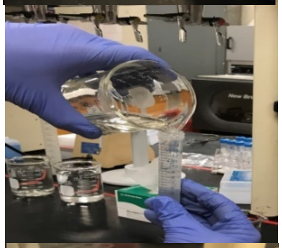

3. Decant[1] 10 ml of the dilute enzyme[2] solution from the beaker of matching color into the falcon tube containing your cellulose[3] substrate.[4]

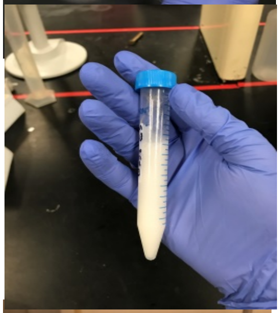

4. Tightly screw on the cap and shake vigorously for 30 seconds until the substrate is thoroughly suspended and mixed (or dissolved).

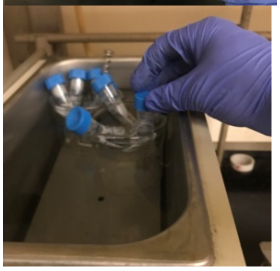

5. **If your falcon tube has a black strip of tape** -place your sample in the incubator at 50°C for 90 minutes.

   **If your falcon tube has a white strip of tape** -place your sample in the tube rack at room temperature (25°C) for 90 minutes.

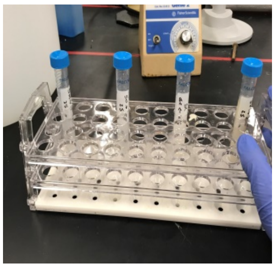

6. Withdraw your sample from its beaker; do not shake the sample to disturb the settled solids (if present)!

---

[1] **Decant** – Gradually pour solution from one tube/container to another
[2] **Enzyme** – Biological catalyst that accelerates chemical reactions
[3] **Cellulose** – A polysaccharide consisting of multiple units of glucose linked to form polymer
[4] **Substrate** – The substance on which the enzyme acts upon

7. Uncap the sample tube, and pour a few milliliters of the solution supernatant[5] into the tube cap.

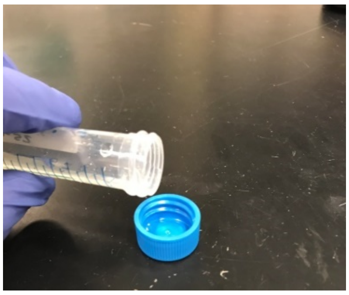

8. Obtain a test strip from the glucometer bottle, immediately recap bottle after.

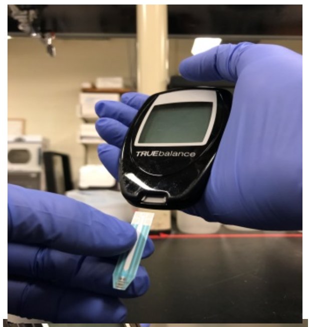

9. Insert the test strip into glucose meter (as shown in image here), and the meter will turn on automatically.

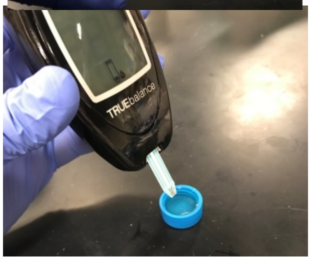

10. When the meter is ready, dip the end of the test strip end into your solution meniscus (do not fully immerse the strip into the solution). Immediately remove the meter/tip from the cap after the meter beeps.

11. Wait for ten seconds, and then read display to find the glucometer displayed reading given in mg/dL[6] units. Repeat in case of meter error reading using a fresh test strip.

12. Please note that this glucometer reading (say 'y') will need to be used along with the meter calibration equation (y = 1.9735x - 52.167) to estimate the actual glucose concentration in your sample solution ('x' calculated from the calibration equation with concentration units of mg/dL).

[5] **Supernatant** – The liquid lying above a solid residue
[6] **mg/dL** – Milligrams per deciliter (dissolved glucose concentration in solution). Note that 100 mg/dL is equivalent to 1 mg/mL (milligrams per milliliter).

#### (SI-X) 'Grass to Gas!' Outreach Activity Instructor Protocol & Notes

### Grass to Gas! – Instructor Protocol & Notes

**Note that this 'Grass to Gas! – Instructor Pre-Activity Protocol & Notes' is an abridged version of the reported SI-VII assay. Here details are provided on how exactly the instructor or staff should pre-weigh cellulose (or cellobiose) into tubes and prepare the diluted enzyme-buffer stock solutions immediately before the student activity.**

**Pre-activity Setup Protocol:** The aim of this pre-activity protocol is to prepare a number of substrate samples and several different dilute enzyme solutions at various concentration ranges for subsequent usage in student enzymatic hydrolysis activity protocol reported in SI-IX. C-Tec 2 stock cellulase enzyme solution must be diluted with a sodium citrate buffer (pH 4.5) and DI water before being used by students during the actual activity for addition into the cellulosic substrate containing tube to begin enzymatic hydrolysis. Preparing stock dilute enzyme solutions and stock substrate samples ahead of time allows for a more effective hands on demonstration for participating students within a 2-3 h total outreach activity time. The instructor should refer to SI-VII protocol and relevant notes to identify the ideal enzyme and substrate loading range if using cellobiose for the student activity instead. Here we only provide details on assays conducted using the cellulose-cellulase system. See accompanying Excel file with additional details on the color-coding scheme used to prepare tubes and samples for the student outreach activity.

1. **Hazards & Safety:** Ensure all relevant safety guidelines are followed while working with laboratory reagents and equipment. Refer to SI-VII protocol for details.

2. **Experimental Supplies Needed for Student Activity:**

   **Reagents (Chemicals, Catalog Number, and Vendor Details):**

   *a.* Cellic C-Tec 2 cellulase enzyme from Novozymes (Gift from Novozymes with stock concentration of 40 mg/mL used for all subsequent dilutions reported here; also available commercially from Sigma-Aldrich; Catalog No. SAE0020-50ML; $105). *Note: Alternatively, Celluclast 1.5L cellulase enzyme manufactured by Novozyme (from Trichoderma) is also commercially available from Sigma-Aldrich (Catalog No. C2730-50ML; $99).*
   b. Cellulose (Avicel PH-101, Sigma-Aldrich; Catalog No. 11365-1KG; $125)
   c. Sodium citrate (Trisodium citrate-dihydrate, Sigma-Aldrich; Catalog No. S4641-500G; $40).
   d. Trividia Health TRUEBalance Meter Test Strips (50 Ct-Box, Trividia/Amazon, ~$20-30)
   e. Deionized (DI) Water

   **General Lab Glassware and Consumables:**

   a. Graduated Cylinder (100 mL)
   b. Plastic Conical Bottom Falcon Tubes (15 mL)

c. Graduated Beakers (250 mL)
d. Pipette tips (20 µL, 200 µL, 1000 µL)

**Equipment:**

a. Analytical Balance (with minimum 0.1 mg accuracy)
b. Single-channel Micropipettors (20 µL, 200 µL, 1000 µL)
c. Trividia Health TRUEBalance Meter (1 Starter Kit Meter, Trividia/Amazon, ~$20-30)

#### 3. Pre-Activity Checklist:

Read the entirety of this protocol before beginning the experiment.

Wear safety gear (Lab Coat, Gloves, and Safety Goggles) before you begin with the experiment

Determine the total number of students participating in the outreach activity so the appropriate amount of substrate samples and dilute enzyme solutions can be prepared.

Label all 15 mL falcon tubes according to the substrate sample added. Label each dilute enzyme solution according to its concentration.

#### 4. Pre-Activity Experimental Procedure:

1. Measure out the appropriate number of cellulose substrate samples into aliquoted amounts of 50 mg and 200 mg into individual 15 mL falcon tubes. *Note: For amorphous cellulose that is present in wet slurry like state, transfer slurry directly using pipette based on actual slurry solids concentration (e.g., To add 50 mg cellulose on a dry weight basis, transfer 5 mL of 10 mg/mL PASC slurry).*
2. Label half the falcon tubes with 50 mg substrate with a blue strip of tape, and the other half with a red strip of tape.
3. Label half the falcon tubes with 200 mg substrate with a gray strip of tape, and the other half with a yellow strip of tape.
   *(Depends on total number of students in class)*

4. Prepare the enzyme solution $\left(0.025\ \frac{mg\ enzyme}{mL\ solution}\right)$, for delivering 5 mg/g-glucan enzyme loading to 50 mg substrate samples, in a 250 mL beaker (labeled pink).
   - Add 2000 µL of stock 1M Na-Citrate buffer into 97 mL DI water. *Note: To prepare 0.1 L Na-Citrate stock buffer (1M) dissolve 29.4 g trisodium citrate-dihydrate in DI water and adjust to pH 4.5 using 1N HCl. Stock buffer can be stored at room temperature for 3 months or at 4°C for upto 6 months.*
   - Add 62.5 µL C-Tech 2 into the buffer solution. *(Prepares enough for 10 samples)*
   - Make up to 100 mL volume by slowly adding DI water.

5. Prepare the enzyme solution 1 $\left(0.10\ \frac{mg\ enzyme}{mL\ solution}\right)$, for adding 5 mg/g-glucan enzyme loading to 200 mg substrate samples, in a 250 mL beaker (labeled red).
   - Add 2000 µL Na Citrate stock into 97 mL DI water.

- Add 250 µL C-Tech 2 into the buffer solution. *(Prepares enough for 10 samples)*
- Make up to 100 mL volume by slowly adding DI water.

6. Prepare the enzyme solution 2 $\left(0.25\ \frac{mg\ enzyme}{mL\ solution}\right)$, for adding 50 mg/g-glucan enzyme loading to 50 mg substrate samples, in a 250 mL beaker (label orange).
   - Add 2000 µL Na Citrate stock into 97 mL DI water.
   - Add 625 µL C-Tech 2 into the buffer solution. *(Prepares enough for 10 samples)*
   - Make up to 100 mL volume by slowly adding DI water.

7. Prepare the enzyme solution $\left(1.00\ \frac{mg\ enzyme}{mL\ solution}\right)$, for adding 50 mg/g-glucan enzyme loading to 200 mg substrate samples, in a 250 mL beaker (label blue).
   - Add 2000 µL Na Citrate stock into 95 mL DI water.
   - Add 2500 µL C-Tech 2 into the buffer solution. *(Prepares enough for 10 samples)*
   - Make up to 100 mL volume by slowly adding DI water.

8. Place 15 mL falcon tubes containing cellulosic substrate into an adequately sized rack for easy transportation and utilization.
9. Transfer dilute enzyme solutions into color-coded glass bottles or beakers for transportation purposes, if necessary.
10. Wipe down work area with 70% ethanol mixture for sterilization.

**Answers to pre-lab questions for Instructor** (*in italics*)**:**

**Question 1**: What are carbohydrates, monosaccharides, & polysaccharides? List as many examples of well-known carbohydrates that you find at home or elsewhere.

*Carbohydrates are represented by the stoichiometric formula (CH2O)n, where n is the number of carbons in the molecule. This formula provides the origin of the term "carbohydrate": carbon ("carbo") + water ("hydrate"). Carbohydrates are often classified into three subtypes: monosaccharides, disaccharides, and polysaccharides. Few examples of common responses expected from high school students include; glucose/dextrose, fructose as monosaccharides; sucrose, maltose, lactose as disaccharides; amylose/amylopectin/starch, cellulose, chitin and glycogen as polysaccharides. Depending on the number of carbons in the sugar; pentoses (five carbons), and hexoses (six carbons); are some of the most common building blocks of monosaccharides found in nature.*

**Question 2**: What are enzymes & how do they work? List any example that you can think of.

*Enzymes are biological molecules (typically 3D-folded proteins with a defined molecular structure) that act as catalysts and can significantly speed up the rate of virtually all known biochemical reactions that take place within or outside living cells. Enzymes act on molecules called substrates and converts the substrates into different molecules called products. Like all catalysts, enzymes can increase the reaction rate by lowering its activation energy barrier. Some common examples of enzymes include amylases (hydrolyze amylose), lactase (hydrolyze lactose), cellulase (hydrolyze cellulose), lipase (hydrolyze lipid-based glyceride esters), protease (hydrolyze proteins).*

**Question 3**: What is the impact of temperature on the rate of a chemical reaction? Can you briefly outline any theory that explains how temperature impacts chemical reaction rate?

*Enzymatic activity decreases outside its optimal temperature and pH range depending on the microbial origin of the enzyme. Many enzymes are permanently denatured (or unfolded) when exposed to extreme ranges of non-optimal pH or heat subsequently losing their well-defined 3D-structure and catalytic properties. However, within the optimal working temperature range, increasing the reaction temperature increase the rate of the enzymatic reaction as seen for catalysts performing chemical reactions as well. The Arrhenius equation provides a mathematical formula to relate the temperature dependence of reaction rates. Therefore, enzyme activity initially increases with temperature until the enzyme's structure unfolds fully at non-optimal temperatures.*

**Question 4**: List as many examples of biofuels or lignocellulose biorefining processes used to produce biofuels or biochemicals in industries that you are familiar.

*Bio-gas (by anaerobic digestion of organic biomass), syngas (by water-gas shift reaction), ethanol (by sucrose, starch, and/or cellulosic biomass hydrolysis/fermentation), biodiesel (by transesterification of triglycerides with short-chain alcohols)*

**Question 5**: Do you know what chemical and biochemical engineers do at work? List examples of job profiles of chemical and biochemical engineers.

*A chemical engineer is a professional, who is equipped with the knowledge of chemical engineering to convert basic raw materials into a variety of products, and deals with the design and operation of plants and equipment. Biochemical engineers combine principles of chemical engineering along with biological engineering to focus on bioproducts. The role of chemical and biochemical engineers is to ultimately take the scientific findings made by chemists and biologists in a laboratory and translate those findings into a large-scale manufacturing process or commercially-viable product.*

**Answers to post-lab questions for Instructor** (*in italics*)**:**

**Question 1**: Clearly list the reaction conditions (e.g., enzyme loading, i.e. $\frac{mg\ Enzyme}{g\ Cellulose}$, cellulose or cellobiose substrate concentration, and reaction temperature) that you were assigned for the enzymatic hydrolysis experiment.

*Students should note down the exact reaction conditions used to setup their enzymatic assays to allow them to draw conclusions systematically as a group during post-activity data analysis. Here, the instructor must also ensure that students are provided detailed information about the concentration of enzymes and substrates assigned to each student and/or group ahead of time. Also, ask students to calculate the absolute amount of enzyme added on a 100-gram cellulose basis to help them get a realistic sense of the relative mass of enzymes/proteins needed in a commercial scale biorefinery and hence the economic challenges associated with cellulosic biofuels production. For example, a 50 mg enzyme per gram glucan loading corresponds to needing 5 grams of enzyme for 100 grams cellulose (or 5% w/w basis). Often catalysts are either recycled in a commercial process or is used at much lower absolute mass loadings. e.g., 0.1% w/w basis or lower amylase enzyme loadings are currently used for cost-effectively converting corn grain into fermentable sugars for ethanol production. The students should realize that we need to reduce enzyme usage by at-least 10-100 fold to make the cellulose hydrolysis process as efficient as the commercially-mature corn grain amylase conversion process.*

**Question 2**: List if your sample in the tube dissolved immediately upon addition of the enzyme/buffer solution or was it insoluble? Did you notice any change in the insoluble sample size, morphology, and/or total reaction volume before and after enzymatic hydrolysis for 90 mins? Try to explain your findings.

*Crystalline and amorphous cellulose are completely insoluble in water. Only cellobiose is expected to dissolve completely in water at the given concentrations. Students should also observe that the higher enzyme loadings and higher reaction temperature based samples should show the largest change in sample size/volume (compared to before the hydrolysis reaction was started). Students will notice that amorphous cellulose samples should show a more significant change in sample volume compared to crystalline cellulose. Also, the change in sample size/morphology will be likely correlated with the final glucometer reading (i.e., residual cellulose volume remaining after hydrolysis should be inversely proportional to the amount of glucose released into solution). No changes will be visible during cellobiose hydrolysis since this reaction is fully homogeneous. Students may notice slight discrepancy in the low solids versus high solids concentration assays in terms of total reaction volume. This difference largely arises from the fact that cellulose has a bulk density that differs greatly from water and is also not completely dissolved in solution. This discrepancy makes it technically challenging to accurately calculate %cellulose-to-glucose conversion for the high solids concentration experiments, as highlighted elsewhere.*[19,20] *However, for sake of simplicity, we ignore solids volume related effects at high solids when calculating % conversion for this activity.*

**Question 3**: Would increasing reaction temperature cause enzymes to work faster or slower? Is there an upper limit for the maximum temperature possible to be used? Why?

*Within the optimal working temperature range (25-50ºC for most Trichoderma or Aspergillus derived cellulases/cellobiases), increasing the reaction temperature will increase the rate of the enzymatic reaction to result in higher product yield. The Arrhenius equation provides a mathematical formula to relate the temperature dependence of reaction rates. However, enzyme activity would only increase with temperature until the enzyme's structure unfolds fully at non-optimal temperatures. Most mesophilic sourced cellulolytic enzymes cannot remain properly folded at temperatures exceeding 45-55ºC since these enzymes have co-evolved with the microorganisms that cannot live in comparable temperature range based environments. However, cellulase or cellobiose enzymes isolated from thermophilic organisms living in hot springs can handle much higher temperatures between 55-75ºC (e.g., Acidothermus cellulolyticus, Clostridium thermocellum).*[21–23] *The rates of enzymatic reactions tend to double as the temperature is increased by 10°C.*[24] *Therefore, we should have* *expected the rate of product formation should have ideally increased by >4-fold upon increasing the temperature from 25 to ~50°C. However, as seen in Fig 3 (and SI-VII supporting data), this is clearly not the case for both crystalline or amorphous cellulose upon digestion by cellulases. The actual fold increase seen in overall glucose produced ranged between 1.1-2.5 fold depending on the actual enzyme and substrate loadings. However, in light of the fact that multiple families of cellulolytic enzymes synergistically catalyze cellulose-to-glucose conversion, mass transfer related limitations for insoluble substrates hydrolysis (in the absence of any mixing during hydrolysis), and competing protein denaturation related effects relevant at higher temperatures (like 50 °C), it is not entirely surprising that we didn't see the maximum fold increase in catalytic activity. Note that a ~3-3.5-fold increase in catalytic activity has been reported for Novo188 cellobiase active on cellobiose over a 20 °C increase in temperature (from 40 to 60 °C) and hence would be better experimental system to instruct students on this general topic.*[25]

**Question 4**: In a biorefinery, $Cellulose \xrightarrow{Enzyme} Glucose \xrightarrow{Yeast} Ethanol.$ Will increasing cellulose concentration during hydrolysis make it cheaper to produce bioethanol? Why?

*Most native yeast and engineered microbes can ferment sugars to produce ethanol (e.g., think alcoholic beer versus wine based beverages) at concentrations ranging between 5-15% (w/v). Most microbial fermentations can proceed to completion at lower sugar concentrations, however, the microbial fermentation processes start to slow down at higher concentrations of ethanol due to a myriad number of reasons (e.g., ethanol can inhibit microbial growth at very high concentrations). There is a practical bottleneck of cellulolytic enzymes also getting inhibited by high concentrations of glucose product in the hydrolysate and slightly lower %cellulose-to-glucose conversion at high solids concentration.*[19,20] *Nevertheless, in general, higher cellulose concentrations will result in higher glucose concentrations for comparable hydrolysis yields, and therefore result in higher ethanol concentrations at the end of the fermentation process. Since one of the most energy-intensive unit operation needed in biorefinery is distillation (along with membrane pervaporation or azeotropic distillation) to produce ~100% anhydrous ethanol starting from a dilute 5-15% w/v ethanol broth, higher starting ethanol concentrations reduce the energy required to separate ethanol from water (hence reducing overall cost).*[26,27]

**Question 5**:
(A) Based on the calibration curve data provided below for TrueBalance test strips on actual glucose standards of known concentrations and the results displayed on your glucometer display for your unknown concentration samples, can you estimate the true glucose concentration in your sample using either the fitted equation or graphically?
(B) Knowing the actual glucose concentration in your recovered enzymatic hydrolysate sample, can you estimate what percent of the initial cellulose (or cellobiose) substrate was converted into glucose? Show your work below.

*For part (A), please note that students can simply use the calibration standard based curve to graphically correlate the glucometer reading to the actual glucose concentration (see arrows in red depicting how to use y=200 to roughly estimate x=125 mg/dL). Advanced students who are familiar with the regression method and the regression equation provided, can calculate x=(y+52.167)/1.9735. For example, as shown here, for y=200, x= 127.8 mg/dL.*

*The regression equation method gives an exact solution to the student. Instructor should generate data for known glucose concentrations (50-200 mg/dL) using fresh TRUEBalance test strips to confirm if the calibration curve provided to students for analysis is indeed reliable.*

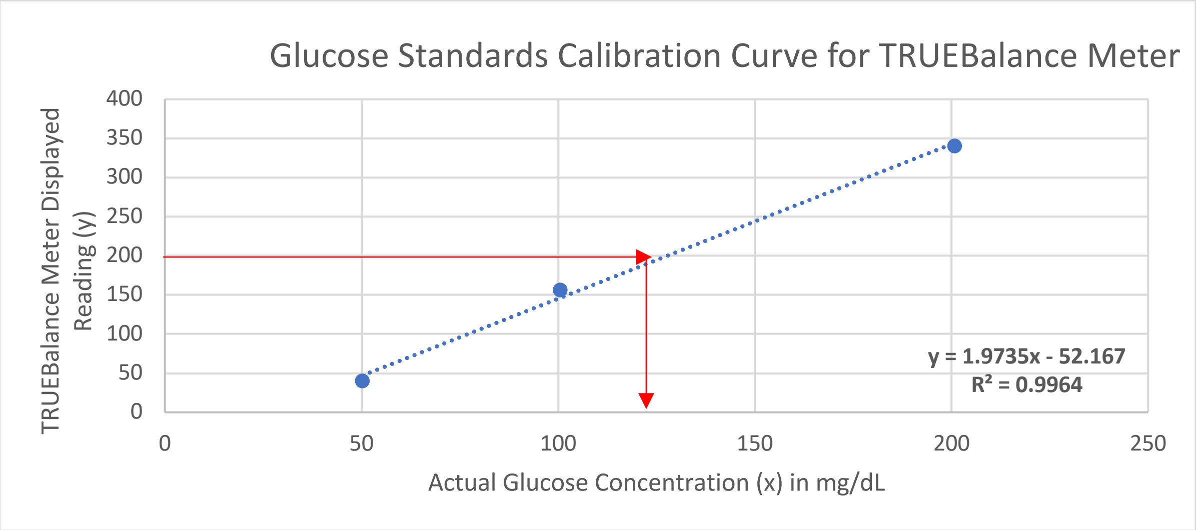

*For part (B), we need to use the reaction stoichiometry to firstly estimate how many moles (or equivalent mass) of glucose would be produced for every mole (or equivalent mass) of cellulose. Here, we have provided the reaction stoichiometry for cellulose similar to what was provided to the students for the simplistic model cellulose like cellobiose (where n=1). Based on this equation it is clear that 1 mole of cellulose (or 2n moles of anhydroglucosyl eq.) would produce 2n moles of glucose, or 1 mole of moles of anhydroglucosyl unit would produce 1 mole of glucose. Therefore, 162 g/L of cellulose polymer (as anhydroglucosyl eq.) would produce 180 g/L glucose on this reaction stoichiometry. Now, based on the actual concentration of cellulose in the tube for students, they can calculate the maximum glucose expected to be released upon completion of hydrolysis reaction. For example, 50 mg cellulose (dry weight basis) in 10 mL total volume is equivalent to a cellulose polymer concentration of 5 mg/ml and would translate to (180/162)*5 or 5.56 g/L of final glucose concentration. If for this starting concentration, one student measured the actual glucose concentration in their reaction tube to be say 278 mg/dL (or equivalent to 2.78 mg/ml or g/L), then the % cellulose-to-glucose yield is simply %100*(2.78/5.56) or 50%. Instructors should note that a correction factor of 180/162 or 1.11 can be used to convert cellulose or glucan to glucose mass units and this method is valid when n is large (n>3). However, while this correction factor can be used for smaller cellulose polymers as well ($1 \leq n \leq 3$), like cellobiose, it would result in slightly erroneous under-prediction of actual yield (by ~max 4.5%). Students can check this by using the exact cellobiose hydrolysis stoichiometric equation to estimate the true correction factor of 1.05 versus the over-predicted generic factor of 1.11.*

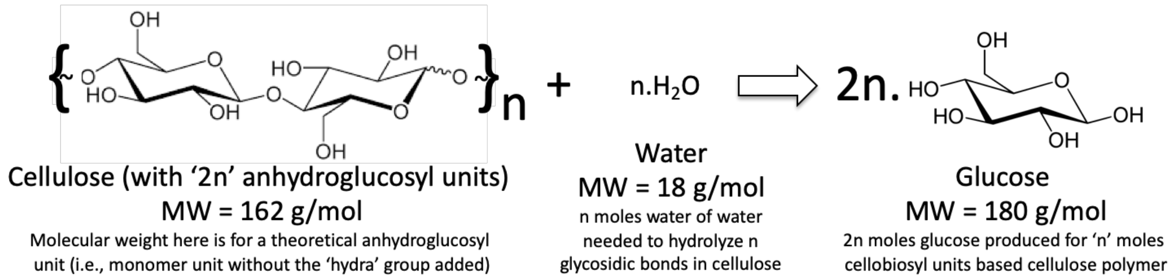

***Bonus Post-Activity Discussion and Advanced Analysis for Advanced Students Only.** *Please note that this bonus question could be used to analyze actual cellulose vs. cellobiose hydrolysis kinetics dataset generated either ahead of time by the instructor to compare cellulase vs. cellobiase reaction kinetics, respectively, or be part of an extended student data analysis activity similar to previously published enzyme kinetics protocols and data analysis for soluble lactose hydrolysis by lactase enzymes.*[28] *Instructors can refer to previously published Michaelis-Menten (MM) parameters available for commercial-grade Novo188 cellobiase enzyme cocktail if they use crude enzyme cocktail to measure MM model parameters for crude Novo188 enzyme cocktail using cellobiose as substrate.*[29] *Instructors should be also aware of the major limitations of using traditional MM type models to characterize insoluble substrate (cellulose) hydrolysis by a synergistic cocktail of multiple classes of cellulase enzymes, unlike soluble cellobiose hydrolysis by cellobiase.*[30]

**Background:** To understand and model how enzymes function, Michaelis-Menten (MM) based models were developed first in 1913 to mathematically characterize the relationship between the rate of product formation and starting substrate concentration to estimate $k$ and $K_m$ parameters specific to each enzyme (see Figure 1 for overview to MM method).[31,32] Details about the derivation of the MM model and curve-fitting methods used to estimate the model parameters are elsewhere in the literature.[31,32] Now, based on your experimental findings from the previous activity, you need to compare the relative rate of hydrolysis of insoluble versus soluble cellulosic substrates and draw some conclusions about the relative hydrolytic activity of cellulases versus cellobiases, respectively.

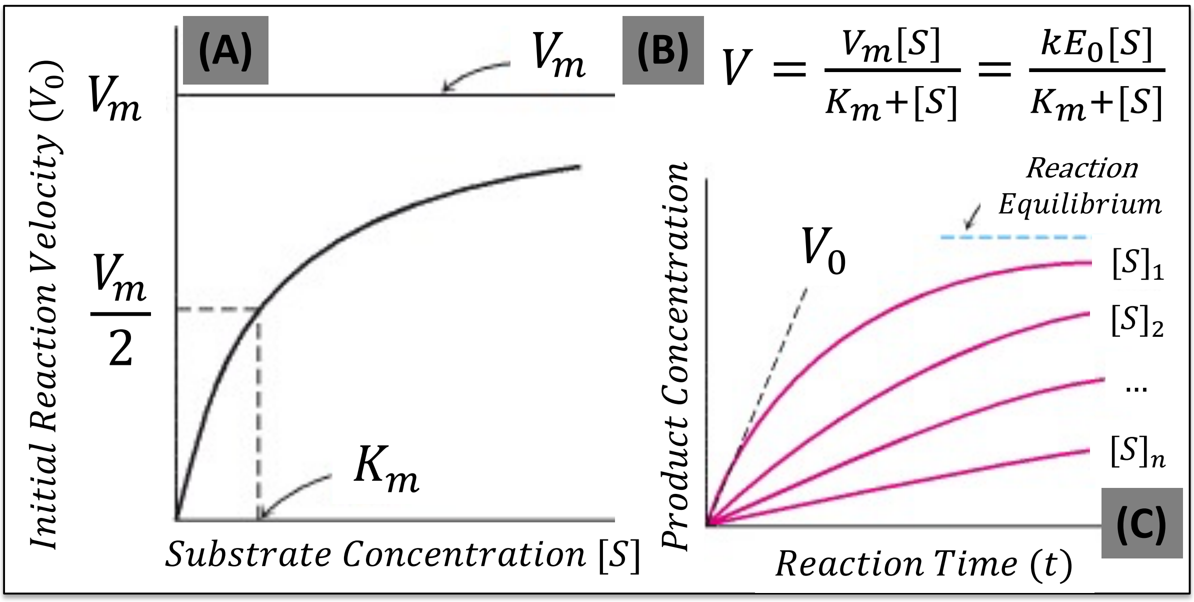

**Figure 1. Overview to Michaelis-Menten Enzyme Kinetics Model. (A) Here, a plot of the reaction velocity as a function of the substrate concentration for an enzyme that obeys MM kinetics is shown. (B) Michaelis-Menten Enzyme Kinetics Model used to plot curve in (A) is shown here. The MM equation is used to model the rate of product formation ($V$) as a function of initial substrate concentration as $V = \frac{V_m[S]}{K_m+[S]}$, where $V_m = k[E_0]$. Here, $[E_0]$ is the concentration of added initial enzyme, $[S]$ is the concentration of the added initial substrate, $K_m$ is equivalent to the substrate concentration that yields half the maximum reaction velocity for the enzyme ($0.5V_m$), and $k$ is the reaction rate or turnover constant for the enzyme. (C) The amount of product formed at different substrate concentrations is plotted as a function of enzyme reaction time. The initial velocity ($V_0$) for each substrate concentration can be determined from the slope of the curve at the beginning of each reaction as shown here for $[S]_1$. Here reaction equilibrium is the same as enzymatic reaction going to completion (e.g., 100% cellobiose-to-glucose conversion by cellobiase enzyme at any given starting substrate concentration).**

***Bonus Question 6:** In order to plot the MM plot shown here for either cellulose or cellobiose hydrolysis to glucose and determine the relevant MM model parameters by curve fitting, researchers must first conduct detailed enzymatic assays to determine the rate of product formation as a function of reaction time for a series of substrate concentrations. As part of a preliminary investigation, you have been provided with enzyme-substrate reaction kinetics data for both cellulose and cellobiose substrates conversion into product glucose as a function of reaction time for relevant crude enzyme cocktails (see Figure 2 below). Before performing additional assays at varying starting substrate concentrations to develop a comprehensive MM type model to characterize cellulase and cellobiase activities, compare the initial rate of cellulose-to-glucose and cellobiose-to-glucose conversion using the results shown in Figure 2.

a) Based on this initial velocity activity data, can you comment if cellulose is more or less efficiently hydrolyzed by cellulases compared to cellobiose hydrolysis by cellobiase enzymes into glucose? What is the relative catalytic rate for cellulose hydrolysis compared to cellobiose hydrolysis by corresponding enzymes?

*Firstly, instructors should note that this example dataset was generated based on a single substrate concentration-based hydrolysis assay conducted along with multiple time point samples removed for glucose assay as discussed in SI-VII. Instructor can generate kinetic datasets for their available cellulase-cellulose and cellobiase-cellobiose systems for a range of initial substrate concentrations (and fixed enzyme loadings). The enzyme loadings can be pre-decided such that %conversion of substrate to glucose doesn't exceed 5-10% (i.e., the rate of initial glucose release per unit time follows a highly linear relationship as shown in the figure below), to accurately estimate the initial reaction velocity where most conditions of the MM model are satisfied. For the current bonus question, use the data generated and provided by the authors for illustrative purposes.*

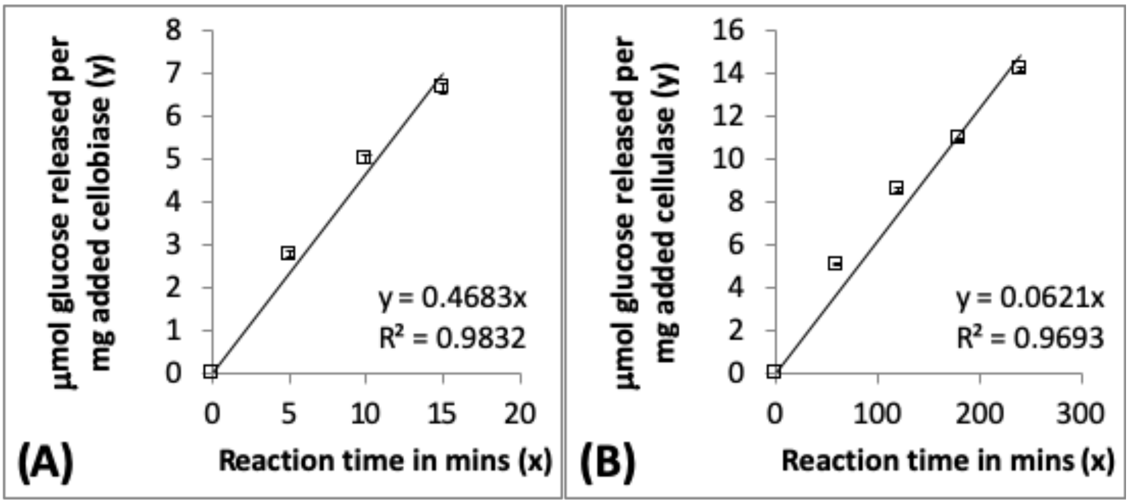

**Figure 2. Initial reaction velocity ($V_0$) with units of µmoles glucose released per mg crude enzyme cocktail added for cellobiase-cellobiose (A) and cellulase-cellulose (B) systems are shown here. Here, Novo188 and Celluclast commercial enzymes were used for cellobiase and cellulase cocktails, respectively. Amorphous cellulose was used here instead of microcrystalline cellulose. The substrate concentration was fixed at ~10 g/L, enzyme concentration was fixed at ~1 mg enzyme/g substrate, and the reaction was conducted without shaking at 50 C. Glucose released was monitored using TRUEBalance type glucose assay method as highlighted in the activity protocol. Fitted line and associated fit-equation for the data based on a linear regression analysis using Excel is shown here.**

*Here, students can use an analytical, or graphical approach, to estimate the rate of glucose product formation for each substrate. Using the analytical approach, since the slopes of initial enzymatic reactions are provided here, we can easily determine the initial velocity profile for the cellulase-cellulose and cellobiase-cellobiose systems to be 0.0621 and 0.4683 moles glucose released per min, respectively. Based on this initial velocity values it is clear that cellobiose is hydrolyzed by cellobiose enzymes at roughly 7.5-fold higher rates than cellulose hydrolysis by cellulase enzymes. This means that cellulose is less efficiently hydrolyzed and would require a greater amount of cellulase enzyme to achieve comparable hydrolysis rates and product yields. Instructors should note that here crude enzyme cocktails were used to estimate the initial velocity data but similar data can be generated for purified single enzymes as well on either substrate, as often described in the literature.*[2,14,33–35]

b) Can you hypothesize why one form of cellulosic substrate is more readily hydrolyzed into glucose by the relevant enzymes?

*There are multiple reasons why highly soluble substrates like cellobiose and hydrolyzed more readily than insoluble cellulose and is an topic of active research within the scientific community.*[36,37] *Briefly, the rate of enzymatic hydrolysis of cellulosic substrates is closely dependent on two broad class of factors; substrate-dependent factors and enzyme-dependent factors.*[37–39] *Substrate-dependent factors like substrate solubility, glycosidic bond accessibility, cellulose polymer chain length, cellulose crystallinity etc all impact enzyme kinetics. Similarly, enzyme-dependent factors based on the processive/non-processive nature of active site catalytic mechanism, multidomain structure dynamics, inhibition due to non-productive binding interactions etc also impact enzyme kinetics. While it is challenging to clearly draw conclusions when comparing activity of two different classes of enzymes (e.g., Cellulases from Trichoderma vs. Cellobiases from Aspergillus), one can clearly draw some conclusions when testing the activity of the same enzyme family on different forms of cellulose. As shown earlier in the main text, amorphous cellulose generated from crystalline cellulose clearly showed high rates of hydrolysis by the same class of cellulase enzymes. Similarly, removal of lignin also helps improve enzyme accessibility to cellulose/hemicellulose polymers and prevents non-productive enzyme binding to result in higher specific activity as shown in the supplementary information section of the acid-chlorite pretreated corn stover samples. Instructor should highlight that pretreatment is therefore an important unit operation in biorefinery that helps reduce recalcitrance of complex cellulosic biomass to hydrolysis by cellulases (and other carbohydrate-active enzymes) to produce fermentable sugars like glucose (and xylose etc).*

#### (SI-XI) References for Supporting Information Document

1. Miller, G. L. Use of dinitrosalicylic acid reagent for determination of reducing sugar. *Anal Chem* **31**, 426–428 (1959).
2. Gao, D., Chundawat, S. P. S., Krishnan, C., Balan, V. & Dale, B. E. Mixture optimization of six core glycosyl hydrolases for maximizing saccharification of ammonia fiber expansion (AFEX) pretreated corn stover. *Bioresour. Technol.* **101**, 2770–2781 (2010).
3. Chundawat, S. P. S., Balan, V. & Dale, B. E. High-throughput microplate technique for enzymatic hydrolysis of lignocellulosic biomass. *Biotechnol. Bioeng.* **99**, 1281–1294 (2008).
4. Dangkulwanich, M., Kongnithigarn, K. & Aurnoppakhun, N. Colorimetric Measurements of Amylase Activity: Improved Accuracy and Efficiency with a Smartphone. *J. Chem. Educ.* **95**, 141–145 (2018).
5. Ucar, G. & Balaban, M. Hydrolysis of Polysaccharides with 77% Sulfuric Acid for Quantitative Saccharification. *Turkish J. Agric. For.* **27**, 361–365 (2003).
6. Sluiter, J. B., Ruiz, R. O., Scarlata, C. J., Sluiter, A. D. & Templeton, D. W. Compositional Analysis of Lignocellulosic Feedstocks. 1. Review and Description of Methods. *J. Agric. Food Chem.* **58**, 9043–9053 (2010).
7. Ishizawa, C. *et al.* Can delignification decrease cellulose digestibility in acid pretreated corn stover? *Cellulose* **16**, 677–686 (2009).
8. Haarmeyer, C. N., Smith, M. D., Chundawat, S. P. S., Sammond, D. & Whitehead, T. A. Insights into cellulase-lignin non-specific binding revealed by computational redesign of the surface of green fluorescent protein. *Biotechnol. Bioeng.* **114**, 740–750 (2017).
9. Wise, L. E., Murphy, M. & D Adieco, A. A. Chlorite holocellulose, its fractionation and bearing on summative wood analysis and studies on the hemicelluloses. *Paper Trade Journal* **122**, 35–43 (1946).
10. Kumar, R., Hu, F., Hubbell, C. A., Ragauskas, A. J. & Wyman, C. E. Comparison of laboratory delignification methods, their selectivity, and impacts on physiochemical characteristics of cellulosic biomass. *Bioresour. Technol.* **130**, 372–381 (2013).
11. Walseth, C. S. Occurrence of cellulase in enzyme preparations from microorganisms. *Tappi* **35**, 228–233 (1952).
12. Zhang, Y. H. P., Cui, J. B., Lynd, L. R. & Kuang, L. R. A transition from cellulose swelling to cellulose dissolution by o-phosphoric acid: Evidence from enzymatic hydrolysis and supramolecular structure. *Biomacromolecules* **7**, 644–648 (2006).
13. Chundawat, S. P. S. & Agarwal, U. P. Swelling by Hydrochloric Acid Partially Retains Cellulose-I Type Allomorphic Ultrastructure But Enhances Susceptibility toward Cellulase Hydrolysis Such as Highly Amorphous Cellulose. in *Understanding Lignocellulose: Synergistic Computational and Analytic Methods (ACS Symposium Series Vol. 1338)* 69–88 (ACS Symposium Series Vol. 1338, 2019). doi:10.1021/bk-2019-1338.ch005
14. Chundawat, S. P. S. *et al.* Proteomics based compositional analysis of complex cellulase-hemicellulase mixtures. *J. Proteome Res.* **10**, 4365–4372 (2011).
15. Cantarel, B. L. *et al.* The Carbohydrate-Active EnZymes database (CAZy): an expert resource for Glycogenomics. *Nucl. Acids Res.* **37**, D233-238 (2009).
16. Gao, D. *et al.* Hemicellulases and auxiliary enzymes for improved conversion of lignocellulosic biomass to monosaccharides. *Biotechnol. Biofuels* **4**, 5 (2011).
17. Wolfenden, R. *et al.* Spontaneous Hydrolysis of Glycosides. *JACS* **7863**, 6814–6815

(1998).
18. Wolfenden*, R. Degrees of Difficulty of Water-Consuming Reactions in the Absence of Enzymes. (2006). doi:10.1021/CR050311Y
19. Weiss, N. D., Felby, C. & Thygesen, L. G. Enzymatic hydrolysis is limited by biomass–water interactions at high-solids: improved performance through substrate modifications. *Biotechnol. Biofuels* **12**, 3 (2019).
20. Kristensen, J. B., Felby, C. & Jørgensen, H. Determining Yields in High Solids Enzymatic Hydrolysis of Biomass. *Appl. Biochem. Biotechnol.* **156**, 127–132 (2009).
21. Heinzelman, P. *et al.* A family of thermostable fungal cellulases created by structure-guided recombination. *Proc. Natl. Acad. Sci.* **106**, 5610–5615 (2009).
22. Brunecky, R. *et al.* Revealing nature's cellulase diversity: the digestion mechanism of Caldicellulosiruptor bescii CelA. *Science* **342**, 1513–6 (2013).
23. Tucker, M. P., Mohagheghi, A., Grohmann, K. & Himmel, M. E. Ultra-Thermostable Cellulases From Acidothermus cellulolyticus: Comparison of Temperature Optima with Previously Reported Cellulases. *Nat Biotech* **7**, 817–820 (1989).
24. Laidler, K. J. & Peterman, B. F. [10] Temperature effects in enzyme kinetics. *Methods Enzymol.* **63**, 234–257 (1979).
25. Bravo, V., Paez, M. P., Aoulad, M. & Reyes, A. The influence of temperature upon the hydrolysis of cellobiose by β-1,4-glucosidases from Aspergillus niger. *Enzyme Microb. Technol.* **26**, 614–620 (2000).
26. Klein-Marcuschamer, D., Oleskowicz-Popiel, P., Simmons, B. A. & Blanch, H. W. Technoeconomic analysis of biofuels: A wiki-based platform for lignocellulosic biorefineries. *Biomass and Bioenergy* **34**, 1914–1921 (2010).
27. Tao, L. *et al.* Process and technoeconomic analysis of leading pretreatment technologies for lignocellulosic ethanol production using switchgrass. *Bioresour. Technol.* **102**, 11105–14 (2011).
28. Heinzerling, P., Schrader, F. & Schanze, S. Measurement of Enzyme Kinetics by Use of a Blood Glucometer: Hydrolysis of Sucrose and Lactose. *J. Chem. Educ.* **89**, 1582–1586 (2012).
29. Dekker, R. F. H. Kinetic, inhibition, and stability properties of a commercial β-D-glucosidase (cellobiase) preparation from aspergillusniger and its suitability in the hydrolysis of lignocellulose. *Biotechnol. Bioeng.* **28**, 1438–1442 (1986).
30. Kari, J., Andersen, M., Borch, K. & Westh, P. An Inverse Michaelis–Menten Approach for Interfacial Enzyme Kinetics. *ACS Catal.* **7**, 4904–4914 (2017).
31. Berg, J. M., Tymoczko, J. L. & Stryer, L. *The Michaelis-Menten Model Accounts for the Kinetic Properties of Many Enzymes*. (W H Freeman, 2002).
32. Johnson, K. A. & Goody, R. S. The Original Michaelis Constant: Translation of the 1913 Michaelis–Menten Paper. *Biochemistry* **50**, 8264–8269 (2011).
33. Kabel, M. A., van der Maarel, M. J. E. C., Klip, G., Voragen, A. G. J. & Schols, H. A. Standard assays do not predict the efficiency of commercial cellulase preparations towards plant materials. *Biotechnol. Bioeng.* **93**, 56–63 (2006).
34. Sharrock, K. R. Cellulase assay methods: a review. *J. Biochem. Biophys. Methods* **17**, 81–105 (1988).
35. Zhang, Y. H. P., Hong, J. & Ye, X. Cellulase Assays. in *Biofuels: Methods and Protocols (Methods in Molecular Biology)* (ed. Mielenz, J. R.) **581**, 213–231 (Humana Press, 2009).
36. Chundawat, S. P. S., Beckham, G. T., Himmel, M. & Dale, B. E. Deconstruction of

Lignocellulosic Biomass to Fuels and Chemicals. *Annu. Rev. Chem. Biomol. Eng.* **2**, 121–145 (2011).

37. Payne, C. M. *et al.* Fungal Cellulases. *Chem. Rev.* **115**, 1308–1448 (2015).
38. Bansal, P., Hall, M., Realff, M. J., Lee, J. H. & Bommarius, A. S. Modeling cellulase kinetics on lignocellulosic substrates. *Biotechnol. Adv.* **27**, 833–848 (2009).
39. Gan, Q., Allen, S. J. & Taylor, G. Kinetic dynamics in heterogeneous enzymatic hydrolysis of cellulose: an overview, an experimental study and mathematical modelling. *Process Biochem.* **38**, 1003–1018 (2003).
